# *RNF213* variation, a broader role in neurovascular disease in Caucasian and Japanese populations

**DOI:** 10.1101/2020.03.31.013078

**Authors:** Oswaldo Lorenzo-Betancor, Patrick R. Blackburn, Luca Farrugia, Alexandra I. Soto-Beasley, Ronald L. Walton, Emily Edwards, Rabih G. Tawk, Eric W. Klee, William D. Freeman, David Miller, James Meschia, Owen A. Ross

## Abstract

Moyamoya disease (MMD) is a chronic, occlusive cerebrovascular disease that predominantly affects East Asian populations. The major genetic mutation associated with MMD in Asian populations is the p.R4810K substitution in Ring Finger Protein 213 (RNF213). Interestingly, variants in the *RNF213* gene have also been implicated in intracranial aneurysms (IA) in French-Canadian population, suggesting that variation in this gene may play a broader role in cerebrovascular phenotypes. In a recent genome-wide association study (GWAS) in a Caucasian population, variants rs6565653 and rs12601526 in the *Solute Carrier Family 26 Member 11* (*SLC26A11*) gene, which is less than 10kb away from *RNF213*, showed a suggestive association with young onset ischemic stroke. We propose that the signal could be tagging an association with common variation in the *RNF213* gene. We analyzed the linkage disequilibrium (LD) pattern in the *SLC26A11-RNF213* gene region and we observed a high LD between variants in this region based on D’ values. We show that *SLC26A11* rs6565653 variant tags *RNF213* rs12944088, a missense variant that is more common among subjects with IA than in healthy individuals. Given the fact that rs6565653 tags several *RNF213* variants, it is highly likely that some of these tagged variants modify the risk of suffering stroke. The LD analyses suggest that the *SLC26A11* signal from the young onset ischemic stroke GWAS performed in a Caucasian population is also tagging variation at the *RNF213* loci, supporting the hypothesis that *RNF213* variation may result in a variety of neurovascular disorders including an increased risk and/or worse prognosis following ischemic stroke in Caucasian population.

## Introduction

Moyamoya disease (MMD) is a chronic, occlusive cerebrovascular disease characterized by bilateral steno-occlusive changes that affect mainly the terminal portion of the internal carotid artery and the presence of abnormal vascular networks in the basal ganglia (Suzuki and Takaku 1969). MMD is most common in East Asian populations and its incidence ranges from 0.35 to 2.3 cases per 100,000 Japanese subjects (Wakai et al. 1997; Ahn et al. 2014) compared to 0.086 cases per 100,000 people in the US population (Uchino et al. 2005). MMD can be divided into early onset and late onset forms with a biphasic peak pattern in incidence that occurs in the first and fourth decades of life (Ahn et al. 2014).

While the etiology of MMD remains unknown, up to 10-15% of cases are familial and are suggestive of an autosomal dominant pattern of inheritance with incomplete penetrance. A major genetic determinant associated with MMD in the Asian population is the p.R4810K mutation in the *Ring Finger Protein 213* (*RNF213*) which is present in approximately 90% of all Japanese, 79% of Korean and 23% of Chinese MMD patients (Liu et al. 2011). Interestingly, this mutation is also present in 2.3% of the East Asian healthy controls, which highlights the reduced penetrance of the mutation and suggests there are likely strong modifiers of disease (Liu et al. 2011). It is also worth noting that the p.R4810K mutation has never been reported in the European/Caucasian population, in either MMD cases or controls, although other rare *RNF213* variants have been reported (Kobayashi et al. 2016).

The p.R4810K mutation has also been described to increase the risk of ischemic stroke attributable to large-artery atherosclerosis in East Asian population (Okazaki et al. 2019). Interestingly, in a genome-wide association study (GWAS) in young onset ischemic stroke performed in a Caucasian population (Cheng et al. 2016), two single nucleotide polymorphisms (SNPs) in the *Solute Carrier Family 26 Member 11* (*SLC26A11*) gene (rs6565653 and rs12601526), reached suggestive *p*-values of 5.51×10^-7^ and 1.14×10^-6^, respectively (figure S1). Notably, the *SLC26A11* gene is less than 10kb away from the *RNF213* gene. Therefore, it is possible that the signal in the *SLC26A11* gene from this study may actually be tagging an association with common variation in the *RNF213* gene.

To examine this hypothesis we investigated the linkage disequilibrium (LD) patterns for common variation in the 17q25.3 chromosomal region that includes the *SLC26A11* and the *RNF213* genes in two ethnically different populations from the 1000 Genomes project. Additionally, we analyzed the expression changes in both genes according to their genetic variation in arterial tissues using the Genotype-Tissue Expression (GTEx) project data.

## Materials and methods

### *RNF213* and *SLC26A11* variation analysis

The linkage disequilibrium (LD) pattern between the *SLC26A11* and the *RNF213* gene SNPs was analyzed using common variants (minor allele frequency [MAF] > 0.01) located in both genes in two different publicly available populations: 99 unrelated Utah Residents with Northern and Western European ancestry (CEU population) and 104 unrelated Tokyo subjects with Japanese ancestry (JPT population) from the 1000 Genomes Project (http://www.internationalgenome.org/). In order to carry out the LD analyses between the *SLC26A11* and the *RNF213* genes, all variants located between chr17:78192200 and chr17:78374581 positions (build GRCh37) were extracted from the phase3 supporting genotype file from 1000 Genomes Project (ftp://ftp.1000genomes.ebi.ac.uk/vol1/ftp/release/20130502/supporting/vcf_with_sample_level_annotation/), and the CEU and JPT variants were filtered according to the European and East Asian MAFs, respectively. Proxy Single Nucleotide Polymorphisms (SNPs) were selected for both sets of variants with SNAP Proxy Search tool (http://www.broadinstitute.org/mpg/snap/) setting a minimum LD threshold of r^2^ > 0.8. The CEU and the JPT + Han Chinese in Beijing (CHB) populations from the Hapmap 3 SNP dataset were used to analyze the CEU and JPT variants, respectively. The subset of tagging variants (92 CEU and 80 JPT SNPs, respectively, tables S1 and S2) was analyzed with the R LDHeatmap v.0.99-2 package (http://stat.sfu.ca/statgen/research/ldheatmap.html) to generate LD plots for this genomic region in both populations (figure 1). Additionally, the LDproxy tool from the LDlink online suite (https://analysistools.nci.nih.gov/LDlink/?tab=home) was used to search for proxy SNPs of rs6565653 and rs12601526 in the CEU and JPT populations (figures S2 and S3).

**Figure 1.**
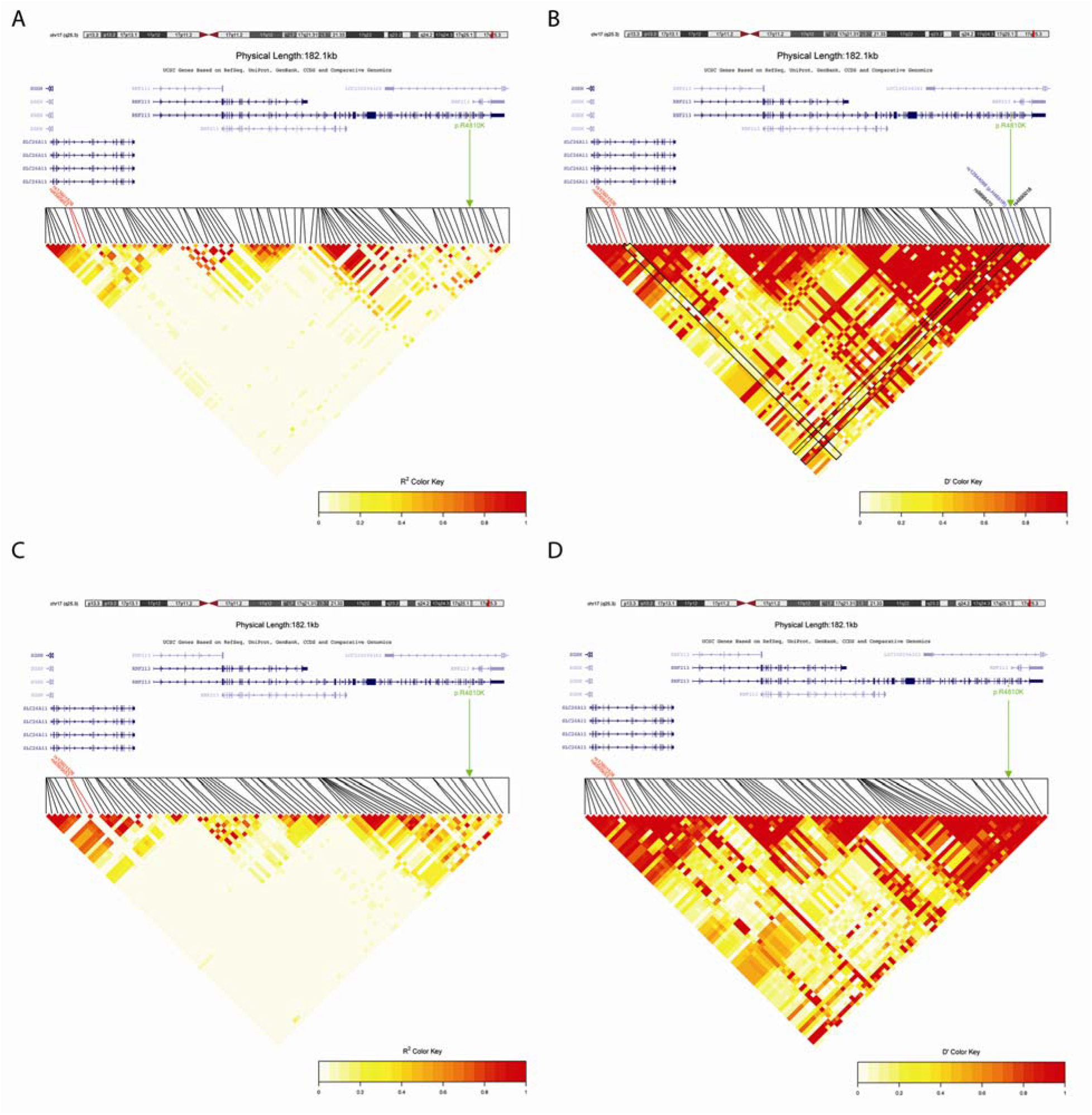
Pairwise LD for common tagging SNPs located in the *SLC26A11* and *RNF213* genes in CEU and JPT populations. (A) R^2^ values in CEU population; (B) D’ values in CEU population; (C) R^2^ values in JPT population; (D) D’ values in JPT population. The two SNPs that were associated with young onset ischemic stroke in Caucasian population located in *SLC26A11* gene are highlighted in red. The Moyamoya disease p.R4810K mutation located in *RNF213* exon 60 is highlighted in green. The missense p.H4691R variant in LD with rs6565653 is highlighted in blue. Black polygons in (B) highlight D’ LD scores between rs6565653, rs12601526 and the SNPs around the RNF213 p.R4810K mutation. LD plot was generated with R LDheatmap package.

### SLC26A11 and RNF213 expression changes in arterial tissues

Local expression quantitative trait loci (cis-eQTLs) and expression changes due to the SNPs driving these eQTLs were extracted from available vascular tissues (aorta, coronary and tibial arteries) from the Genotype-Tissue Expression project (GTEx) for *SLC26A11* and *RNF213* genes to determine whether there were any significant expression changes in these tissues. In order to analyze the cis-eQTLs for *SLC26A11* and *RNF213* genes, all SNPs included in an up and downstream flanking region of 25kb from both genes were extracted from the eQTL files from the three available vascular tissues (aorta, coronary artery and tibial artery) from the Genotype-Tissue Expression project (GTEx) Analysis V6p dataset (http://www.gtexportal.org/home/datasets). Several R packages were used to plot all variants with significant eQTL values assuming a false discovery rate (FDR) <5% (figure 2 and figures S4 and S5). Additionally, all the mRNA expression variations associated to these eQTLs were also plotted for both genes (figure 2 and figures S4 and S5).

**Figure 2.**
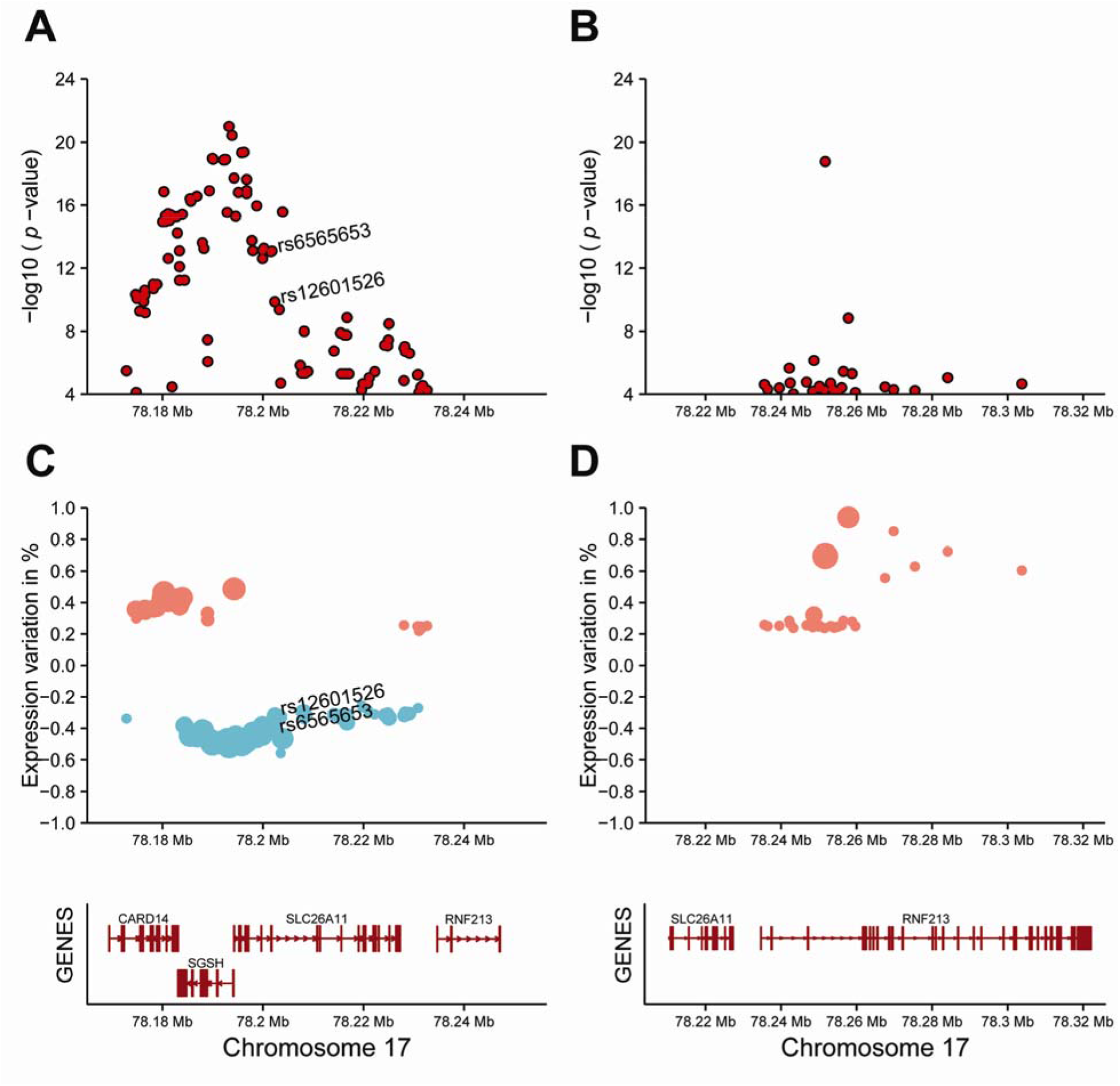
Plots showing eQTL values and expression data of *SLC26A11* and *RNF213* genes in tibial artery tissue. Significant local expression quantitative trait loci (cis-eQTL) values for *SLC26A11* (A) and *RNF213* (B) SNPs, respectively. Red dots in A and B are significant cis-eQTLs (at false discovery rate <5%) for *SLC26A11* and *RNF213* genes in tibial artery tissue. Protein expression changes for *SLC26A11* (C) and *RNF213* (D) mRNA due to SNPs effect in tibial artery. In C and D, red dots represent overexpression and blue dots underexpression, respectively. The size of each dot represents the −log_10_ (*p*-value) between the SNP and the expression change. Figures elaborated using GTEx data. *CARD14* = Caspase Recruitment Domain Family Member 14; *SGSH* = N-Sulfoglucosamine Sulfohydrolase; *SLC26A11* = Solute Carrier Family 26 Member 11; *RNF213* = Ring Finger Protein 213; Mb = megabase.

### Data availability

Supplemental files available at FigShare. Figure S1 includes the genome-wide association results of early-onset ischemic stroke based on a transethnic meta-analysis and a European-only meta-nalysis and the proximity of the two significant *SLC26A11* SNPs to the *RNF213* gene. Figure S2 and S3 show R^2^ and D’ values for proxy SNPs for rs6565653 and rs12601526 *SLCA26A11* variants in CEU and JPT populations, respectively. Figure S4 shows significant local expression quantitative trait loci (cis-eQTL) values for *SLC26A11* and *RNF213* SNPs, affecting arterial tissues. Figure S5 shows expression changes in SLC26A11 and RNF213 due to SNP effects in arterial tissues. Tables S1 and S2 show chromosome, position and allele frequencies for SNPs used in the LD analyses in CEU and JPT populations, respectively. File S1 includes the R code that was used to generate the results of the current analysis and to plot all the figures from the manuscript itself as well as from the supplementary materials.

## Results

### LD patterns in CEU and JPT populations

S1 and S2 Tables show the allele frequencies for the SNPs used in the LD analyses in CEU and JPT populations, respectively. The differences between allele frequencies across SNPs for this region explain the apparent low LD between *SLC26A11* and *RNF213* variants in both populations according to r^2^ values (figures 1A and 1C). However, if we measure the D’ values, which do not take into account allele frequencies, for the same variants in both populations we can see that the LD between variants in *SLC26A11* and *RNF213* is greater than expected (figures 1B and 1D). The lack of genotypes for *RNF213* p.R4810K variant in both populations makes it difficult to establish the potential LD of variants in *SLC26A11* with this specific mutation (figure 1 and figures S2 and S3). However, in CEU population, rs6565653 reaches D’ scores of 0.99 with the missense *RNF213* rs12944088 (p.H4691R) variant and even 1.0 with intronic *RNF213* rs4890018 and rs9898470 SNPs. These three SNPs are located in the vicinity of the pathogenic p.R4810K *RNF213* mutation (figure 1). Additionally, in the JPT population there is a missense variant in the *RNF213* gene (rs142798005, MAF = 0.0096; figure S3) with a D’ value of 1 for both *SLC26A11* SNPs (rs6565653, MAF = 0.024; rs12601526, MAF = 0.0288). However, due to the different MAFs of the three SNPs, the r^2^ values reach only 0.3942 and 0.3269 scores when evaluating rs142798005 as a proxy SNP of rs6565653 and rs12601526, respectively.

### *SLC26A11* and *RNF213* mRNA expression patterns in arterial tissues

Given the fact that rs6565653 tags multiple *RNF213* variants, including the missense p.H4691R change, we cannot rule out that *SLC26A11* variants are tagging *RNF213* variants that modify expression. From the analysis of the eQTL results, it is evident that variation in both *SLC26A11* and *RNF213* genes affects their respective expression. It is interesting though that the eQTLs and the expression variation is greater in the tibial artery than in the aorta or in the coronary arteries (figures S4 and S5). Of note, while most of the variation in the *SLC26A11* gene, including rs6565653 and rs12601526, tends to downregulate its expression, the variation in the *RNF213* gene, shows a trend towards its overexpression, suggesting a possible toxic gain-of-function for *RNF213* mutations (figure 2 and figures S4 and S5).

## Discussion

In this paper we analyzed the *SLC26A11* and *RNF213* LD patterns in CEU and JPT populations in order to assess whether *SLC26A11* variation could be tagging *RNF213* variants and thus would implicate *RNF213* as a potential disease gene in ischemic stroke. We have shown, in CEU population, that *SLC26A11* rs6565653 tags the *RNF213* rs12944088 (p.H4691R) variant. Interestingly, this variant has a MAF of 0.0159 in Caucasian population and was found to be more frequent among subjects with IA (0.0365) than in healthy controls (0.0163) in the French-Canadian IA study (Zhou et al. 2016). However, this result was not observed in the JPT population, because most of the variation in the *RNF213* gene seems to be specific to a given ethnic background. In fact, figure S3 shows a paucity of SNPs with high r^2^ scores in *SLC26A11* and *RNF213* genes region in the JPT population, suggesting that the SNPs in this region have highly variable MAFs and many of them are rare variants specific to the JPT population.

Evidence from several recent studies suggests that *RNF213* is a susceptibility gene for a number of different neurovascular conditions including ischemic and hemorrhagic stroke. Miyawaki et al. also showed an increased risk of intracranial major artery stenosis / occlusion (ICASO) in a selected Japanese population who carry the *RNF213* p.R4810K mutation (Miyawaki et al. 2013). Similarly, Bang et al. found that *RNF213* p.R4810K is associated with an increased risk of intracranial atherosclerotic stenosis (ICAS) in East Asians. Furthermore, this study found that ICAS patients with the common *RNF213* variant were younger than those without the variant and hypothesized that this variant could lead to vascular fragility in a subset of patients, resulting in ischemic and hemorrhagic neurovascular presentations (Bang et al. 2016). Studies performed in the French-Canadian population suggest that variation in *RNF213* is also a risk factor for developing intracranial aneurysms (IA) (Zhou et al. 2016).

Recently, a multiancestry association study that included 520,000 subjects identified 32 genome-wide significant loci, 22 of which were novel, associated with stroke (Malik et al. 2018). However, *RNF213* gene was not one of these novel hits. There are several reasons that can explain the absence of the signal in the multiancestry study. One of them is that the effect of rare *RNF213* mutations, including the p.R4810K variant, is ethnic specific affecting predominantly East Asian population and the number of cases and controls in the multiancestry study for each population is not proportional. In this paper, we showed that *RNF213* variation is different between CEU and JPT populations. In fact, p.R4810K has never been described in European samples and missense variants in this gene are rare among Caucasian population and different from the ones that have been described in Asian populations. To be more specific, the multiancestry genome-wide association study included 17 European studies accounting for 40,585 cases and 406,111 controls, two East Asian studies that included 17,369 cases and 28,195 controls and 3 South Asian studies that included 2,437 cases and 6,707 controls. In total there are 40,585 European cases and 406,111 European controls against the 19,806 Asian cases and 34,902 Asian controls. Therefore, the European population size could be hiding the *RNF213* variation effect.

The original *SLCA26A11* GWAS signal was found in a European early onset ischemic stroke cohort (Cheng et al. 2016). A study performed in 2018 in 70 Japanese patients (20-60 years of age) with intracranial arterial stenosis suffering non-cardioembolic or transient ischemic stroke but without moyamoya disease, examined the prevalence of the *RNF213* p.R4810K variant and identified this variant in 17 out of 70 patients (24.3%) (Kamimura et al. 2019). Additionally, the *RNF213* p.R4810K variant was found in 35% of patients with stenosis in the M1 segment of the middle cerebral artery or the A1 segment of the anterior cerebral artery but in only one patient (9%) with intracranial posterior circulation stenosis. It is possible that the European *SLCA26A11* GWAS signal and that the *RNF213* p.R4810K variant contribute mainly to early onset ischemic stroke. As a result, the signal may not reach significance in the multiancestry GWAS because the population of the later study either did not include enough early onset stroke cases, as the purpose of this GWAS was not to assess risk variants for this specific phenotype.

Interestingly, a meta-analysis of 22 migraine GWA studies, including 59,674 patients and 316,078 controls identified 16 SNPs in the *RNF213* locus, two of which were missense mutations, that were associated with the disease (Gormley et al. 2016). The underlying mechanisms of migraine are poorly understood. However, the neurovascular theory holds that a complex series of neural and vascular events initiates migraine (May and Goadsby 1999). These facts suggest that variants in the *RNF213* could contribute to the development of several neurovascular diseases.

RNF213 is an E3 ubiquitin-protein ligase with two AAA+ ATPase domains, which are characteristic of energy-dependent unfoldases (Koizumi et al. 2016). It has been shown that RNF213 is involved in angiogenesis by promoting vessel regression (Scholz et al. 2016). Hitomi et al. showed that the angiogenic activities of iPSC-derived vascular endothelial cells (iPSECs) from MMD patients and carriers were lower than in subjects who did not carry the *RNF213* p.R4810K allele. Furthermore, they showed that overexpression of *RNF213* p.R4810K inhibited angiogenic activity and proliferation of human umbilical vein endothelial cells (HUVECs) while overexpression of normal RNF213 did not (Hitomi et al. 2013). The underlying pathogenic mechanism of RNF213 in vascular disease (whether loss- or toxic gain-of-function) remains unclear, a number of *in vivo* knockdown or overexpression models have demonstrated conflicting results on vasculature (Liu et al. 2011) (Sonobe et al. 2014).

Both *SLCA26A11* and *RNF213* could be involved in neurovascular disorders. *SLCA26A11* may act in the neuron-environment homeostasis after an acute ischemic event by worsening the cytotoxic edema around the original lesion. SLC26A11 is a member of the solute linked carrier 26 family of sulfate/anion exchangers. After an acute injury, such as an ischemic stroke, the depolarized neuronal membranes drive an influx of Na^+^ within the cell. Membrane depolarization also activates the voltage-gated SLC26A11 chloride channel, which leads to Cl^−^ accumulation within the cells (Rungta et al. 2015). The increase of cytoplasmic NaCl generates an osmotic imbalance that leads to water influx, which causes a cytotoxic edema with neuronal swelling and subsequent cell death (Rungta et al. 2015). Lower SLC26A11 expression in vascular tissues secondary to most of its genetic variation could be interpreted as a potential source of dysregulation of intracellular chloride transport.

However, *SLC26A11* rs6565653 tags multiple *RNF213* variants and suggests that the young onset ischemic stroke GWAS signal could be tagging *RNF213* gene variation. This would further support the role of *RNF213* in the development of multiple neurovascular disorders, including MMD, IA and migraine and an increased risk and/or worse prognosis of an ischemic stroke in a population specific manner. In any case, additional studies, including expression analyses in vascular tissues from patients with neurovascular disorders, are warranted to further study the role of both genes in the pathophysiology of neurovascular disorders.

## Acknowledgments

The authors thank the patients and families who donated DNA samples for this work. This study was supported by the Mayo Foundation for Medical Education and Research, benefactors Randall S. and Friedgard D. Acree, James and Esther King Foundation, New Investigator Award from the Department of Health, the American Heart Association, the Mayo Clinic Office of Health Disparities Research, the Myron and Jane Hanley Award for Stroke Research and the Joe Niekro Foundation.

