## Supplemental Figure 1 for "*RNF213* variation, a broader role in neurovascular disease in Caucasian and Japanese populations"

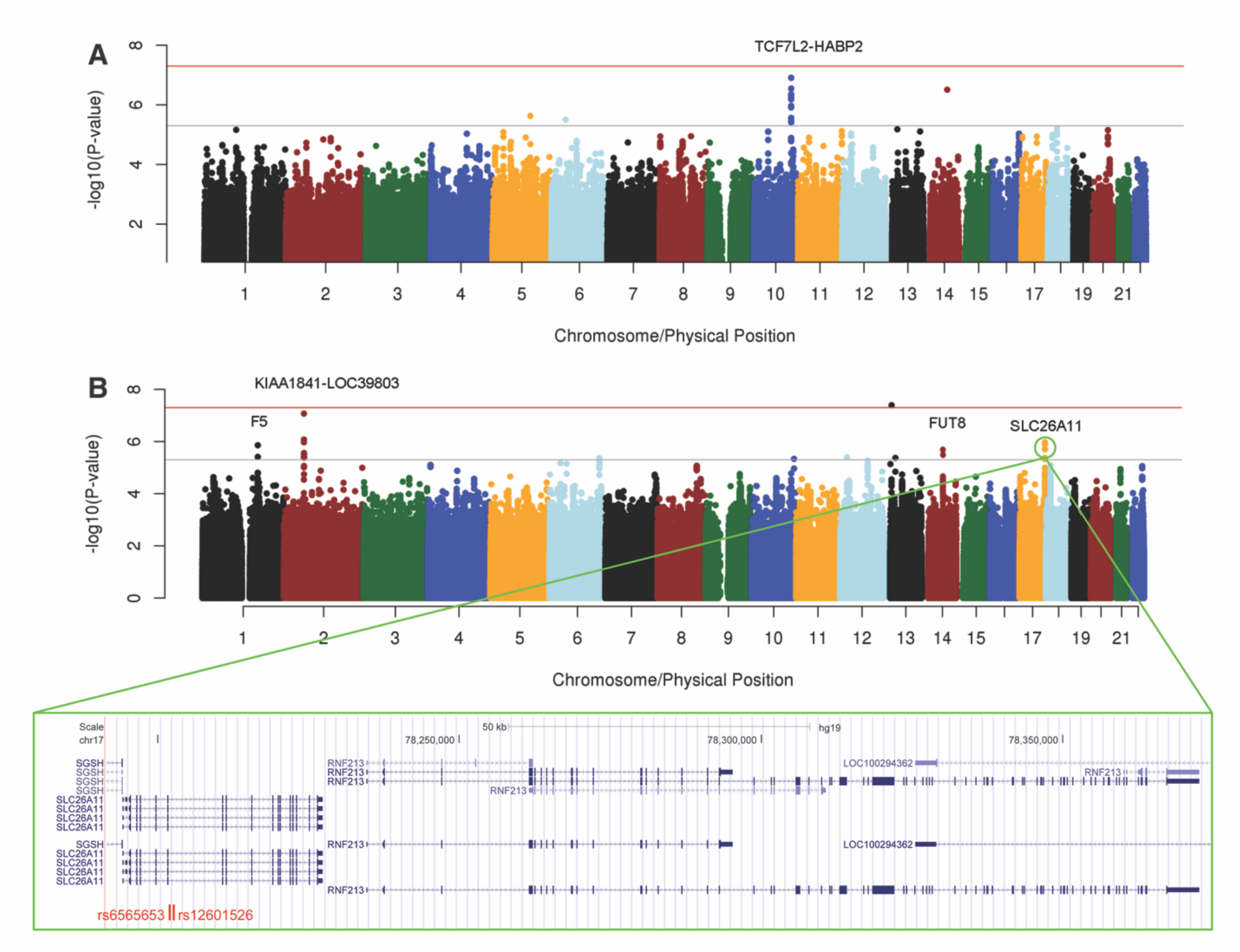


**Figure S1.** Genome-wide association results of early-onset ischemic stroke based on transethnic meta-analysis (A) and European-only meta-analysis (B). Red line: P=5×10−8; grey line: P=5×10−6 (from Cheng et al. Stroke 2016; Copyright © American Heart Association, Inc. All rights reserved). *SLC26A11* signal in the European-only meta-analysis on chromosome 17 is highlighted with a green circle and a Genome Browser zoomed in view of the region is given; rs6565653 and rs12601526 are highlighted in red.
