## Supplemental Figure 2 for "*RNF213* variation, a broader role in neurovascular disease in Caucasian and Japanese populations"

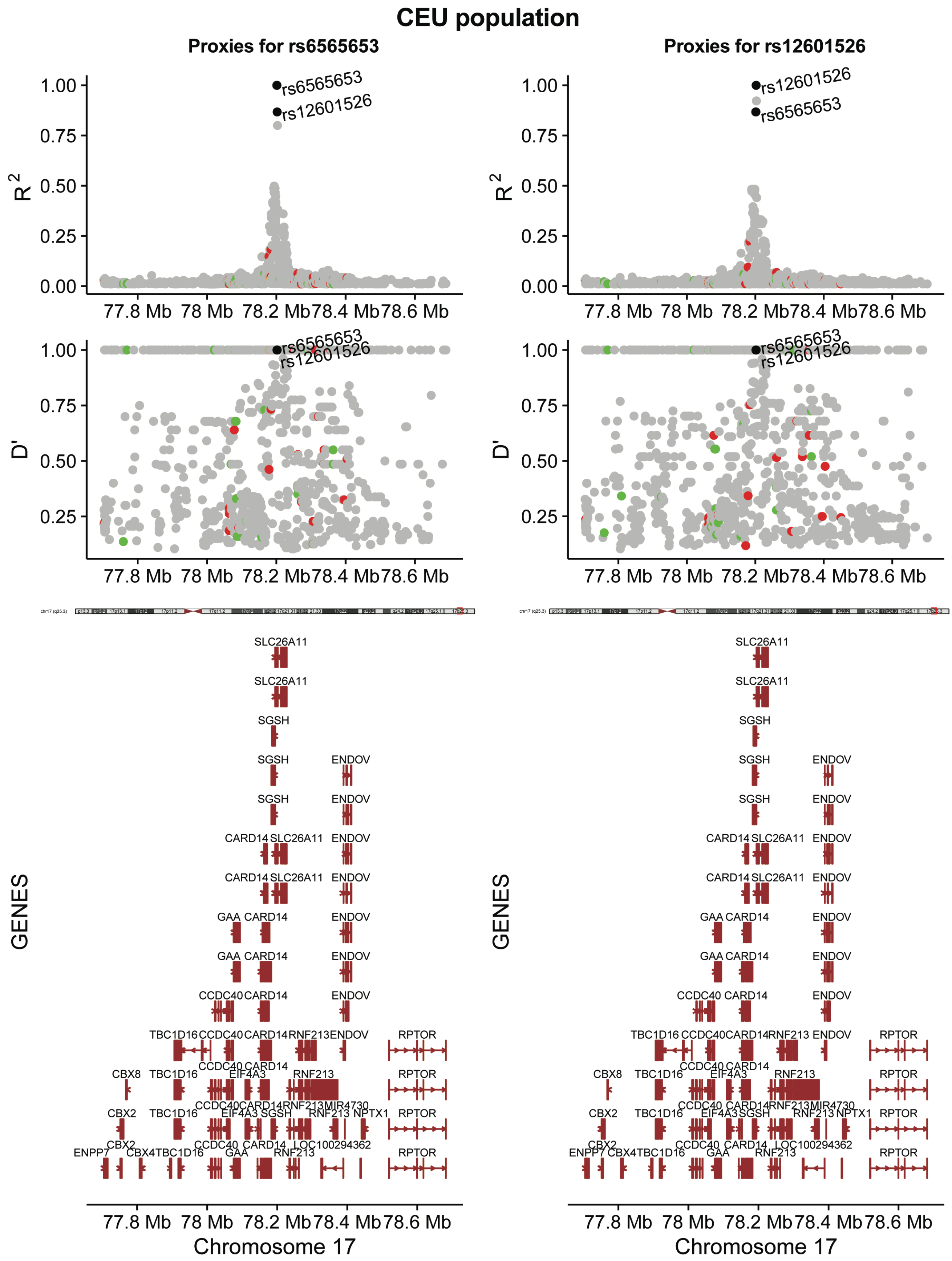


**Figure S2.** $R^2$ and D’ values for proxy SNPs in CEU population for rs6565653 and rs12601526 *SLCA26A11* SNPs. Dot color meaning: red = missense mutation, green = synonymous mutation, grey = non-coding-mutation.
