## Supplemental Figure 4 for "*RNF213* variation, a broader role in neurovascular disease in Caucasian and Japanese populations"

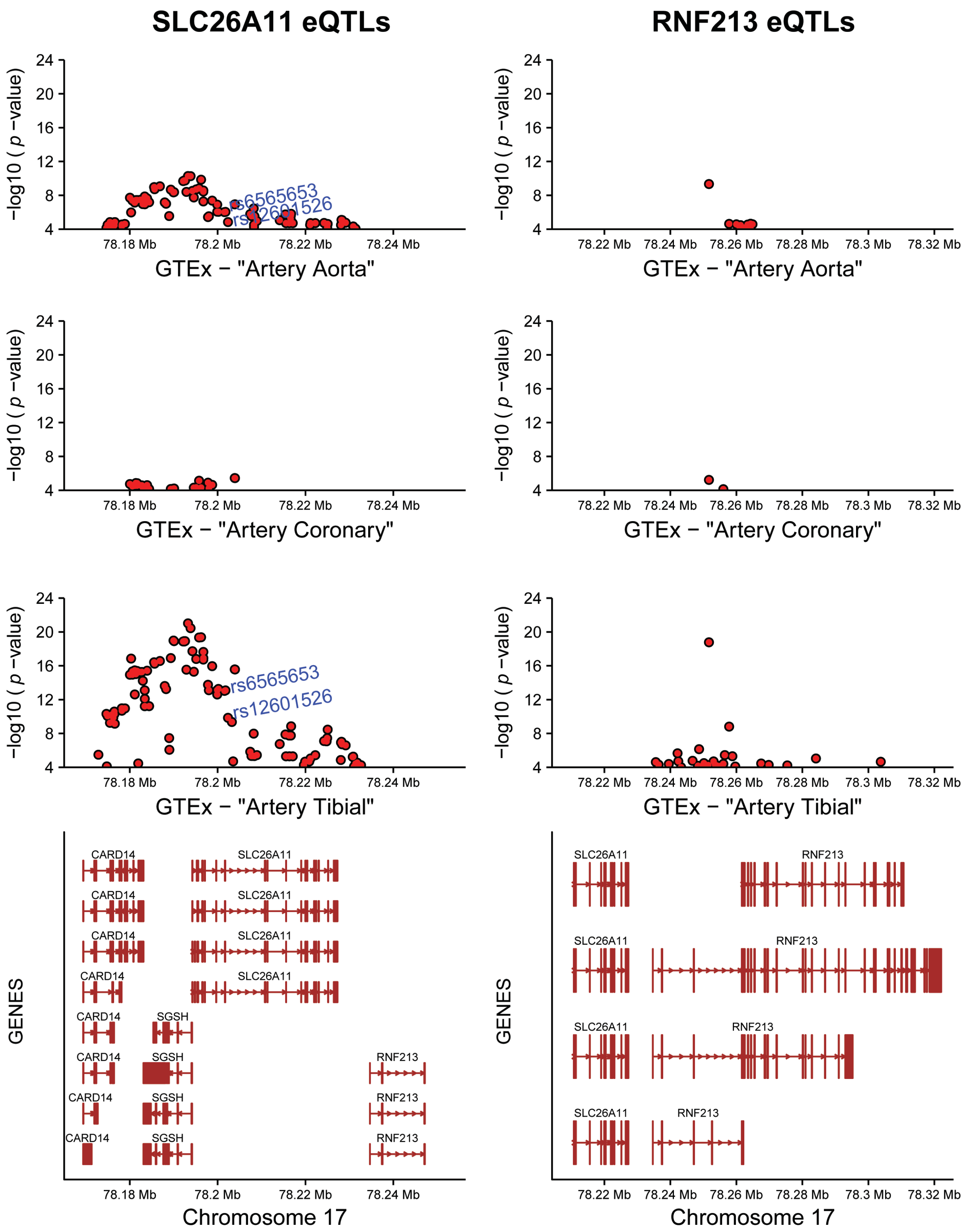


**Figure S4.** Significant local expression quantitative trait loci (cis-eQTL) values for *SLC26A11* and *RNF213* SNPs, affecting arterial tissues. Red dots are significant cis-eQTLs (at false discovery rate <5%) for *SLC26A11* and *RNF213* genes in aortic, coronary and tibial artery tissues, respectively. Each plot has been generated with an up and downstream 25kb flanking region. Figures elaborated with data from the Portal for the Genotype-Tissue Expression project (GTEx). *CARD14* = Caspase Recruitment Domain Family Member 14; *SGSH* = N-Sulfoglucosamine Sulfohydrolase; *SLC26A11* = Solute Carrier Family 26 Member 11; *RNF213* = Ring Finger Protein 213; Mb = megabase.
