## Supplemental Figure 5 for "*RNF213* variation, a broader role in neurovascular disease in Caucasian and Japanese populations"

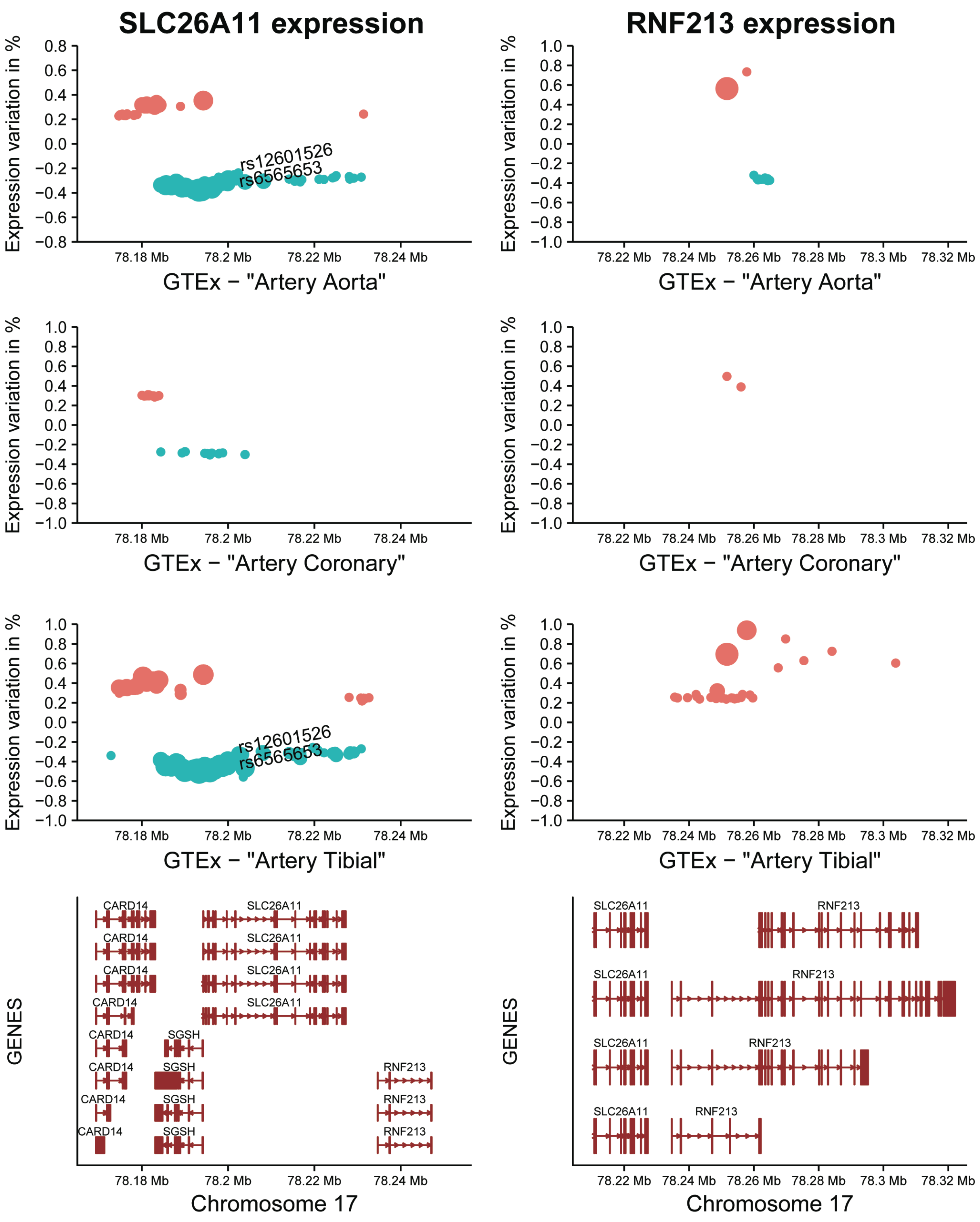


**Figure S5.** Expression changes in SLC26A11 and RNF213 due to SNPs effect in arterial tissues. Red dots show overexpression and blue dots show underexpression, respectively, of SLC26A11 and RNF213 proteins in aortic, coronary and tibial artery tissues. The size of each dot represents the -log10(*p*-value) between the SNP and the expression change. Figures elaborated with data from the Portal for the Genotype-Tissue Expression project (GTEx). *CARD14* = Caspase Recruitment Domain Family Member 14; *SGSH* = N-Sulfoglucosamine Sulfohydrolase; *SLC26A11* = Solute Carrier Family 26 Member 11; *RNF213* = Ring Finger Protein 213; Mb = megabase.
