## Supplemental Table 1 for "*RNF213* variation, a broader role in neurovascular disease in Caucasian and Japanese populations"

**Table S1. Chromosome, position and allele frequencies for SNPs used in the LD analysis in CEU population.**

| rsID | Chr | Position | EUR AF |
|---|---|---|---|
| rs7207000 | 17 | 78192358 | 0.3559 |
| rs6565649 | 17 | 78192414 | 0.3549 |
| rs6565650 | 17 | 78192571 | 0.3569 |
| rs6565651 | 17 | 78193252 | 0.3539 |
| rs4889841 | 17 | 78193831 | 0.3628 |
| rs12946329 | 17 | 78195772 | 0.3161 |
| rs7224212 | 17 | 78199903 | 0.2445 |
| rs6565653 | 17 | 78201783 | 0.4761 |
| rs12601526 | 17 | 78202358 | 0.4284 |
| rs7208687 | 17 | 78206586 | **0.3449** |
| rs6420487 | 17 | 78207690 | **0.2853** |
| rs7216577 | 17 | 78208374 | **0.2853** |
| rs3924207 | 17 | 78210776 | 0.0457 |
| rs11658788 | 17 | 78211732 | **0.3449** |
| rs6420489 | 17 | 78212206 | 0.4394 |
| rs12942462 | 17 | 78215566 | 0.2266 |
| rs12949090 | 17 | 78217097 | 0.2276 |
| rs7406843 | 17 | 78219065 | 0.5765 |
| rs9904116 | 17 | 78222661 | 0.673 |
| rs8064518 | 17 | 78223103 | 0.7167 |
| rs8076666 | 17 | 78223817 | 0.8638 |
| rs12938473 | 17 | 78224232 | 0.3111 |
| rs8078855 | 17 | 78225055 | 0.3241 |
| rs11650413 | 17 | 78228181 | 0.4304 |
| rs9915508 | 17 | 78230756 | 0.495 |
| rs9903254 | 17 | 78232650 | 0.4264 |
| rs7217421 | 17 | 78235460 | 0.5139 |
| rs4889968 | 17 | 78236804 | 0.0288 |
| rs9916351 | 17 | 78239532 | 0.4533 |
| rs12603583 | 17 | 78244237 | 0.0258 |
| rs9902013 | 17 | 78246462 | 0.0447 |
| rs12947578 | 17 | 78246734 | 0.4066 |
| rs9890007 | 17 | 78251427 | **0.4602** |
| rs9901491 | 17 | 78251518 | **0.0249** |
| rs7219131 | 17 | 78252770 | **0.4602** |
| rs8069705 | 17 | 78254067 | 0.4404 |
| rs12601730 | 17 | 78255597 | **0.0249** |
| rs12601017 | 17 | 78256912 | **0.0249** |
| rs7214614 | 17 | 78258794 | 0.4165 |
| rs11869363 | 17 | 78266759 | 0.2893 |
| rs12937242 | 17 | 78268294 | **0.1531** |
| rs9909720 | 17 | 78269111 | **0.1531** |
| rs9905727 | 17 | 78272814 | 0.164 |
| rs8068939 | 17 | 78275265 | 0.2614 |
| rs8076804 | 17 | 78275482 | 0.162 |
| rs12451223 | 17 | 78279334 | **0.2157** |
| rs8066993 | 17 | 78280030 | **0.2157** |
| rs8081176 | 17 | 78283987 | 0.3151 |
| rs9674807 | 17 | 78286177 | 0.2883 |
| rs7501761 | 17 | 78290185 | 0.671 |
| rs6565666 | 17 | 78292945 | 0.1879 |
| rs7222014 | 17 | 78293469 | 0.499 |
| rs7225029 | 17 | 78297779 | 0.0666 |
| rs4889843 | 17 | 78299319 | 0.5 |
| rs4889844 | 17 | 78299497 | 0.7266 |
| rs7501563 | 17 | 78300546 | **0.7048** |
| rs9913473 | 17 | 78300821 | **0.7048** |
| rs4075499 | 17 | 78300859 | **0.7048** |
| rs9907978 | 17 | 78301380 | **0.7058** |
| rs9908583 | 17 | 78301474 | **0.7058** |
| rs8082521 | 17 | 78302157 | **0.7058** |
| rs4890007 | 17 | 78304552 | **0.0348** |
| rs4890008 | 17 | 78305619 | **0.0348** |
| rs10782008 | 17 | 78305871 | **0.67** |
| rs8074015 | 17 | 78306280 | 0.7028 |
| rs8072917 | 17 | 78309108 | 0.667 |
| rs7211876 | 17 | 78311360 | 0.2147 |
| rs4890009 | 17 | 78311508 | 0.7018 |
| rs34399489 | 17 | 78313562 | 0.0179 |
| rs4890010 | 17 | 78316179 | **0.67** |
| rs11150856 | 17 | 78322882 | **0.1531** |
| rs9899221 | 17 | 78323500 | 0.0189 |
| rs12051723 | 17 | 78324259 | 0.161 |
| rs11655038 | 17 | 78327229 | 0.0885 |
| rs7216493 | 17 | 78327358 | 0.7346 |
| rs11656211 | 17 | 78327745 | 0.0964 |
| rs8067292 | 17 | 78333840 | **0.5984** |
| rs4078429 | 17 | 78335372 | 0.0437 |
| rs7221291 | 17 | 78344343 | **0.5984** |
| rs8070106 | 17 | 78344446 | **0.5984** |
| rs4889847 | 17 | 78346548 | 0.0487 |
| rs6565681 | 17 | 78348494 | 0.84 |
| rs9898470 | 17 | 78351368 | 0.0417 |
| rs4889848 | 17 | 78354661 | 0.8419 |
| rs7224239 | 17 | 78355176 | 0.4334 |
| rs12944088 | 17 | 78357478 | 0.0159 |
| rs4890018 | 17 | 78360001 | 0.0646 |
| rs7223701 | 17 | 78362651 | 0.0915 |
| rs8072774 | 17 | 78363054 | **0.0905** |
| rs3185057 | 17 | 78363847 | **0.0905** |
| rs8359 | 17 | 78368051 | 0.0944 |
| rs13341671 | 17 | 78374412 | 0.5676 |

SNPs with at least another variant with the same allele frequency are highlighted in red. EUR AF = European allele frequency
