## Supplemental Table 2 for "*RNF213* variation, a broader role in neurovascular disease in Caucasian and Japanese populations"

**Table S2. Chromosome, position and allele frequencies for the SNPs used in the LD analysis in the JPT population.**

| rsID | Chr | Position | JPT AF |
|---|---|---|---|
| rs7207000 | 17 | 78192358 | **0.3998** |
| rs6565649 | 17 | 78192414 | **0.3998** |
| rs6565650 | 17 | 78192571 | **0.3998** |
| rs6565651 | 17 | 78193252 | 0.4028 |
| rs4889841 | 17 | 78193831 | 0.4008 |
| rs12946329 | 17 | 78195772 | 0.3929 |
| rs7224212 | 17 | 78199903 | **0.38** |
| rs6565653 | 17 | 78201783 | 0.9752 |
| rs12601526 | 17 | 78202358 | 0.9712 |
| rs7208687 | 17 | 78206586 | 0.6022 |
| rs6420487 | 17 | 78207690 | **0.3472** |
| rs7216577 | 17 | 78208374 | **0.3472** |
| rs11658788 | 17 | 78211732 | 0.6002 |
| rs6420489 | 17 | 78212206 | 0.621 |
| rs8079641 | 17 | 78212420 | 0.0377 |
| rs12942462 | 17 | 78215566 | 0.3462 |
| rs12949090 | 17 | 78217097 | 0.3452 |
| rs7406843 | 17 | 78219065 | 0.9464 |
| rs9904116 | 17 | 78222661 | 0.9454 |
| rs8064518 | 17 | 78223103 | 0.994 |
| rs8076666 | 17 | 78223817 | 0.999 |
| rs12938473 | 17 | 78224232 | **0.6161** |
| rs8078855 | 17 | 78225055 | 0.5615 |
| rs11650413 | 17 | 78228181 | 0.4236 |
| rs9915508 | 17 | 78230756 | 0.4196 |
| rs9903254 | 17 | 78232650 | 0.1627 |
| rs7217421 | 17 | 78235460 | **0.6161** |
| rs4889968 | 17 | 78236804 | 0.2024 |
| rs9916351 | 17 | 78239532 | 0.6171 |
| rs12603583 | 17 | 78244237 | 0.1994 |
| rs9902013 | 17 | 78246462 | 0.3294 |
| rs12947578 | 17 | 78246734 | 0.3115 |
| rs9890007 | 17 | 78251427 | 0.6359 |
| rs9901491 | 17 | 78251518 | 0.2163 |
| rs7219131 | 17 | 78252770 | 0.6349 |
| rs8069705 | 17 | 78254067 | 0.505 |
| rs12601730 | 17 | 78255597 | **0.2034** |
| rs12601017 | 17 | 78256912 | **0.2034** |
| rs7214614 | 17 | 78258794 | **0.3006** |
| rs11869363 | 17 | 78266759 | 0.4851 |
| rs12937242 | 17 | 78268294 | **0.0704** |
| rs9909720 | 17 | 78269111 | **0.0704** |
| rs9905727 | 17 | 78272814 | 0.0764 |
| rs8068939 | 17 | 78275265 | 0.0923 |
| rs8076804 | 17 | 78275482 | 0.0774 |
| rs12451223 | 17 | 78279334 | 0.3075 |
| rs8066993 | 17 | 78280030 | **0.3006** |
| rs8081176 | 17 | 78283987 | 0.3244 |
| rs9674807 | 17 | 78286177 | 0.3046 |
| rs7501761 | 17 | 78290185 | 0.6339 |
| rs6565666 | 17 | 78292945 | 0.1468 |
| rs7222014 | 17 | 78293469 | 0.2192 |
| rs4889843 | 17 | 78299319 | 0.1944 |
| rs4889844 | 17 | 78299497 | **0.3948** |
| rs7501563 | 17 | 78300546 | **0.2341** |
| rs9913473 | 17 | 78300821 | **0.2341** |
| rs4075499 | 17 | 78300859 | **0.2341** |
| rs9907978 | 17 | 78301380 | **0.2341** |
| rs9908583 | 17 | 78301474 | 0.2351 |
| rs8082521 | 17 | 78302157 | 0.2262 |
| rs10782008 | 17 | 78305871 | 0.3879 |
| rs8074015 | 17 | 78306280 | **0.3948** |
| rs8072917 | 17 | 78309108 | 0.3661 |
| rs7211876 | 17 | 78311360 | 0.3214 |
| rs4890009 | 17 | 78311508 | 0.375 |
| rs4890010 | 17 | 78316179 | **0.38** |
| rs11150856 | 17 | 78322882 | 0.0317 |
| rs12051723 | 17 | 78324259 | 0.248 |
| rs7216493 | 17 | 78327358 | 0.5486 |
| rs8067292 | 17 | 78333840 | 0.3423 |
| rs4078429 | 17 | 78335372 | 0.1736 |
| rs7221291 | 17 | 78344343 | 0.3869 |
| rs8070106 | 17 | 78344446 | 0.3849 |
| rs6565681 | 17 | 78348494 | 0.5972 |
| rs9898470 | 17 | 78351368 | 0.1766 |
| rs4889848 | 17 | 78354661 | 0.5794 |
| rs7224239 | 17 | 78355176 | 0.4077 |
| rs4890018 | 17 | 78360001 | 0.0288 |
| rs3185057 | 17 | 78363847 | 0.0367 |
| rs13341671 | 17 | 78374412 | 0.3819 |

SNPs with at least another variant with the same allele frequency are highlighted in red. EAS AF = East Asian allele frequency
