## Supplemental R code for "*RNF213* variation, a broader role in neurovascular disease in Caucasian and Japanese populations"

Author Note:

### Supplementary File S1 for paper: *RNF213* variation, a broader role in neurovascular disease in Caucasian and Japanese populations - R code

March 24th, 2020

This file contains all the analyses that were performed in order to generate the results and plots from the current paper.

#### Load required libraries and set working directory

```r
# Load required libraries
library(ggplot2)
require(extrafont)
library(ggbio)
library(grid)
library(gridExtra)
library(png)
library(gtable)
library(cowplot)
library(ensembldb)
library(genetics)
library(GenomicRanges)
library(grDevices)
library(Homo.sapiens)
library(BSgenome.Hsapiens.UCSC.hg19)
library(LDheatmap)
library(plyr)
library(raster)
library(reshape)
library(Rsamtools)
library(rtracklayer)
library(scales)
library(VariantAnnotation)

# Set working directory
dirname <- getwd()
setwd(dirname)
```

#### LD heatmaps for CEU population (Figure 1A and 1B)

```
# CEU population all common SNPs (n=91)
CEU_chr17 <- as.data.frame(read.table("input_files/CEU_files/CEU_SLC26A11_RN213_genotypes.txt",
                                       header = TRUE))
CEU_data <- as.data.frame(read.table("input_files/CEU_files/CEU_common_SNPs.bed",
                                      header = TRUE))
num_CEU <- ncol(CEU_chr17)
colnames_CEU <- as.vector(CEU_data[,4])

for(i in 1:num_CEU){
        CEU_chr17[,i]<-as.genotype(CEU_chr17[,i])
}

plot_title_CEU_r <- as.expression(bquote(atop('Pairwise LD (r'^{~2}~') for common SNPs located in the'~i

plot_title_CEU_D <- as.expression(bquote(atop('Pairwise LD (D\') for common SNPs located in the'~italic(

# Raw LD heatmap r2 CEU
CEU_SNPs_ALL_r <- LDheatmap(CEU_chr17, CEU_data[,2], LDmeasure = "r",
                            title = plot_title_CEU_r, add.map = TRUE,
                            flip = TRUE, color = heat.colors(20), name = "CEULDgrob",
                            add.key = TRUE, newpage = TRUE)
```

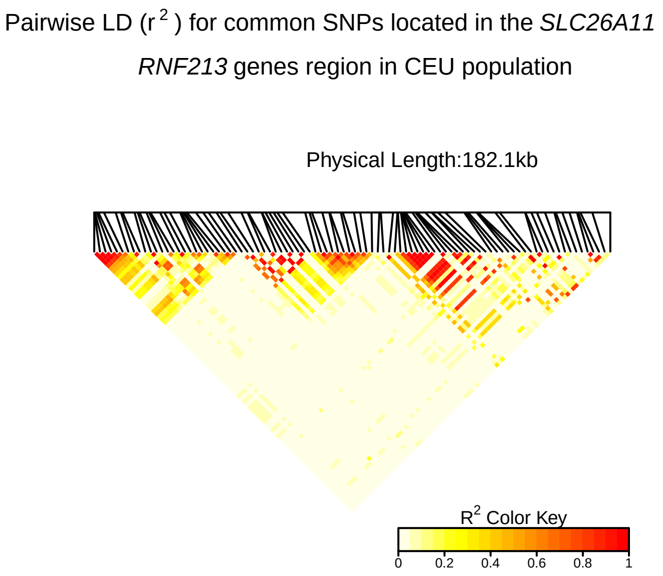


```
# Raw LD heatmap D' CEU
CEU_SNPs_ALL_D <- LDheatmap(CEU_chr17, CEU_data[,2], LDmeasure = "D'",
                            title = plot_title_CEU_D, add.map = TRUE,
                            flip = TRUE, color = heat.colors(20), name = "CEULDgrob",
                            add.key = TRUE, newpage = TRUE)
```

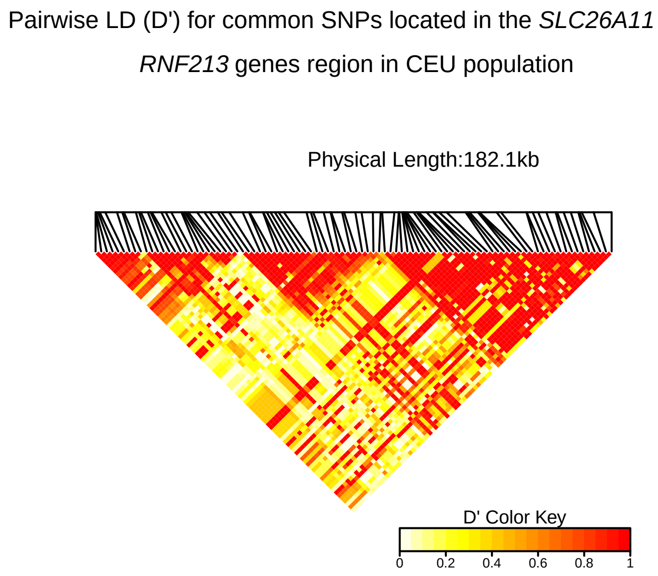


**Figure 1A**

```r
CEU_SNPs_ALL_genes_r <- LDheatmap.addGenes(CEU_SNPs_ALL_r, chr="chr17", genome="hg19")
```

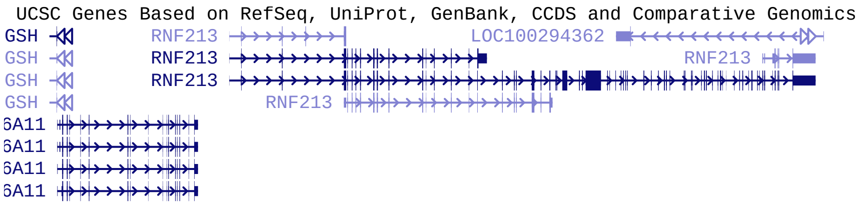

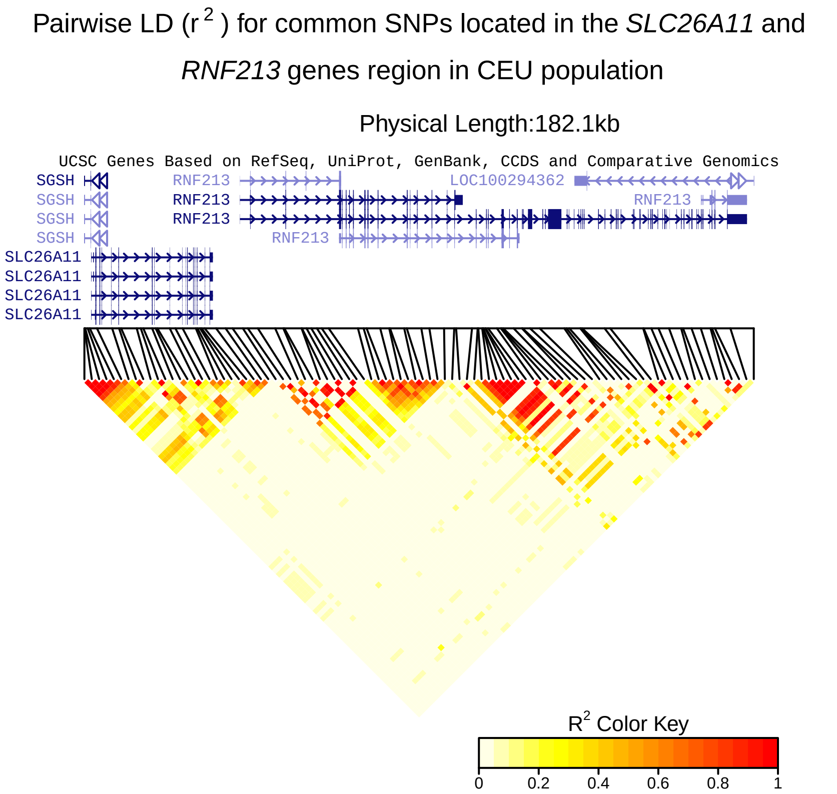


```r
dev.off()
```

```
## null device
##           1
```

#### Figure 1B

```r
CEU_SNPs_ALL_genes_D <- LDheatmap.addGenes(CEU_SNPs_ALL_D, chr="chr17", genome="hg19")
```

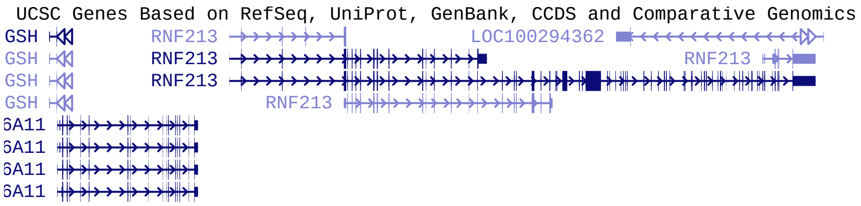

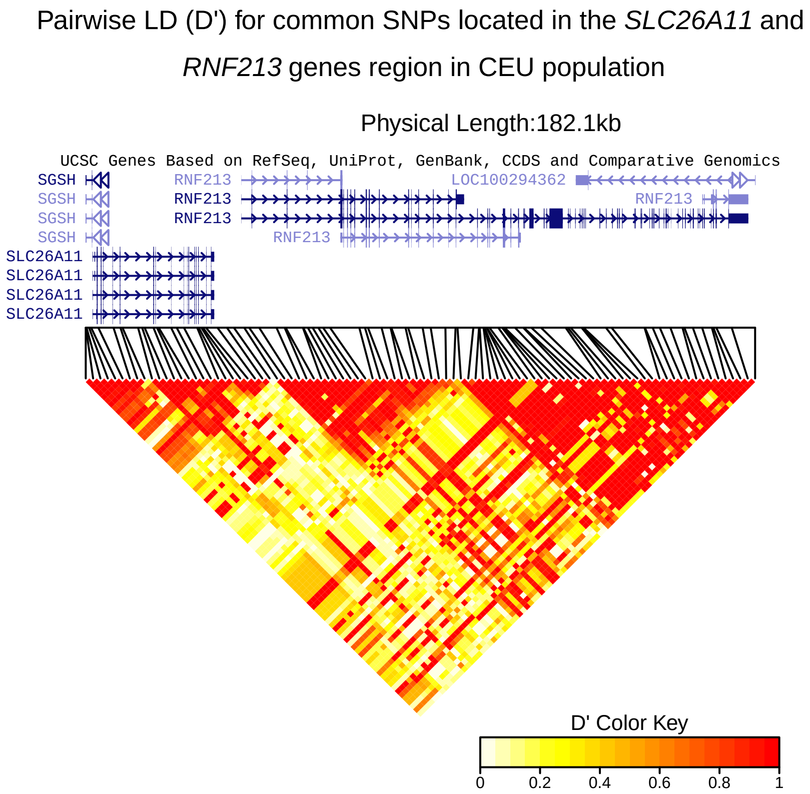


```r
dev.off()
```

```
## null device
##           1
```

### LD heatmaps for JPT population (Figure 1C and 1D)

```r
# JPT population all common SNPs
JPT_chr17 <- as.data.frame(read.table("input_files/JPT_files/JPT_SLC26A11_RN213_genotypes.txt",
                                       header = TRUE))
JPT_data <- as.data.frame(read.table("input_files/JPT_files/JPT_common_SNPs.bed", header = TRUE))
num_JPT<-ncol(JPT_chr17)
colnames_JPT <- as.vector(JPT_data[,4])
class(colnames_JPT)
```

```
## [1] "character"
```

```r
class(JPT_data[,4])
```

```
## [1] "factor"
```

```r
for(j in 1:num_JPT){
        JPT_chr17[,j]<-as.genotype(JPT_chr17[,j])
}

plot_title_JPT_r <- as.expression(bquote(atop('Pairwise LD (r'^{~2}~') for common SNPs located in the'~i
```

```
plot_title_JPT_D <- as.expression(bquote(atop('Pairwise LD (D\') for common SNPs located in the'~italic(

# Raw LD heatmap r2 JPT
JPT_SNPs_ALL_r <- LDheatmap(JPT_chr17, JPT_data[,2], LDmeasure = "r",
                            title = plot_title_JPT_r, add.map = TRUE,
                            flip = TRUE, color = heat.colors(20), name = "JPTLDgrob",
                            add.key = TRUE, newpage = TRUE)
```

Pairwise LD ($r^2$) for common SNPs located in the *SLC26A11*

*RNF213* genes region in JPT population

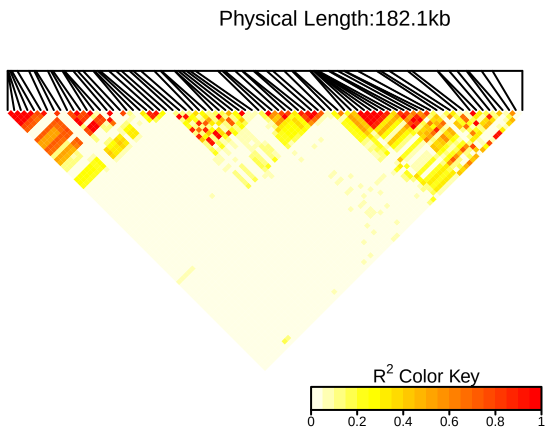


```
# Raw LD heatmap D' JPT
JPT_SNPs_ALL_D <- LDheatmap(JPT_chr17, JPT_data[,2], LDmeasure = "D'",
                          title = plot_title_JPT_D, add.map = TRUE,
                          flip = TRUE, color = heat.colors(20), name = "JPTLDgrob",
                          add.key = TRUE, newpage = TRUE)
```

Pairwise LD (D') for common SNPs located in the *SLC26A11*

*RNF213* genes region in JPT population

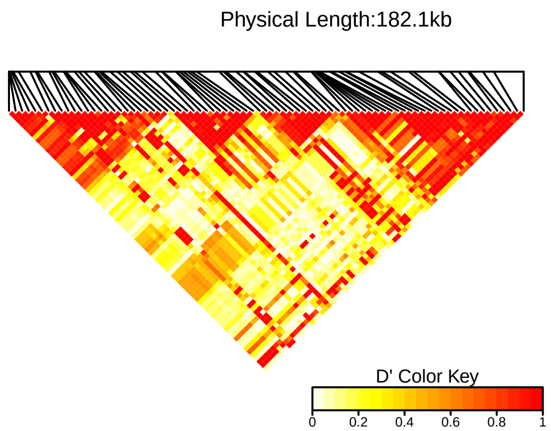

#### Figure 1C

```r
JPT_SNPs_ALL_genes_r <- LDheatmap.addGenes(JPT_SNPs_ALL_r, chr="chr17", genome="hg19")
```

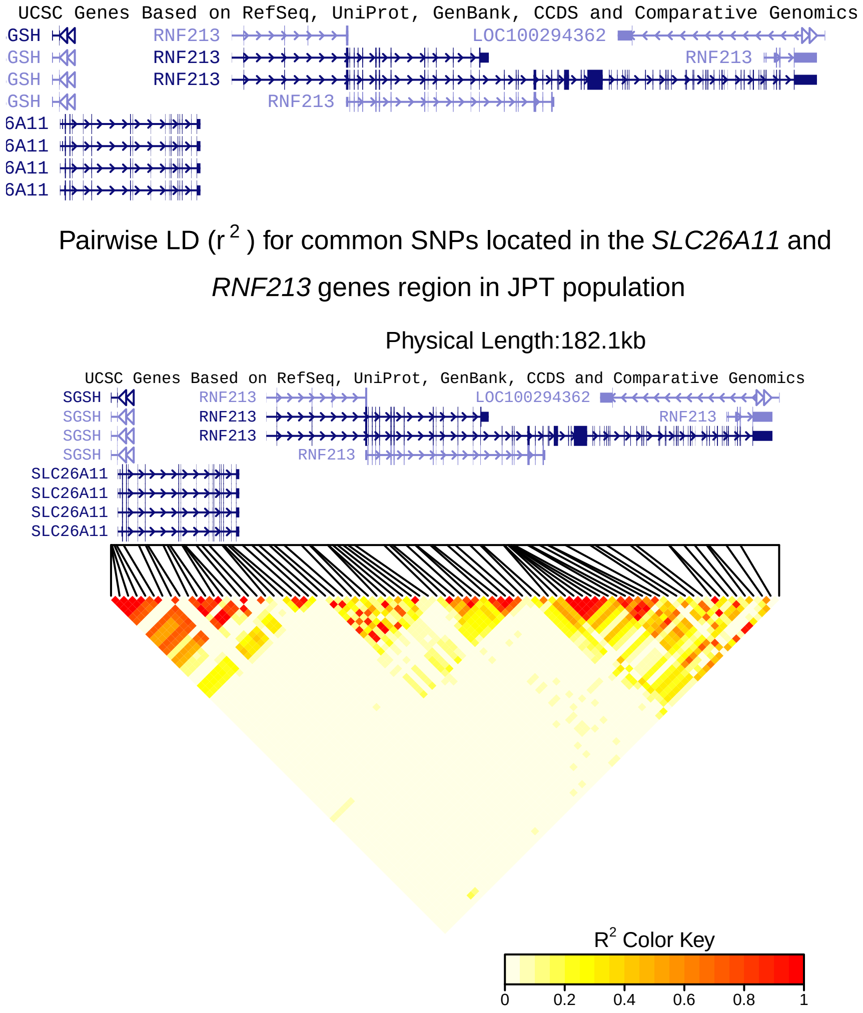


```r
dev.off()
```

```
## null device
##           1
```

#### Figure 1D

```r
JPT_SNPs_ALL_genes_D <- LDheatmap.addGenes(JPT_SNPs_ALL_D, chr="chr17", genome="hg19")
```

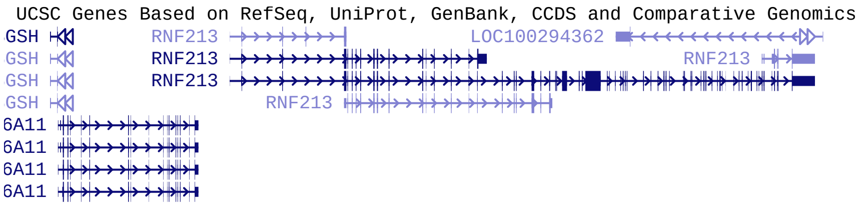


Pairwise LD (D') for common SNPs located in the *SLC26A11* and *RNF213* genes region in JPT population

Physical Length:182.1kb

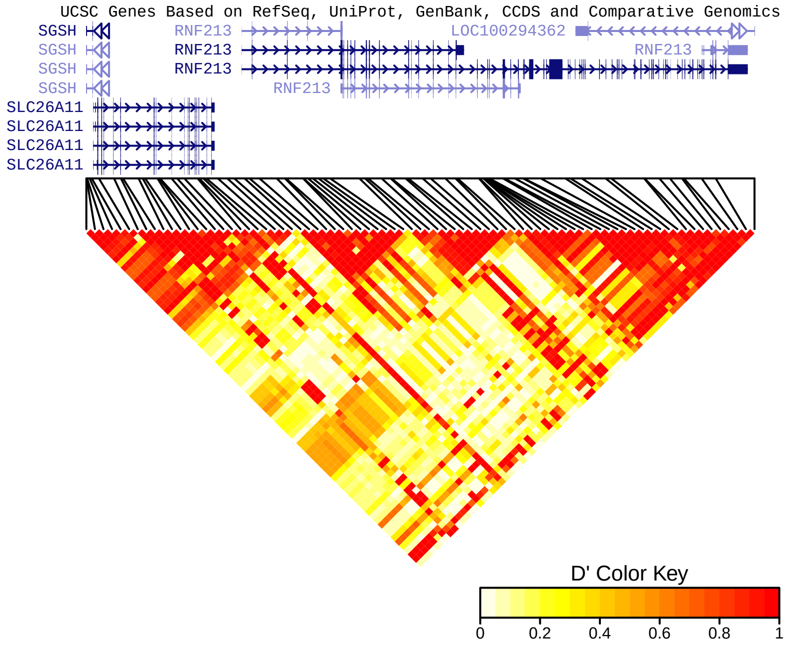


```
dev.off()
```

```
## null device
##           1
```

The four plots of Figure 1 were merged together using Illustrator software and the names of the SNPs of interest were added manually. The chromosome 17 ideogram for each of these plots was added manually also.

### Proxy SNPs for CEU population

Get proxy SNPs for CEU population for for rs6565653 and rs12601526

```
data(hg19IdeogramCyto, package = "biovizBase")
data(hg19Ideogram, package = "biovizBase")
data(genesymbol, package = "biovizBase")

hg19 <- keepSeqlevels(hg19IdeogramCyto, paste0("chr", c(1:22, "X", "Y")))

# Set working directory
```

```
dirname <- getwd()
setwd(dirname)


# Plots for rs6565653 in CEU population
CEU_6565653_df <- as.data.frame(read.table("input_files/CEU_files/Proxy SNPs rs6565653 CEU population.tx
                                            sep = "\t", header = TRUE,
                                            na.strings = c(".", "NA"),
                                            stringsAsFactors = FALSE))

CEU_6565653_df$Function[is.na(CEU_6565653_df$Function)] <- "N/A"

range_start<-head(CEU_6565653_df$Start,n=1)
range_end<-tail(CEU_6565653_df$Start,n=1)
scale_combined <- GRanges('chr17', IRanges(start = range_start, end = range_end))

legend_title <- "SNP function"

y_lab_r <- as.expression(bquote('R'^{~2}~''))

rs6565653_r <- ggplot(CEU_6565653_df, aes(Start,R2)) +
        xlim (scale_combined) +
        geom_point(size = 2, aes(colour = Function)) +
        geom_point(size = 2, data=subset(CEU_6565653_df, DisplayName == "Y"),
                   aes(Start, R2), color = "black") +
        geom_text(data=subset(CEU_6565653_df, DisplayName == "Y"),
                  aes(Start, R2,label=ID, hjust = 0, angle = 10)) +
        labs(y = y_lab_r, color = "SNP Function\n") +
        scale_color_manual(labels = c("missense", "unkwnown", "nonsense", "synonymous"),
                           values = c("red", "grey", "orange", "green")) +
        scale_x_sequnit("Mb") +
        theme(axis.title.x = element_blank (), legend.position = "none")
fixed(rs6565653_r) <- TRUE
rs6565653_r
```

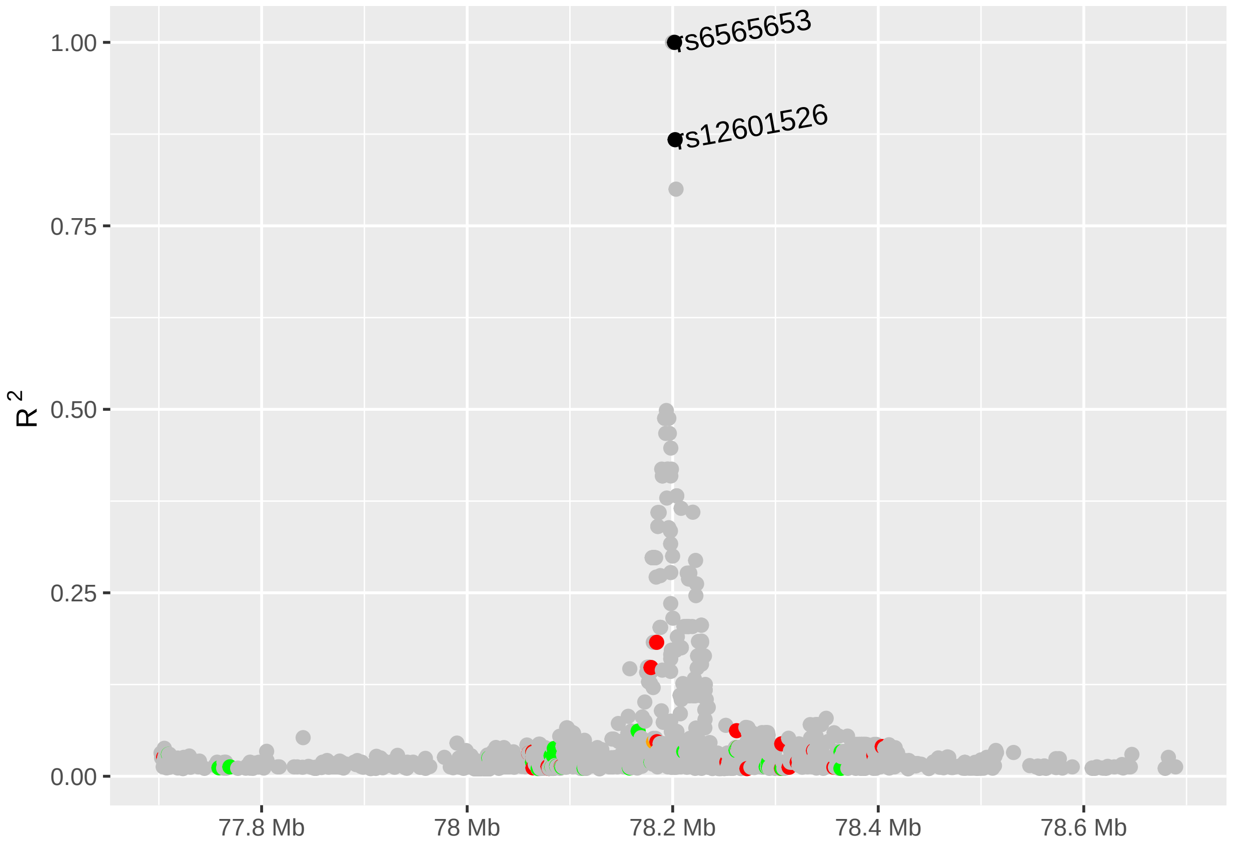


```r
rs6565653_D <- ggplot(CEU_6565653_df, aes(Start,Dprime)) +
        xlim (scale_combined) +
        geom_point(size = 2, aes(colour = Function)) +
        geom_point(size = 2, data=subset(CEU_6565653_df, DisplayName == "Y"),
                   aes(Start, Dprime), color = "black") +
        geom_text(data=subset(CEU_6565653_df, DisplayName == "Y"),
                  aes(Start, Dprime,label=ID, hjust = 0, angle = 10)) +
        labs(y = "D\'", color = "SNP Function\n") +
        scale_color_manual(labels = c("missense", "unkwnown", "nonsense", "synonymous"),
                           values = c("red", "grey", "orange", "green")) +
        scale_x_sequnit("Mb")+
        theme(axis.title.x = element_blank (), legend.position = "none")
fixed(rs6565653_D) <- TRUE
rs6565653_D
```

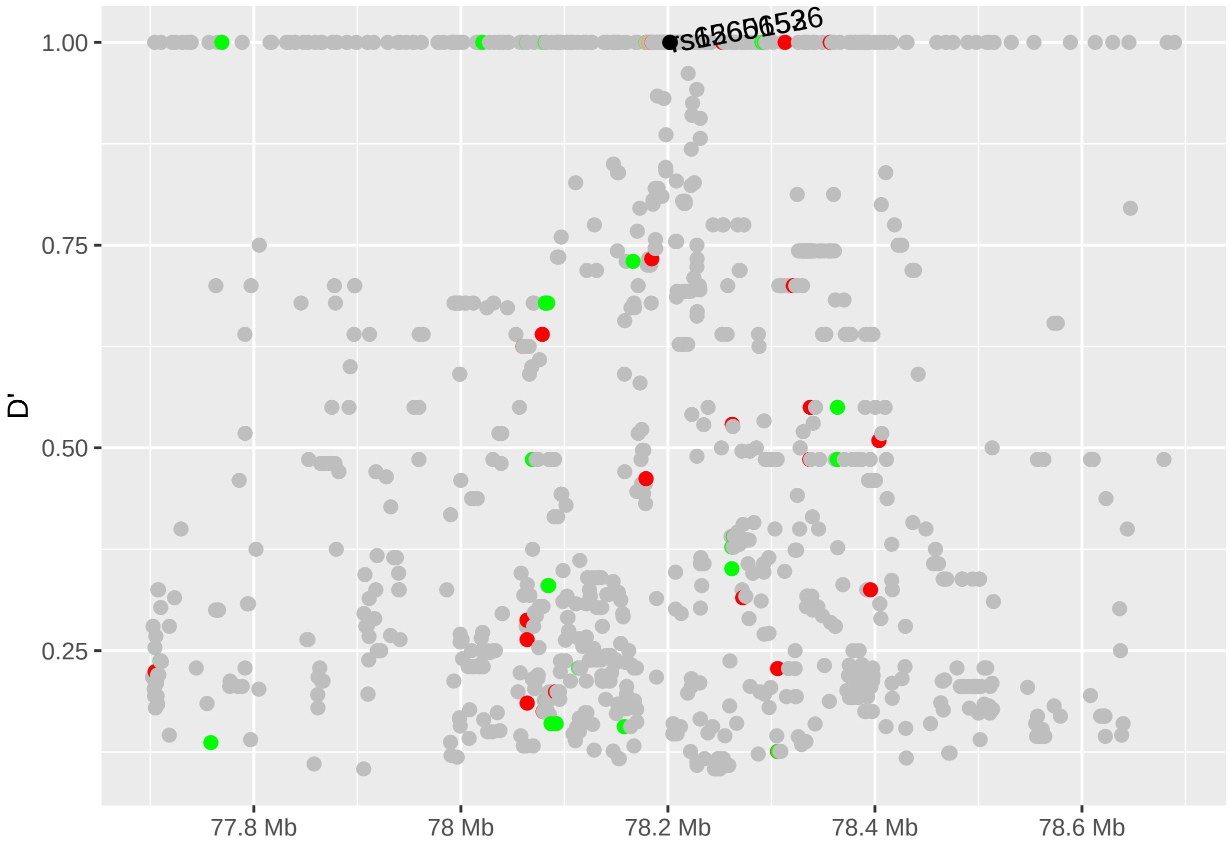


```
# Missing genes: "LOC101928766", "LOC101928738", "MIR4730"
data(genesymbol, package = "biovizBase")
wh <- genesymbol[c("ENPP7", "RPTOR")]
wh <- range(wh, ignore.strand = TRUE)


rs6565653_GENES <- autoplot(Homo.sapiens, which = wh, xlab = "Chromosome 17", ylab = "GENES",
                            label.color = "black", color = "brown", fill = "brown", columns =
                              c("ALIAS", "GO"), scale = "Mb") +
        xlim (scale_combined) +
        scale_x_sequnit("Mb")
rs6565653_GENES
```

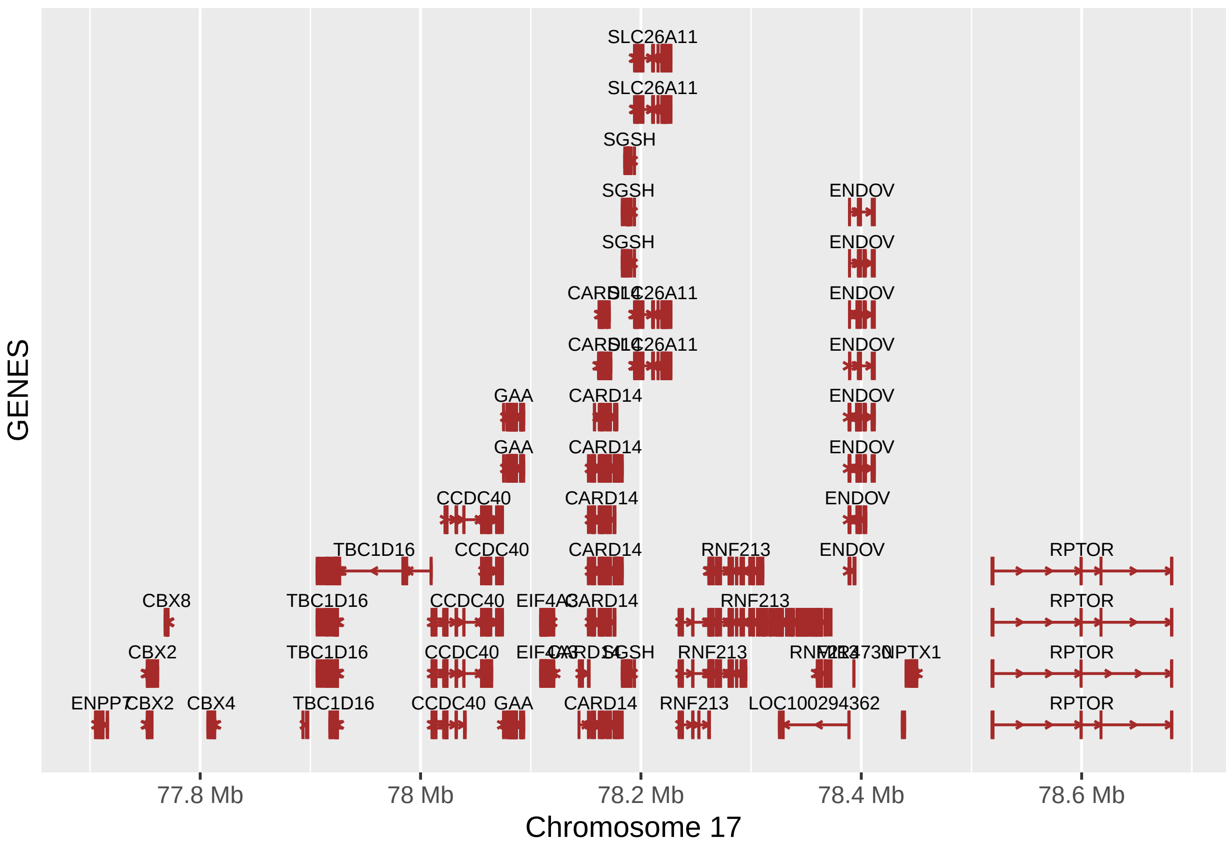


```r
fixed (rs6565653_GENES) <- TRUE

rs6565653_GENES.gg <- rs6565653_GENES@ggplot
rs6565653_GENES.gg
```

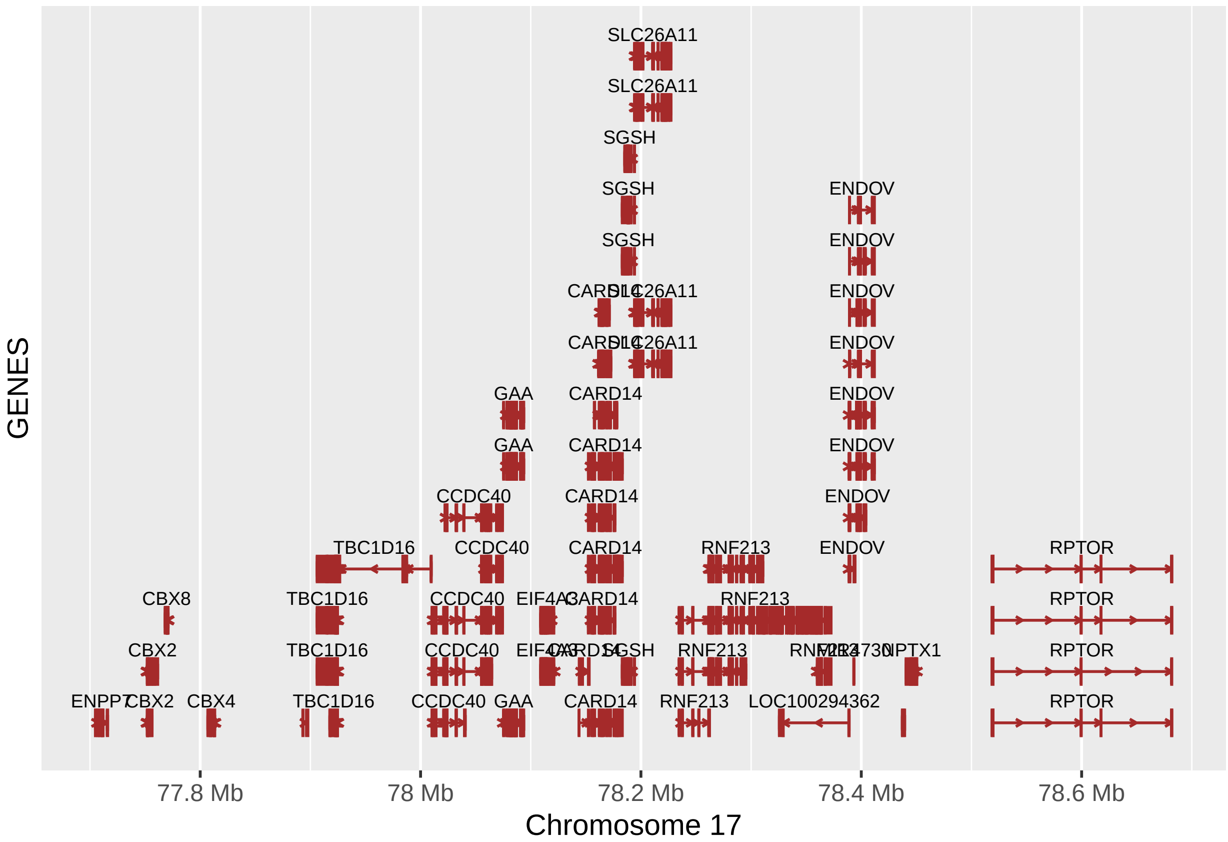


```
# Plots for rs12601526 in CEU population
CEU_12601526_df <- as.data.frame(read.table("input_files/CEU_files/Proxy SNPs rs12601526 CEU population.
                                             sep = "\t", header = TRUE, na.strings = c(".", "NA"),
                                             stringsAsFactors = FALSE))

CEU_12601526_df$Function[is.na(CEU_12601526_df$Function)] <- "N/A"

range_start<-head(CEU_12601526_df$Start,n=1)
range_end<-tail(CEU_12601526_df$Start,n=1)
scale_combined <- GRanges('chr17', IRanges(start = range_start, end = range_end))

legend_title <- "SNP function"

y_lab_r <- as.expression(bquote('R'^{~2}~''))

rs12601526_r <- ggplot(CEU_12601526_df, aes(Start,R2)) +
        xlim (scale_combined) +
        geom_point(size = 2, aes(colour = Function)) +
        geom_point(size = 2, data=subset(CEU_12601526_df, DisplayName == "Y"),
                   aes(Start, R2), color = "black") +
        geom_text(data=subset(CEU_12601526_df, DisplayName == "Y"),
                  aes(Start, R2,label=ID, hjust = 0, angle = 10)) +
        labs(y = y_lab_r, color = "SNP Function\n") +
        scale_color_manual(labels = c("missense", "unkwnown", "synonymous"),
                           values = c("red", "grey", "green")) +
```

```
        scale_x_sequnit("Mb") +
        theme(axis.title.x = element_blank (), legend.position = "none")
fixed(rs12601526_r) <- TRUE
rs12601526_r
```

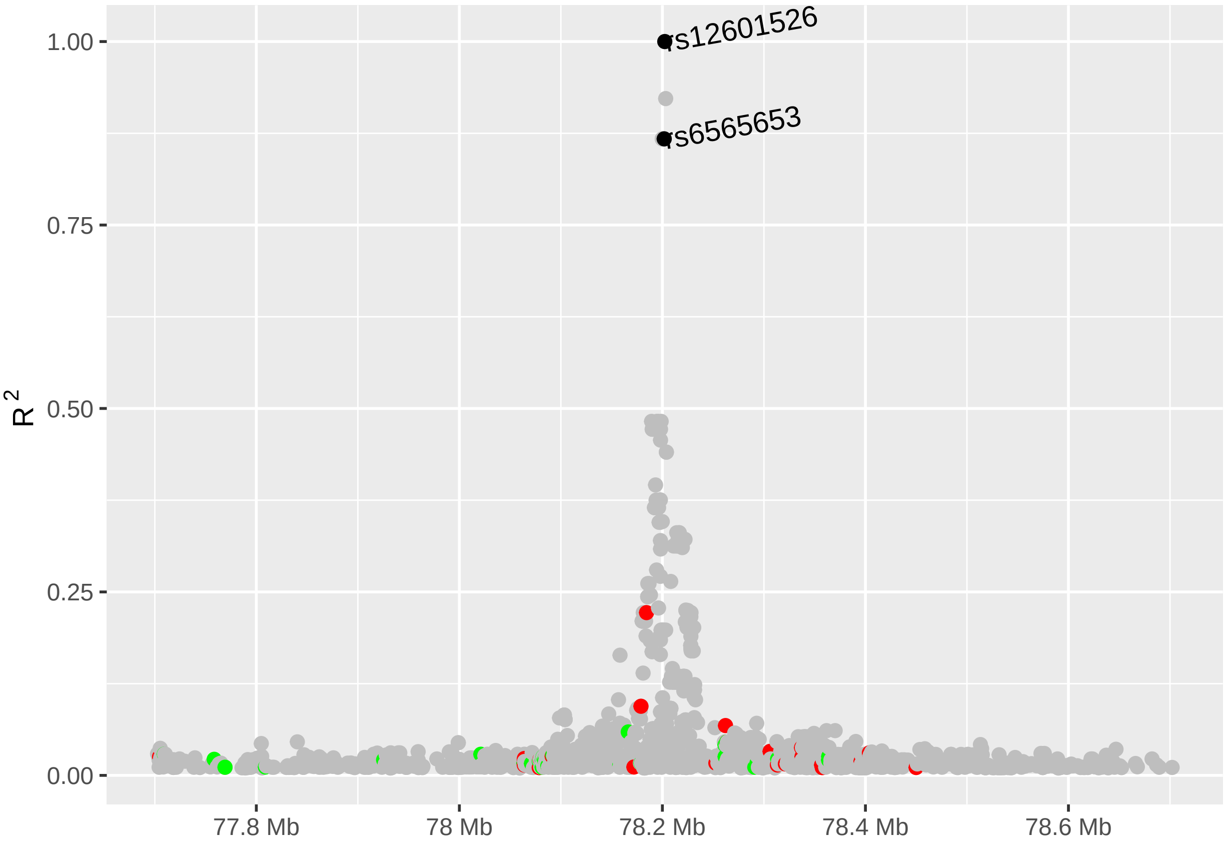


```
rs12601526_D <- ggplot(CEU_12601526_df, aes(Start,Dprime)) +
        xlim (scale_combined) +
        geom_point(size = 2, aes(colour = Function)) +
        geom_point(size = 2, data=subset(CEU_12601526_df, DisplayName == "Y"),
                   aes(Start, Dprime), color = "black") +
        geom_text(data=subset(CEU_12601526_df, DisplayName == "Y"),
                  aes(Start, Dprime,label=ID, hjust = 0, angle = 10)) +
        labs(y = "D\'", color = "SNP Function\n") +
        scale_color_manual(labels = c("missense", "unkwnown", "synonymous"),
                           values = c("red", "grey", "green")) +
        scale_x_sequnit("Mb") +
        theme(axis.title.x = element_blank (), legend.position = "none")
fixed(rs12601526_D) <- TRUE
rs12601526_D
```

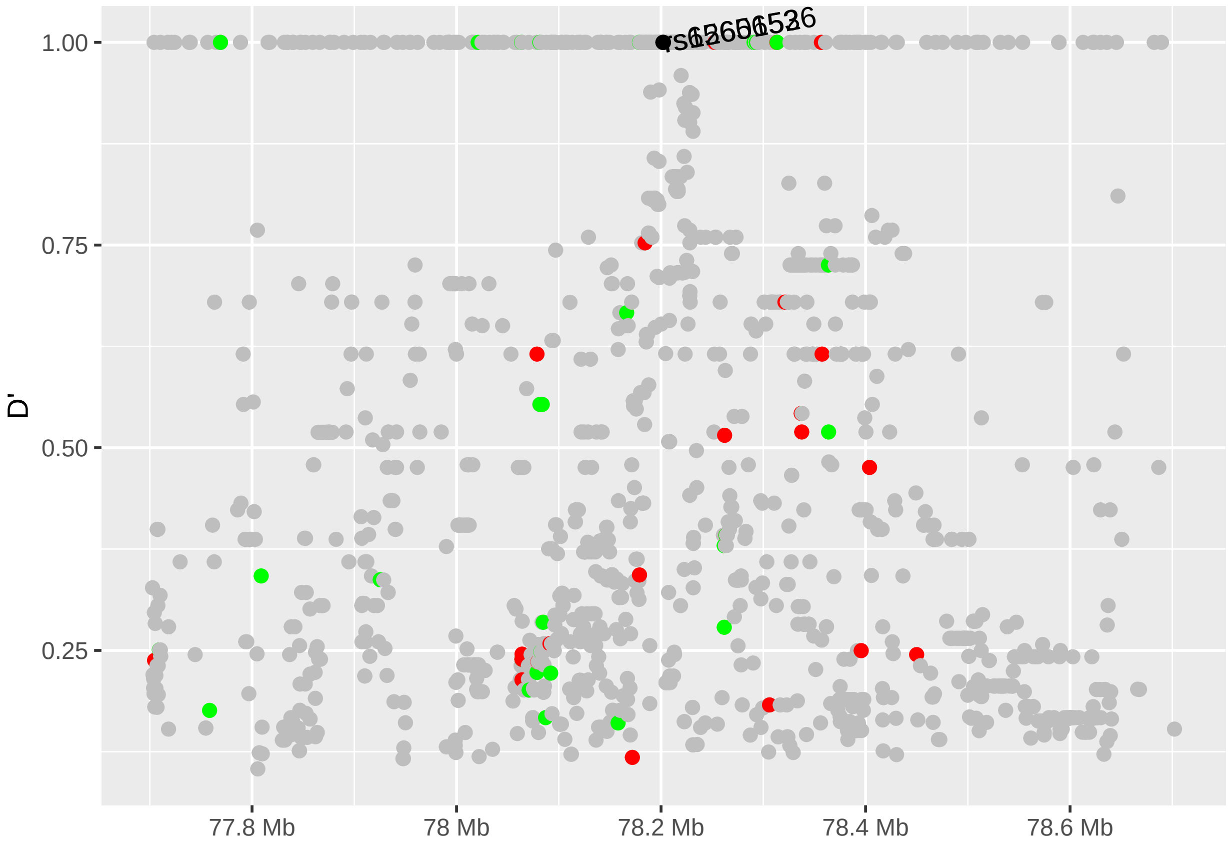


```
# Missing genes: "LOC101928766", "LOC101928738", "MIR4730"

data(genesymbol, package = "biovizBase")
wh <- genesymbol[c("ENPP7", "RPTOR")]
wh <- range(wh, ignore.strand = TRUE)

rs12601526_GENES <- autoplot(Homo.sapiens, which = wh, xlab = "Chromosome 17", ylab = "GENES",
                             label.color = "black", color = "brown",
                             fill = "brown", columns = c("ALIAS", "GO"), scale = "Mb") +
        xlim (scale_combined) +
        scale_x_sequnit("Mb")
rs12601526_GENES
```

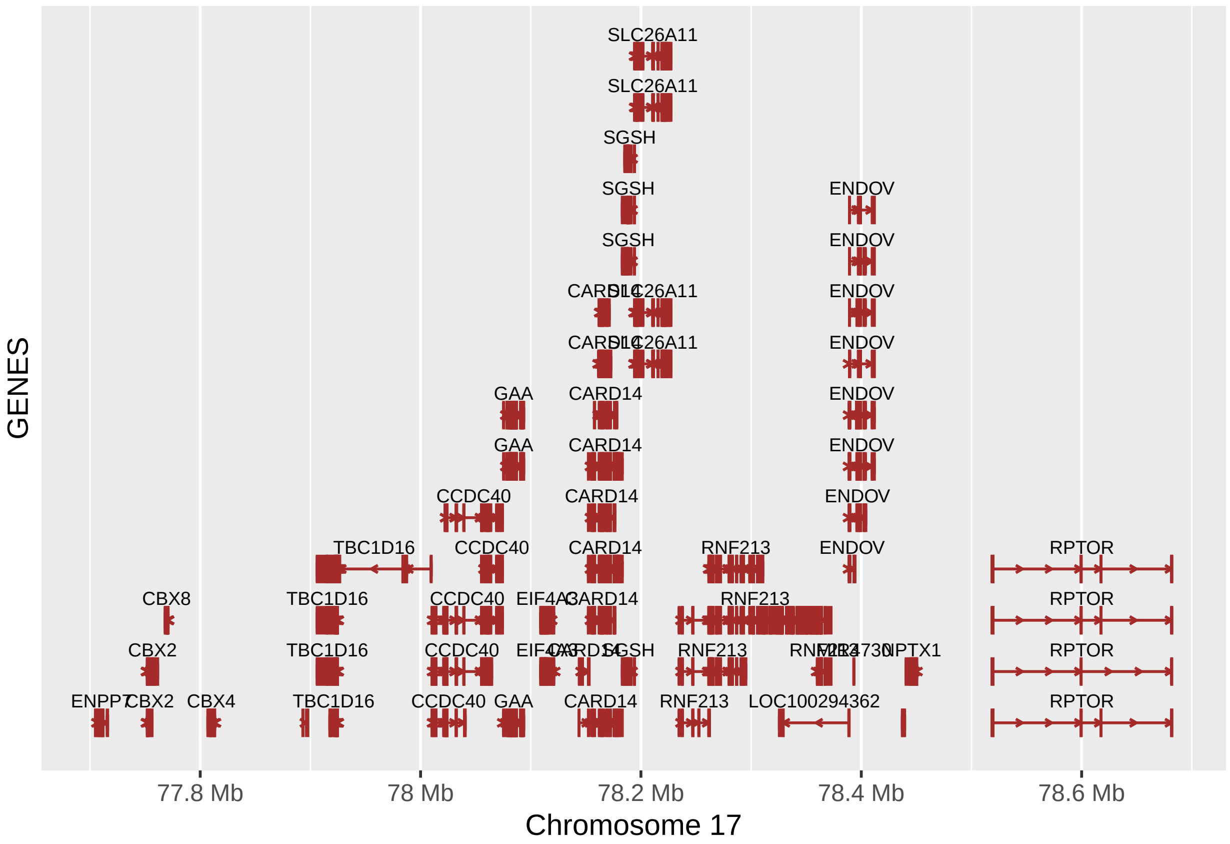


```
fixed (rs12601526_GENES) <- TRUE

rs12601526_GENES.gg <- rs12601526_GENES@ggplot
rs12601526_GENES.gg
```

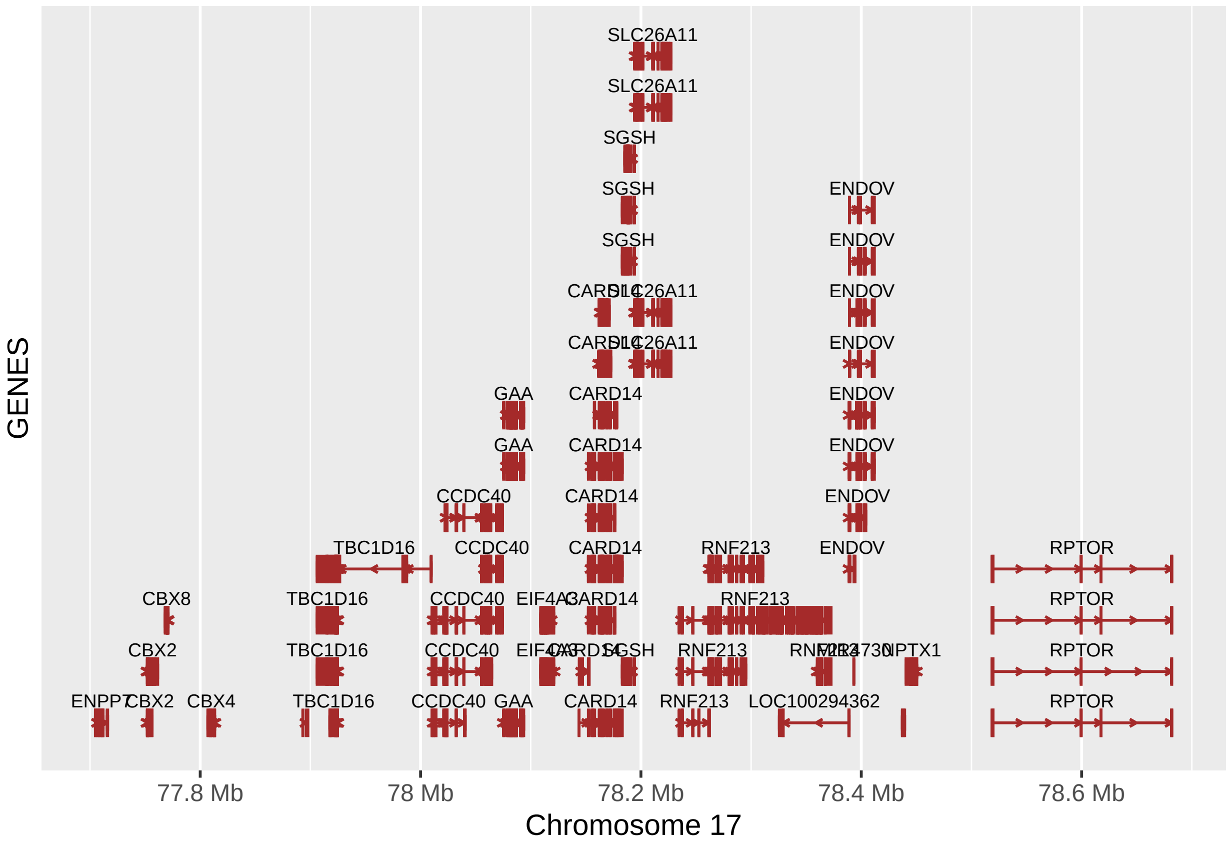


```
# Get the gtables
gA <- ggplotGrob (rs6565653_r)
gB <- ggplotGrob (rs6565653_D)
gC <- ggplotGrob (rs6565653_GENES.gg)

gD <- ggplotGrob (rs12601526_r)
gE <- ggplotGrob (rs12601526_D)
gF <- ggplotGrob (rs12601526_GENES.gg)

# Set the widths
gB$widths <- gA$widths
gC$widths <- gA$widths

gE$widths <- gD$widths
gF$widths <- gD$widths

# Arrange the three charts
rs6565653_title = textGrob("rs6565653", gp=gpar(fontsize=12, font = 2))
grid.newpage()
CEU_plot_rs6565653 <- grid.arrange(gA, gB, gC,  heights = c(4,4,6), top = rs6565653_title)
```

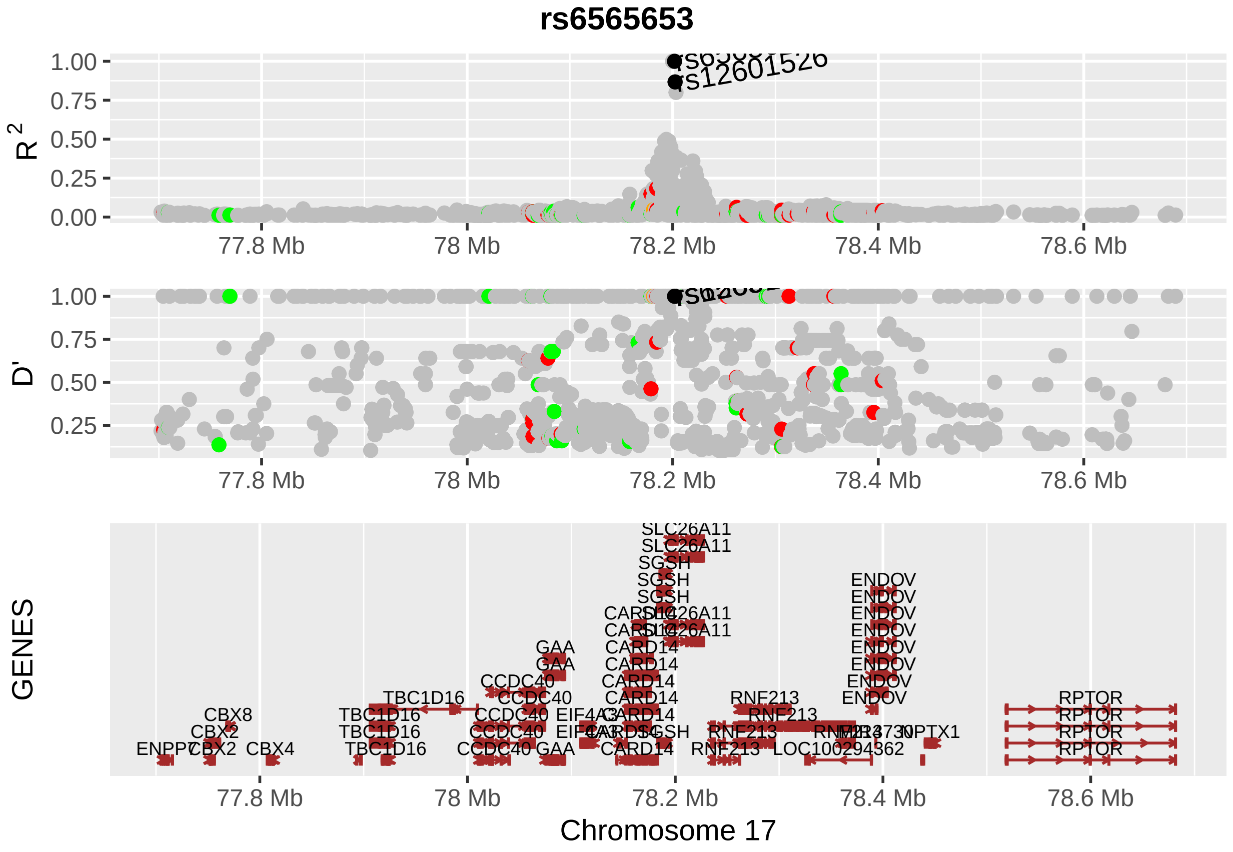


```
rs12601526_title = textGrob("rs12601526", gp=gpar(fontsize=12, font = 2))
grid.newpage()
CEU_plot_rs12601526 <- grid.arrange(gD, gE, gF, heights = c(4,4,6), top = rs12601526_title)
```

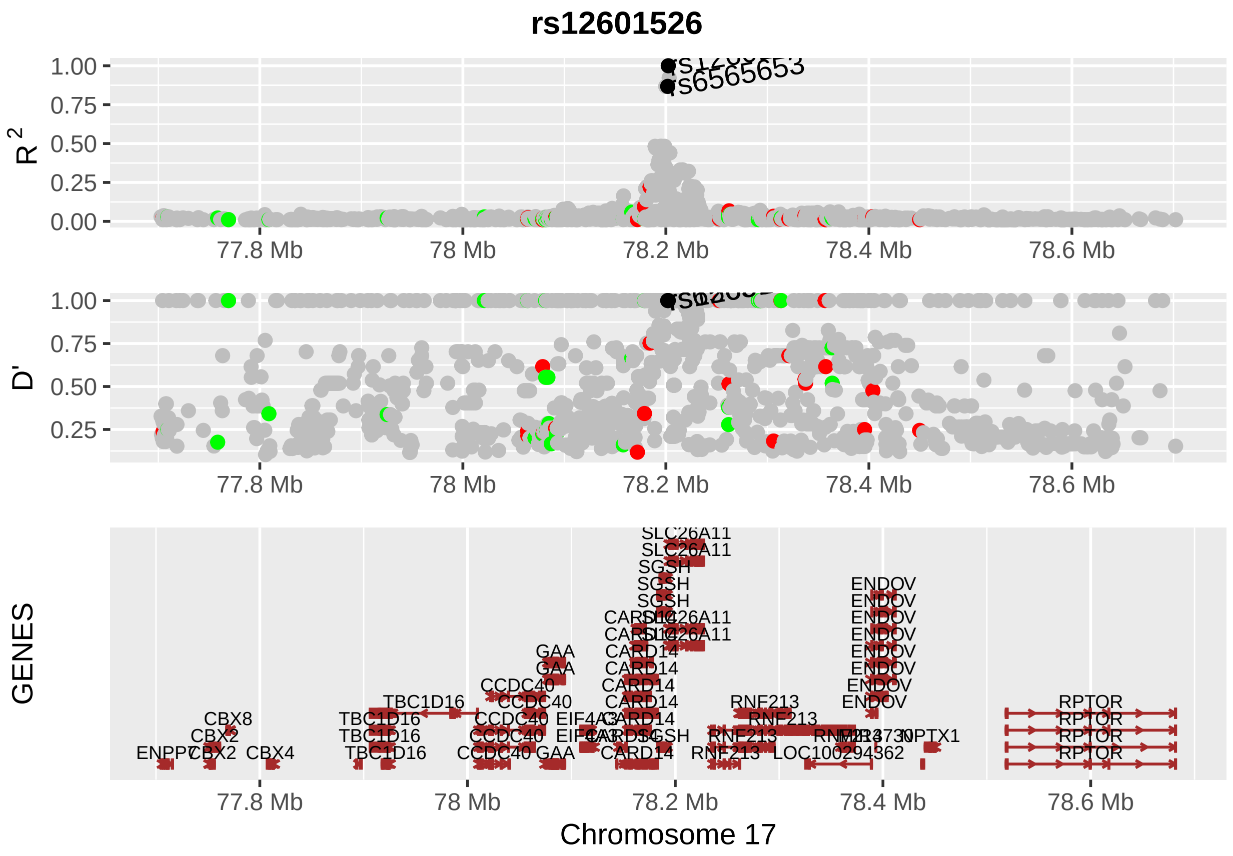


```
main_title = textGrob("rs6565653 and rs12601526 proxy SNPs in CEU population",
                      gp=gpar(fontsize=16, font = 2))
grid.newpage()
```

##### Supplemntary Figure 2 - Plot "rs6565653 and rs12601526 proxy SNPs in CEU population"

```
FINAL_PLOT_CEU <- grid.arrange(CEU_plot_rs6565653, CEU_plot_rs12601526, ncol = 2, top =
                                main_title)
```

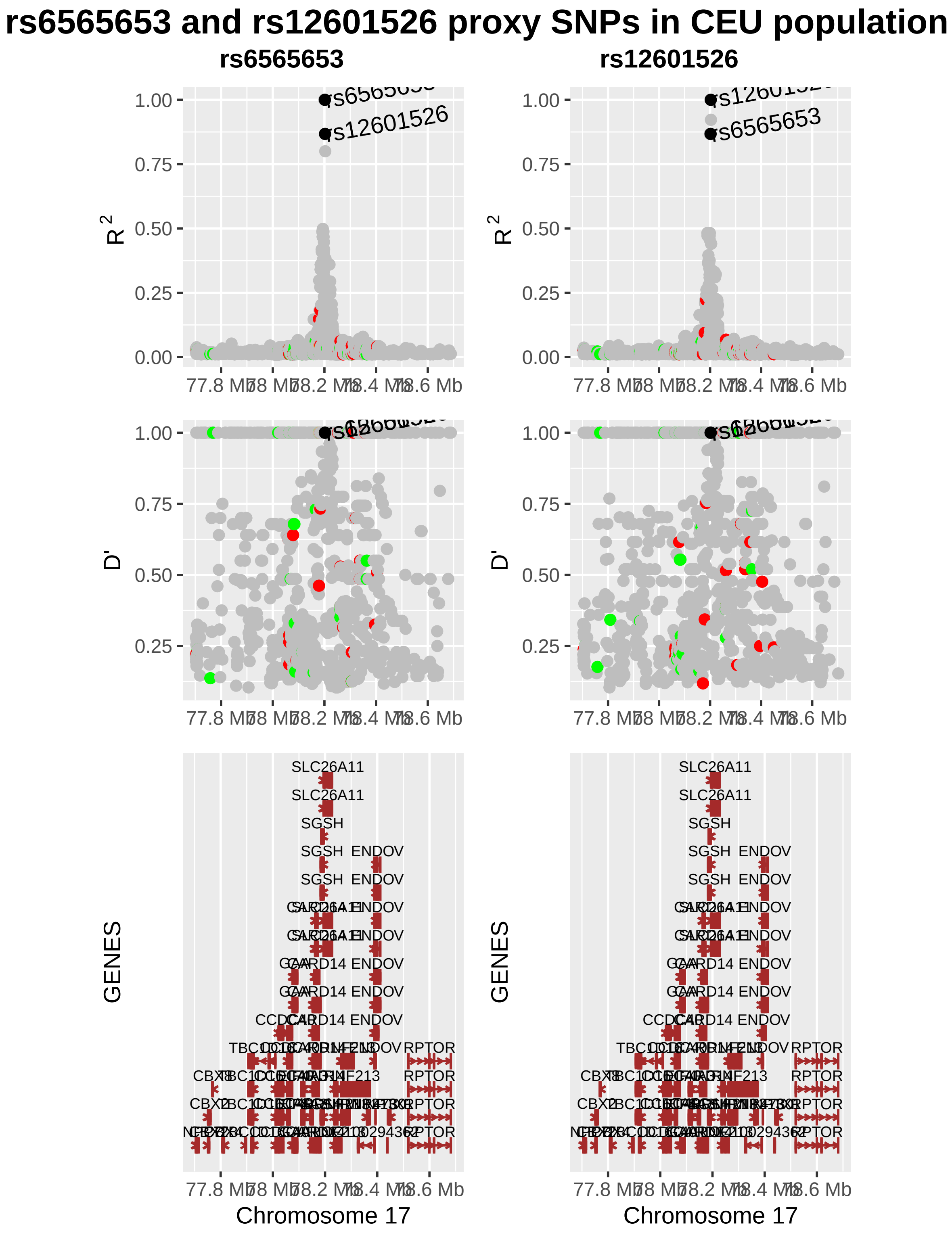


```r
dev.off()
```

```
## null device
##           1
```

#### Proxy SNPs for JPT population

Get proxy SNPs for JPT population for rs6565653 and rs12601526

```
# JPT population rs6565653 proxy SNPs
dirname <- getwd()
setwd(dirname)


# Plots for rs6565653 in JPT population
JPT_6565653_df <- as.data.frame(read.table("input_files/JPT_files/Proxy SNPs rs6565653 JPT population.tx
                                           sep = "\t", header = TRUE, na.strings = c(".", "NA"),
                                           stringsAsFactors = FALSE))

JPT_6565653_df$Function[is.na(JPT_6565653_df$Function)] <- "N/A"

range_start<-head(JPT_6565653_df$Start,n=1)
range_end<-tail(JPT_6565653_df$Start,n=1)
scale_combined <- GRanges('chr17', IRanges(start = range_start, end = range_end))


legend_title <- "SNP function"

y_lab_r <- as.expression(bquote('R'^{~2}~''))

rs6565653_r <- ggplot(JPT_6565653_df, aes(Start,R2)) +
        xlim (scale_combined) +
        geom_point(size = 2, aes(colour = Function)) +
        geom_point(size = 2, data=subset(JPT_6565653_df, DisplayName == "Y"),
                   aes(Start, R2), color = "black") +
        geom_text(data=subset(JPT_6565653_df, DisplayName == "Y"),
                  aes(Start, R2,label=ID, hjust = 0, angle = 10)) +
        labs(y = y_lab_r, color = "SNP Function\n") +
        scale_color_manual(labels = c("missense", "unkwnown", "nonsense", "synonymous"),
                           values = c("red", "grey", "orange", "green")) +
        scale_x_sequnit("Mb") +
        theme(axis.title.x = element_blank (), legend.position = "none")
fixed(rs6565653_r) <- TRUE
rs6565653_r
```

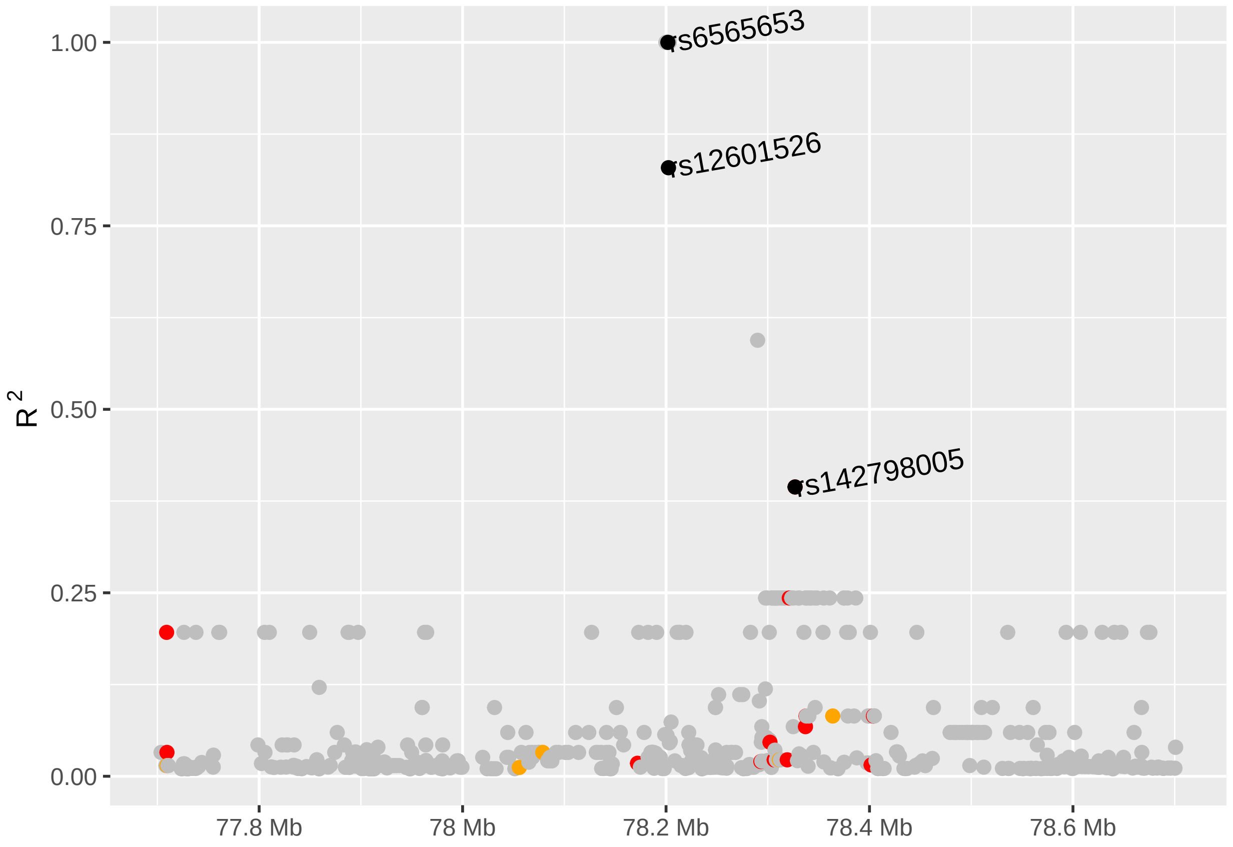


```
rs6565653_D <- ggplot(JPT_6565653_df, aes(Start,Dprime)) +
        xlim (scale_combined) +
        geom_point(size = 2, aes(colour = Function)) +
        geom_point(size = 2, data=subset(JPT_6565653_df, DisplayName == "Y"),
                   aes(Start, Dprime), color = "black") +
        geom_text(data=subset(JPT_6565653_df, DisplayName == "Y"),
                  aes(Start, Dprime,label=ID, hjust = 0, angle = 10)) +
        labs(y = "D\'", color = "SNP Function\n") +
        scale_color_manual(labels = c("missense", "unkwnown", "nonsense", "synonymous"),
                           values = c("red", "grey", "orange", "green")) +
        scale_x_sequnit("Mb")+
        theme(axis.title.x = element_blank (), legend.position = "none")
fixed(rs6565653_D) <- TRUE
rs6565653_D
```

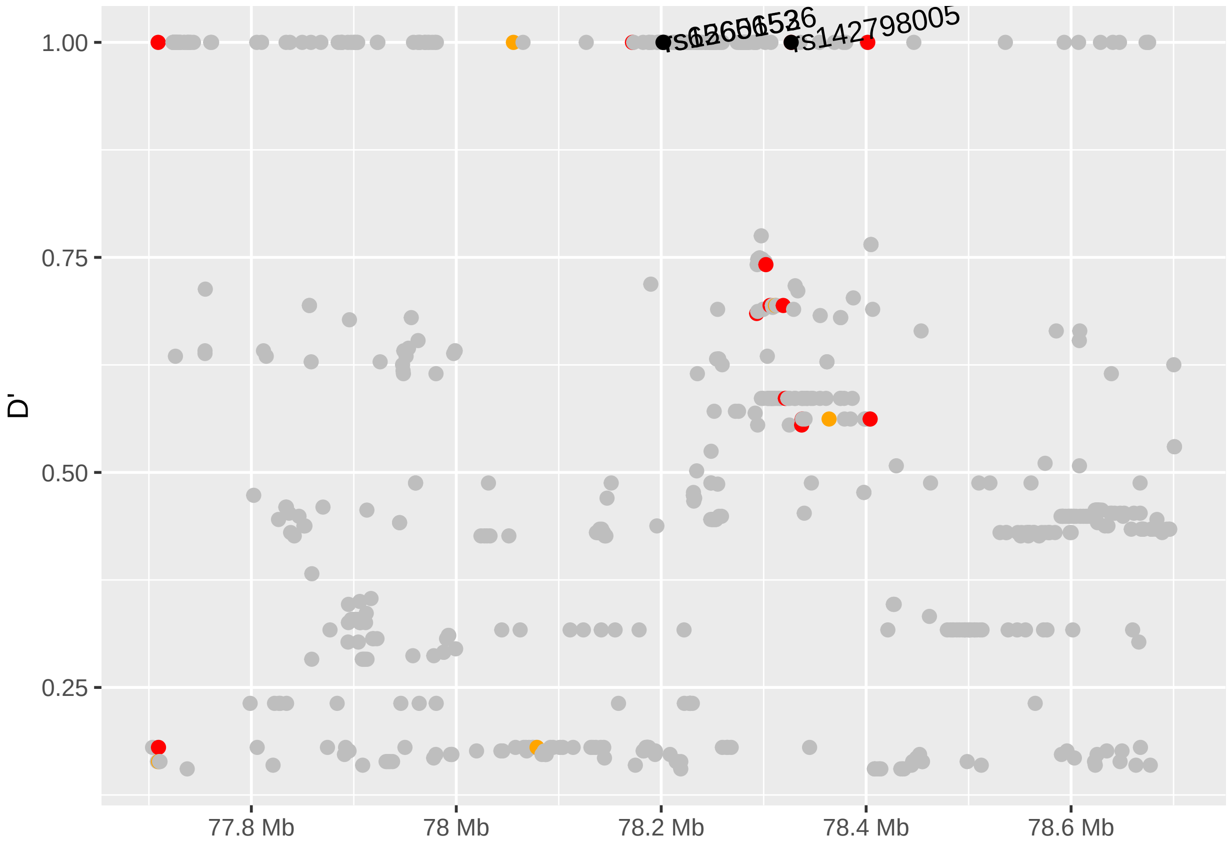


```
# Missing genes: "LOC101928766", "LOC101928738", "MIR4730"

data(genesymbol, package = "biovizBase")
wh <- genesymbol[c("ENPP7", "RPTOR")]
wh <- range(wh, ignore.strand = TRUE)

rs6565653_GENES <- autoplot(Homo.sapiens, which = wh, xlab = "Chromosome 17", ylab = "GENES",
                           label.color = "black", color = "brown", fill = "brown", columns =
                             c("ALIAS", "GO"), scale = "Mb")+
        xlim (scale_combined) +
        scale_x_sequnit("Mb")
rs6565653_GENES
```

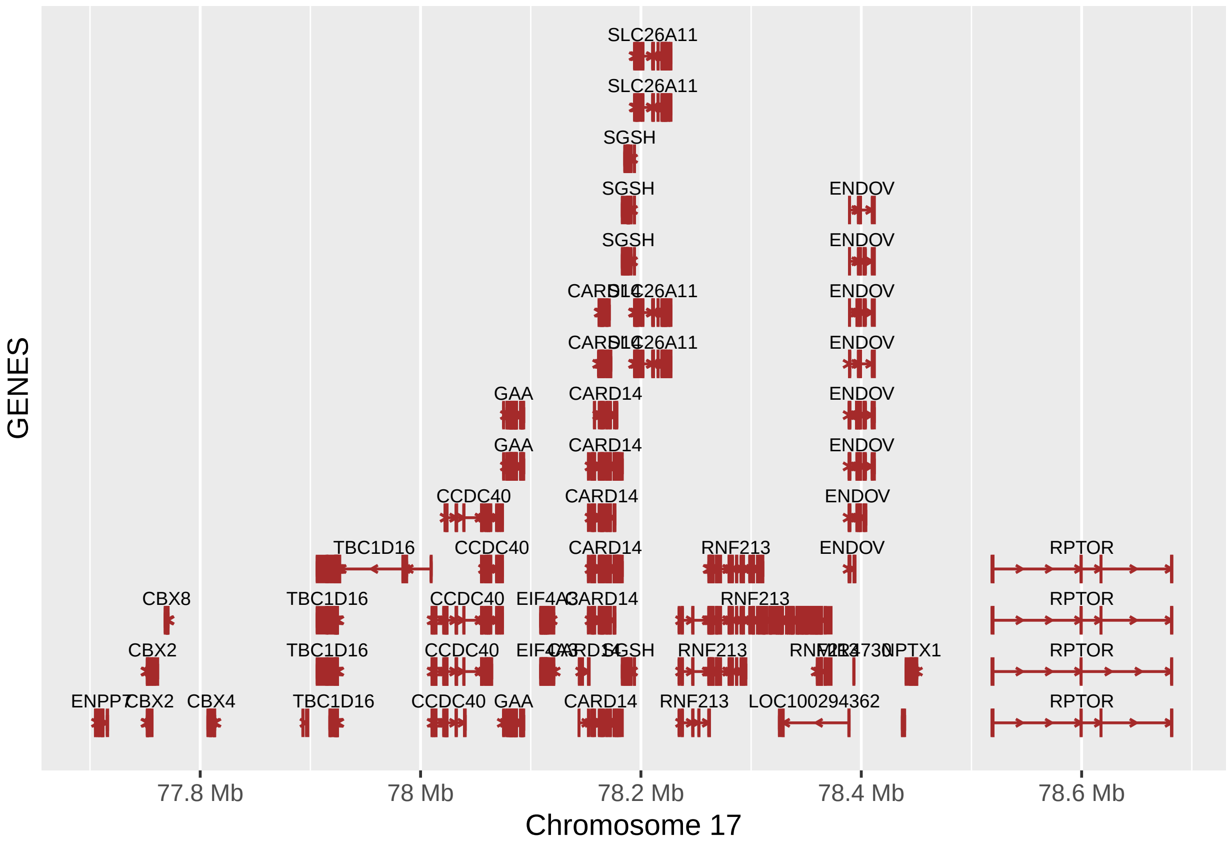


```r
fixed (rs6565653_GENES) <- TRUE

rs6565653_GENES.gg <- rs6565653_GENES@ggplot
rs6565653_GENES.gg
```

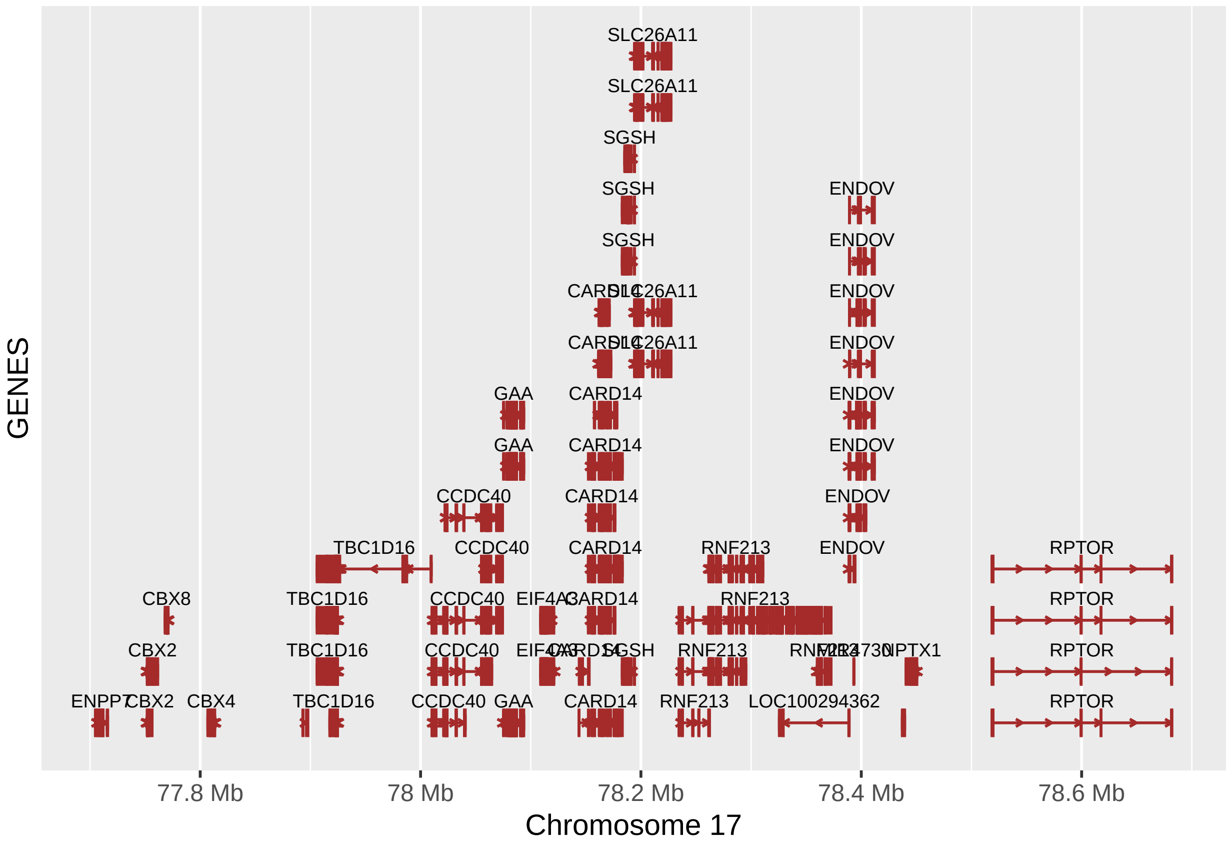


```
###############################################################

# Plots for rs12601526 in JPT population
JPT_12601526_df <- as.data.frame(read.table("input_files/JPT_files/Proxy SNPs rs12601526 JPT population.
                                           sep = "\t", header = TRUE, na.strings = c(".", "NA"),
                                           stringsAsFactors = FALSE))

JPT_12601526_df$Function[is.na(JPT_12601526_df$Function)] <- "N/A"

range_start<-head(JPT_12601526_df$Start,n=1)
range_end<-tail(JPT_12601526_df$Start,n=1)
scale_combined <- GRanges('chr17', IRanges(start = range_start, end = range_end))


legend_title <- "SNP function"

y_lab_r <- as.expression(bquote('R'^{~2}~''))

rs12601526_r <- ggplot(JPT_12601526_df, aes(Start,R2)) +
        xlim (scale_combined) +
        geom_point(size = 2, aes(colour = Function)) +
        geom_point(size = 2, data=subset(JPT_12601526_df, DisplayName == "Y"),
                   aes(Start, R2), color = "black") +
        geom_text(data=subset(JPT_12601526_df, DisplayName == "Y"),
                  aes(Start, R2,label=ID, hjust = 0, angle = 10)) +
```

```
        labs(y = y_lab_r, color = "SNP Function\n") +
        scale_color_manual(labels = c("missense", "unkwnown", "synonymous"),
                           values = c("red", "grey", "green")) +
        scale_x_sequnit("Mb") +
        theme(axis.title.x = element_blank (), legend.position = "none")
fixed(rs12601526_r) <- TRUE
rs12601526_r
```

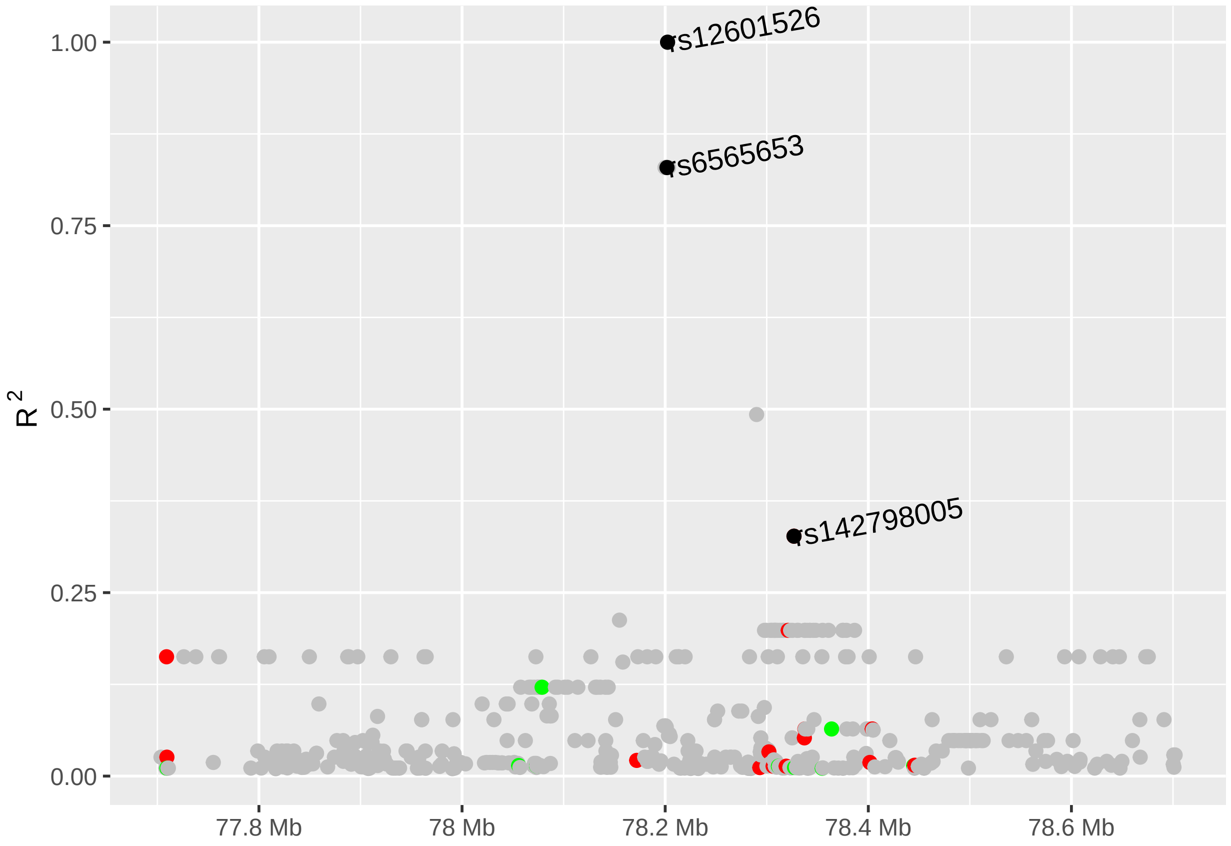


```
rs12601526_D <- ggplot(JPT_12601526_df, aes(Start,Dprime)) +
        xlim (scale_combined) +
        geom_point(size = 2, aes(colour = Function)) +
        geom_point(size = 2, data=subset(JPT_12601526_df, DisplayName == "Y"),
                   aes(Start, Dprime), color = "black") +
        geom_text(data=subset(JPT_12601526_df, DisplayName == "Y"),
                  aes(Start, Dprime,label=ID, hjust = 0, angle = 10)) +
        labs(y = "D\'", color = "SNP Function\n") +
        scale_color_manual(labels = c("missense", "unkwnown", "synonymous"),
                           values = c("red", "grey", "green")) +
        scale_x_sequnit("Mb") +
        theme(axis.title.x = element_blank (), legend.position = "none")
fixed(rs12601526_D) <- TRUE
rs12601526_D
```

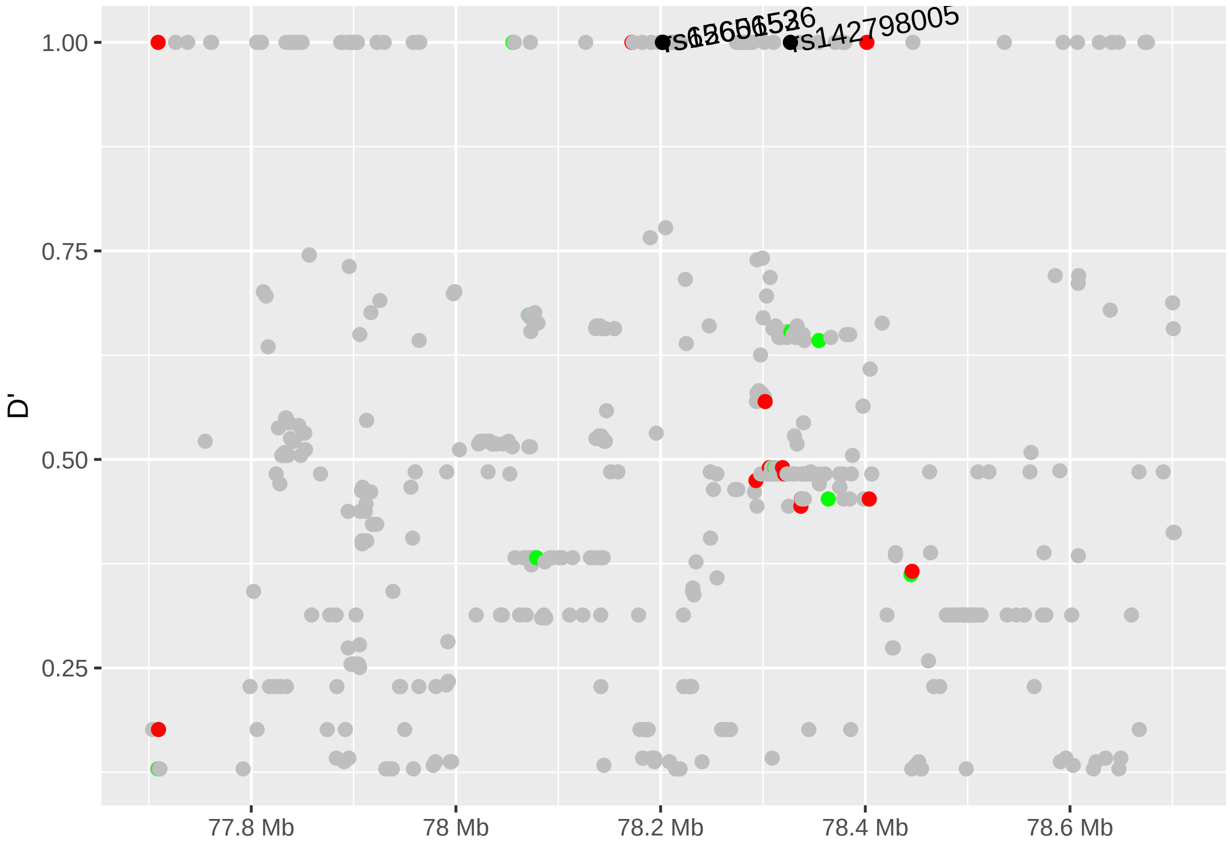


```r
# Missing genes: "LOC101928766", "LOC101928738", "MIR4730"

data(genesymbol, package = "biovizBase")
wh <- genesymbol[c("ENPP7", "RPTOR")]
wh <- range(wh, ignore.strand = TRUE)

rs12601526_GENES <- autoplot(Homo.sapiens, which = wh, xlab = "Chromosome 17", ylab = "GENES",
                             label.color = "black", color = "brown",
                             fill = "brown", columns = c("ALIAS", "GO"), scale = "Mb") +
        xlim (scale_combined) +
        scale_x_sequnit("Mb")
rs12601526_GENES
```

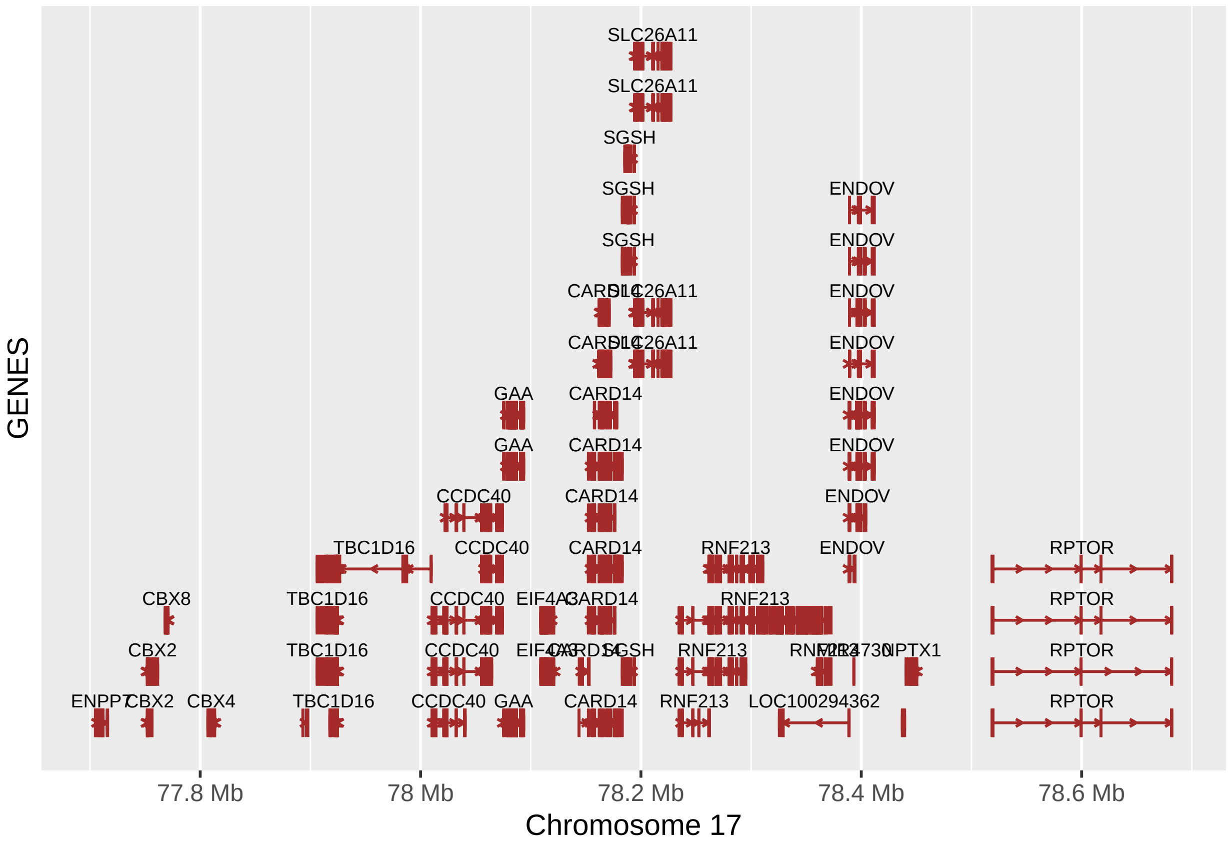


```
fixed (rs12601526_GENES) <- TRUE

rs12601526_GENES.gg <- rs12601526_GENES@ggplot
rs12601526_GENES.gg
```

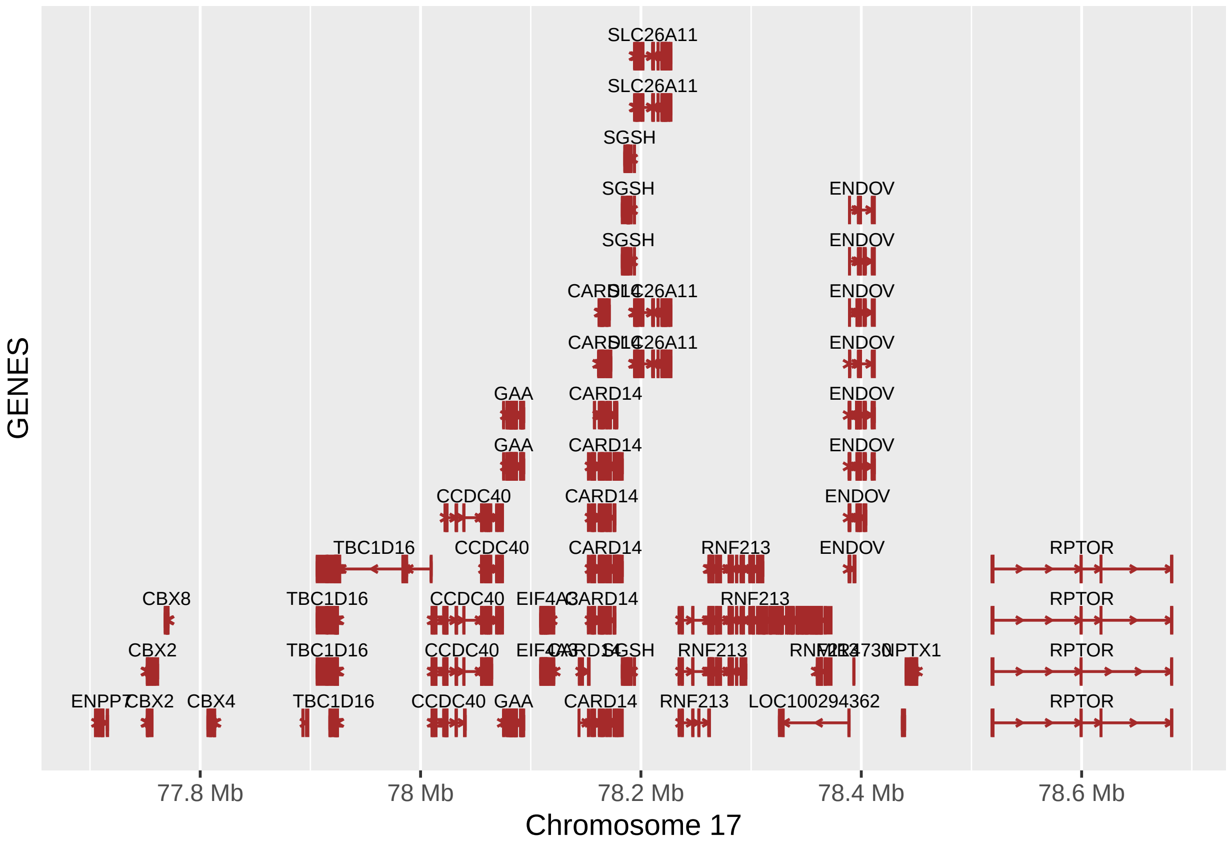


```
# Get the gtables
gA <- ggplotGrob (rs6565653_r)
gB <- ggplotGrob (rs6565653_D)
gC <- ggplotGrob (rs6565653_GENES.gg)

gD <- ggplotGrob (rs12601526_r)
gE <- ggplotGrob (rs12601526_D)
gF <- ggplotGrob (rs12601526_GENES.gg)

# Set the widths
gB$widths <- gA$widths
gC$widths <- gA$widths

gE$widths <- gD$widths
gF$widths <- gD$widths

# Arrange the three charts
rs6565653_title = textGrob("rs6565653", gp=gpar(fontsize=12, font = 2))
grid.newpage()
JPT_plot_rs6565653 <- grid.arrange(gA, gB, gC,  heights = c(4,4,6), top = rs6565653_title)
```

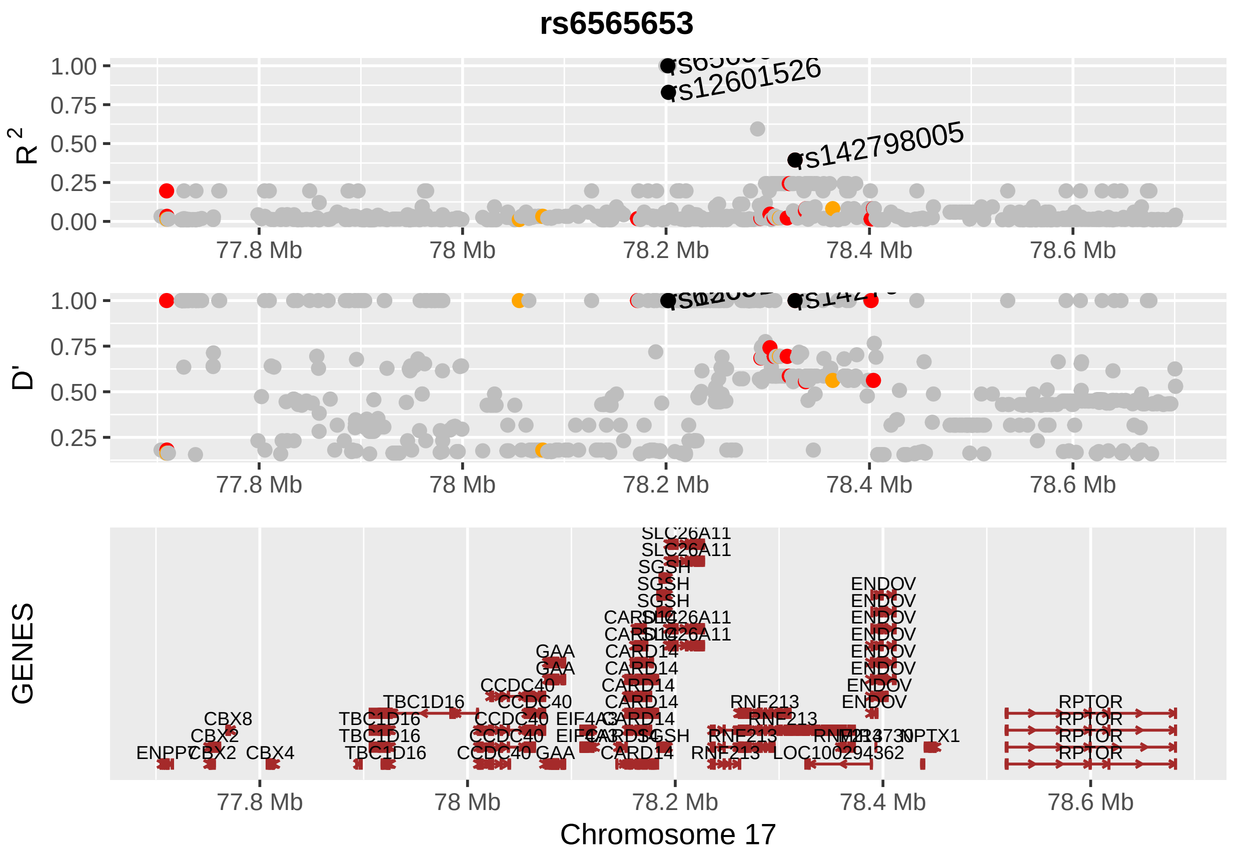


```
rs12601526_title = textGrob("rs12601526", gp=gpar(fontsize=12, font = 2))
grid.newpage()
JPT_plot_rs12601526 <- grid.arrange(gD, gE, gF, heights = c(4,4,6), top = rs12601526_title)
```

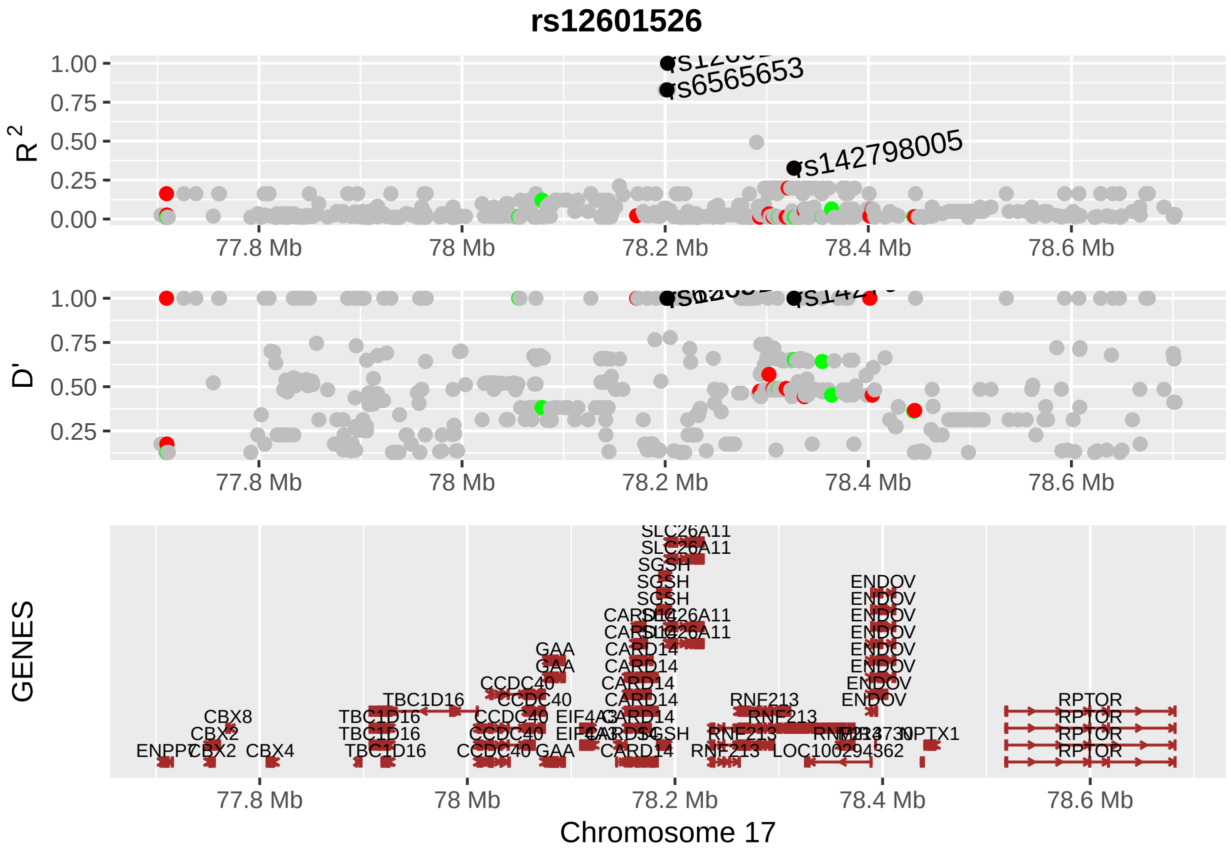


```
main_title = textGrob("rs6565653 and rs12601526 proxy SNPs in JPT population",
                      gp=gpar(fontsize=16, font = 2))
grid.newpage()
```

#### Supplementary Figure 3 - Plot "rs6565653 and rs12601526 proxy SNPs in JPT population"

```
FINAL_PLOT_JPT <- grid.arrange(JPT_plot_rs6565653, JPT_plot_rs12601526, ncol = 2, top =
                               main_title)
```

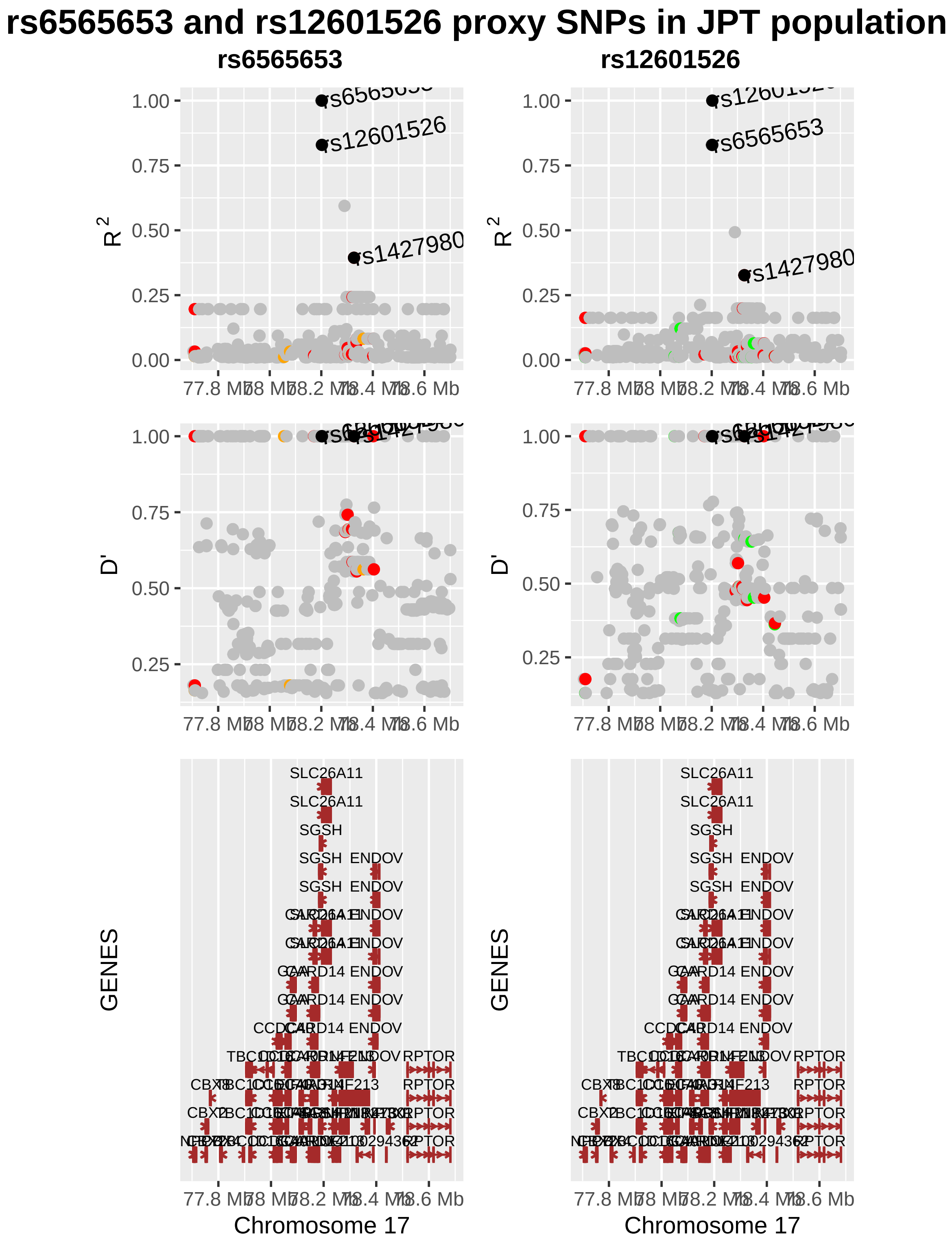


```r
dev.off()
```

```
## null device
##           1
```

#### Artery_eQTL_values_RNF213_SLC26A1

```
data(hg19IdeogramCyto, package = "biovizBase")
data(hg19Ideogram, package = "biovizBase")
data(genesymbol, package = "biovizBase")

hg19 <- keepSeqlevels(hg19IdeogramCyto, paste0("chr", c(1:22, "X", "Y")))

# RNF213_SLC26A11_eQTL_association_files
x_lab_aorta <- "GTEx - \"Artery Aorta\""
x_lab_coronary <- "GTEx - \"Artery Coronary\""
x_lab_tibial <- "GTEx - \"Artery Tibial\""
y_lab <- as.expression(bquote('-log10 ('~italic(p)~'-value)'), fontsize=8, font = 2)


# Plots for SLC26A11 significant eQTL SNPs
# SLC26A11 Genes Plot
wh_SLC26A11 <- genesymbol[c("SLC26A11")]
SLC26A11_start <- as.vector(slot(wh_SLC26A11@ranges, "start") - 25000)
SLC26A11_end <- as.vector(slot(wh_SLC26A11@ranges, "start") + slot (wh_SLC26A11@ranges,
                                                                    "width") - 1 + 25000)
SLC26A11_range <- GRanges('chr17', IRanges(start = SLC26A11_start, end = SLC26A11_end))
wh_SLC26A11 <- range(SLC26A11_range, ignore.strand = TRUE)


SLC26A11_GENES <- autoplot(Homo.sapiens, which = wh_SLC26A11, xlab = "Chromosome 17", ylab =
                              "GENES", label.color = "black", color = "brown", fill = "brown",
                            columns = c("ALIAS", "GO"), scale = "Mb")+
        xlim (SLC26A11_range) +
        scale_x_sequnit("Mb") +
        theme(axis.text.x = element_text(size=8), axis.text.y = element_text(size=10),
              axis.title.x = element_text(size=14), axis.title.y = element_text(size=12))
SLC26A11_GENES
```

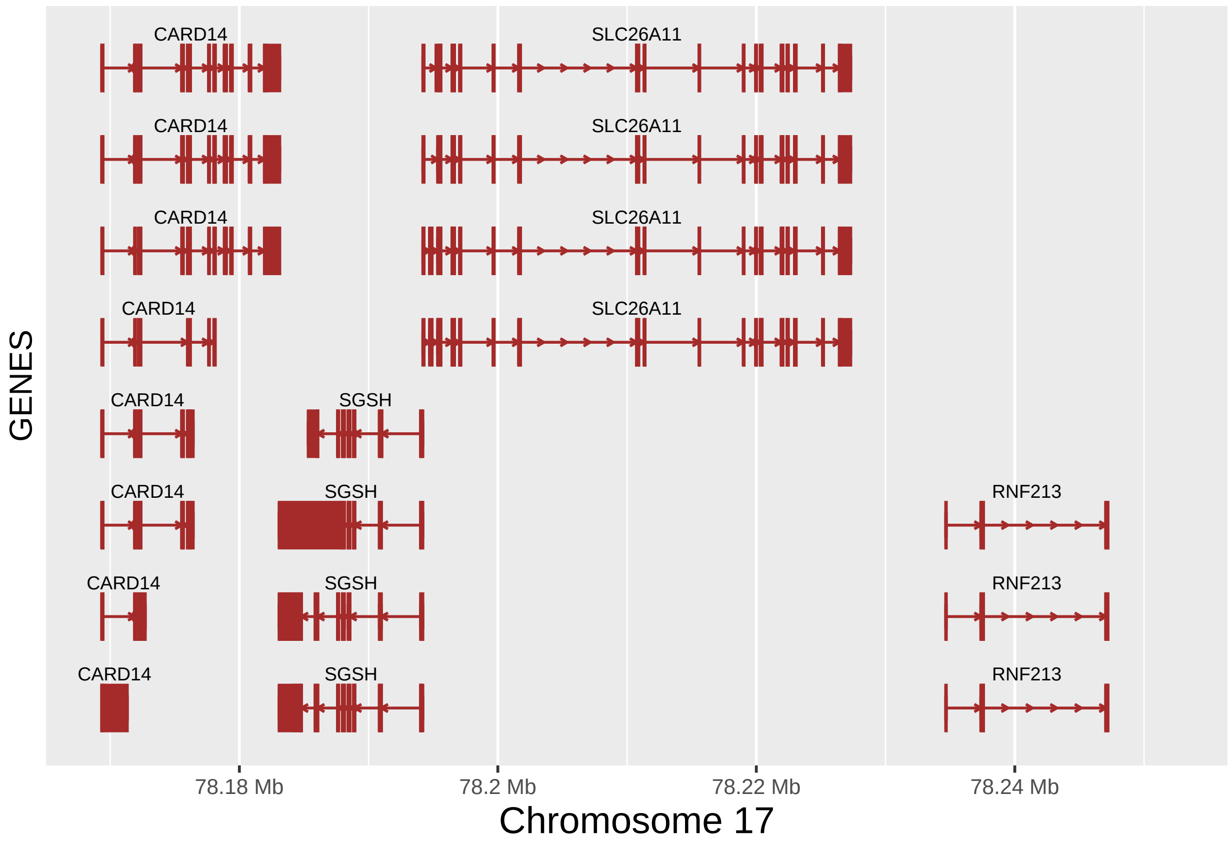


```
fixed (SLC26A11_GENES) <- TRUE

SLC26A11_GENES.gg <- SLC26A11_GENES@ggplot
SLC26A11_GENES.gg
```

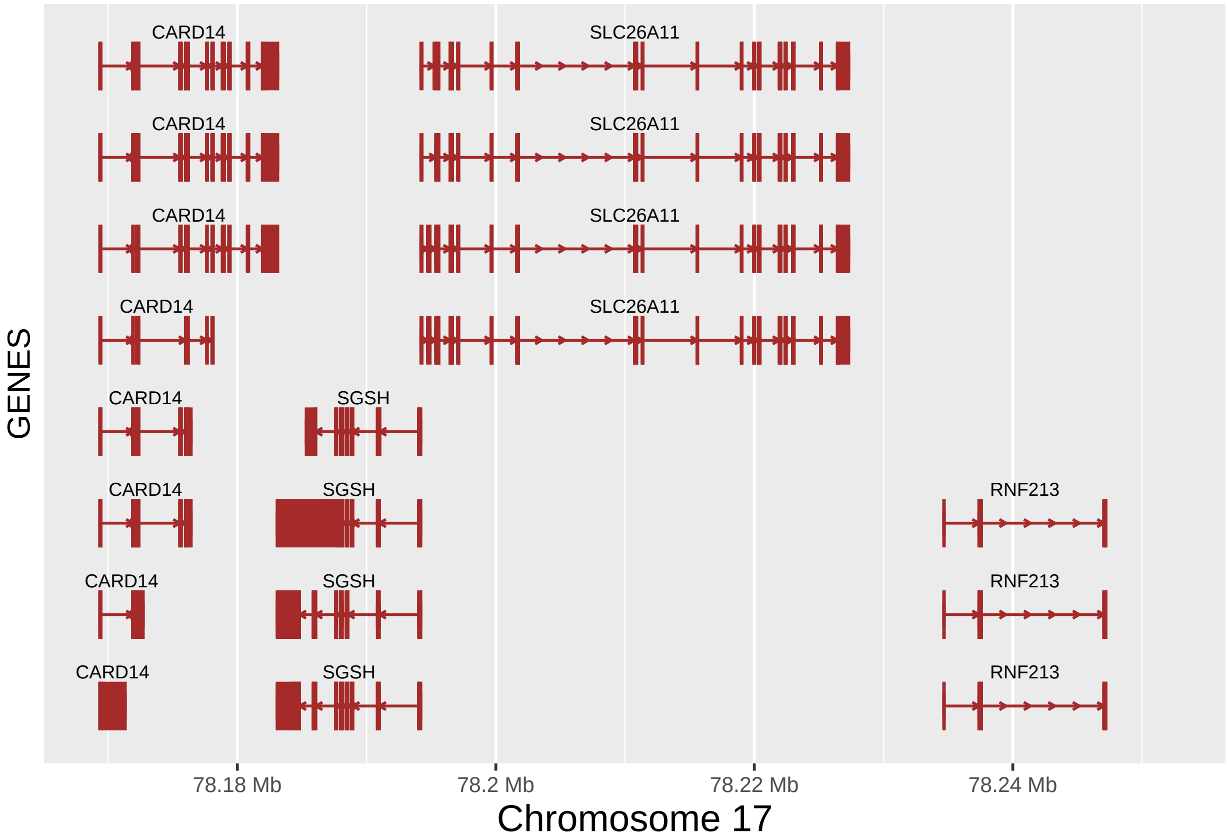


```r
# AORTA
SLC26A11_aorta_df <- as.data.frame(read.table("input_files/GTEx_files/SLC26A11_aorta.txt",
                                              sep = "\t", header = TRUE,
                                              na.strings = c(".", "NA"),
                                              stringsAsFactors = FALSE))

SLC26A11_aorta <- ggplot(SLC26A11_aorta_df, aes(x = position, y = -log10(pval_nominal))) +
        xlim (SLC26A11_range) +
        scale_y_continuous(limits = c(4,24), breaks=seq(4,24,4), expand = c(0,0)) +
        geom_point(size = 2.25, data=subset(SLC26A11_aorta_df,
                                            -log10(SLC26A11_aorta_df$pval_nominal)>4),
                   aes(position, -log10(pval_nominal)), color = "black") +
        geom_point(size = 1.25, data=subset(SLC26A11_aorta_df,
                                            -log10(SLC26A11_aorta_df$pval_nominal)>4),
                   aes(position, -log10(pval_nominal)), color = "red") +
        geom_text(data=subset(SLC26A11_aorta_df, Include == "Y"),
                  aes(position, -log10(pval_nominal),label=SNP_ID, hjust = 0, angle = 10)) +
        scale_x_sequnit("Mb") +
        labs(x = x_lab_aorta, y = y_lab) +
        theme(axis.text.x = element_text(size=8), axis.text.y = element_text(size=10),
              axis.title.x = element_text(size=14), axis.title.y = element_text(size=12))
fixed(SLC26A11_aorta) <- TRUE
SLC26A11_aorta
```

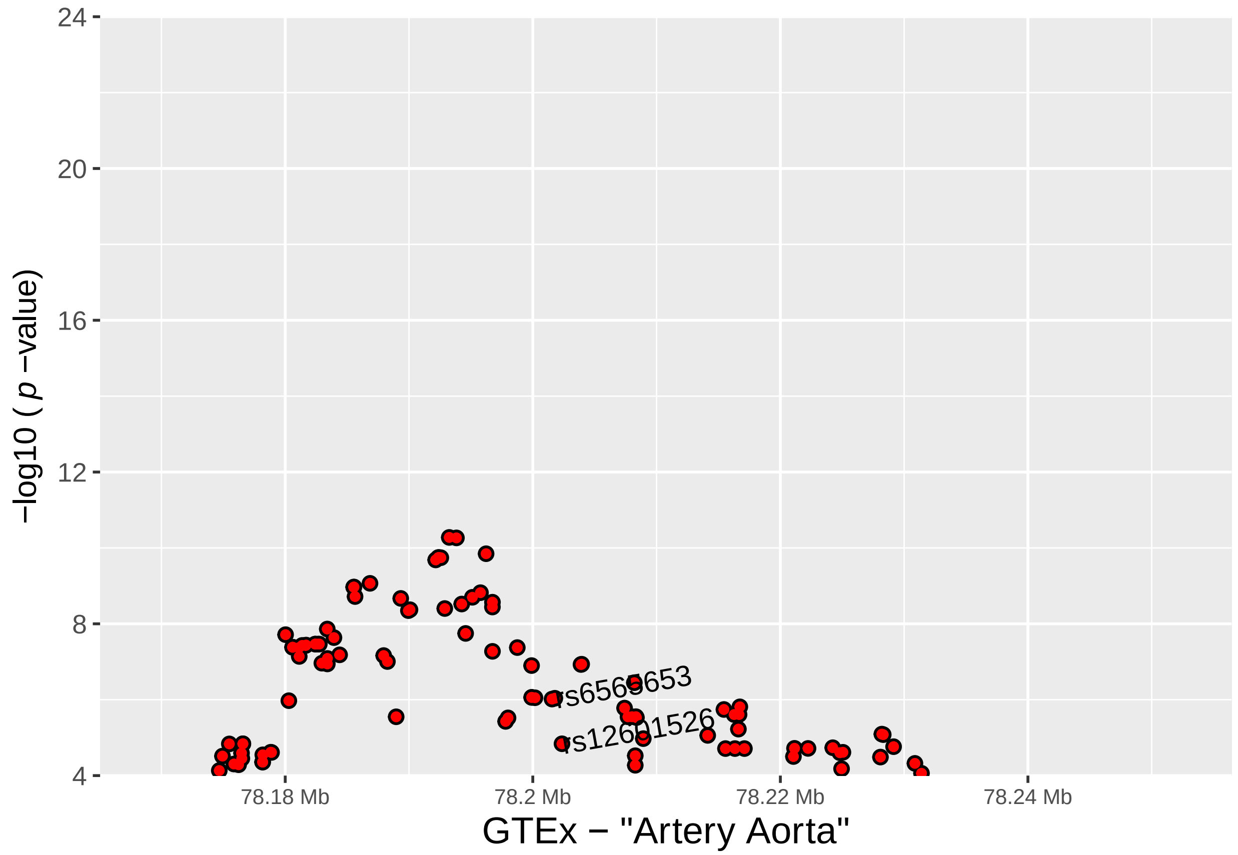


```r
# CORONARY
SLC26A11_coronary_df <- as.data.frame(read.table("input_files/GTEx_files/SLC26A11_coronary.txt",
                                                 sep = "\t", header = TRUE,
                                                 na.strings = c(".", "NA"),
                                                 stringsAsFactors = FALSE))

SLC26A11_coronary <- ggplot(SLC26A11_coronary_df, aes(x = position, y = -log10(pval_nominal))) +
        xlim (SLC26A11_range) +
        scale_y_continuous(limits = c(4,24), breaks=seq(4,24,4), expand = c(0,0)) +
        geom_point(size = 2.25, data=subset(SLC26A11_coronary_df,
                                            -log10(SLC26A11_coronary_df$pval_nominal)>4),
                   aes(position, -log10(pval_nominal)), color = "black") +
        geom_point(size = 1.25, data=subset(SLC26A11_coronary_df,
                                            -log10(SLC26A11_coronary_df$pval_nominal)>4),
                   aes(position, -log10(pval_nominal)), color = "red") +
        ##geom_text(data=subset(SLC26A11_coronary_df, Include == "Y"),
        ##           aes(position, -log10(pval_nominal),label=SNP_ID, hjust = 0, angle = 10)) +
        scale_x_sequnit("Mb") +
        labs(x = x_lab_coronary, y = y_lab) +
        theme(axis.text.x = element_text(size=8), axis.text.y = element_text(size=10),
              axis.title.x = element_text(size=14), axis.title.y = element_text(size=12))
fixed(SLC26A11_coronary) <- TRUE
SLC26A11_coronary
```

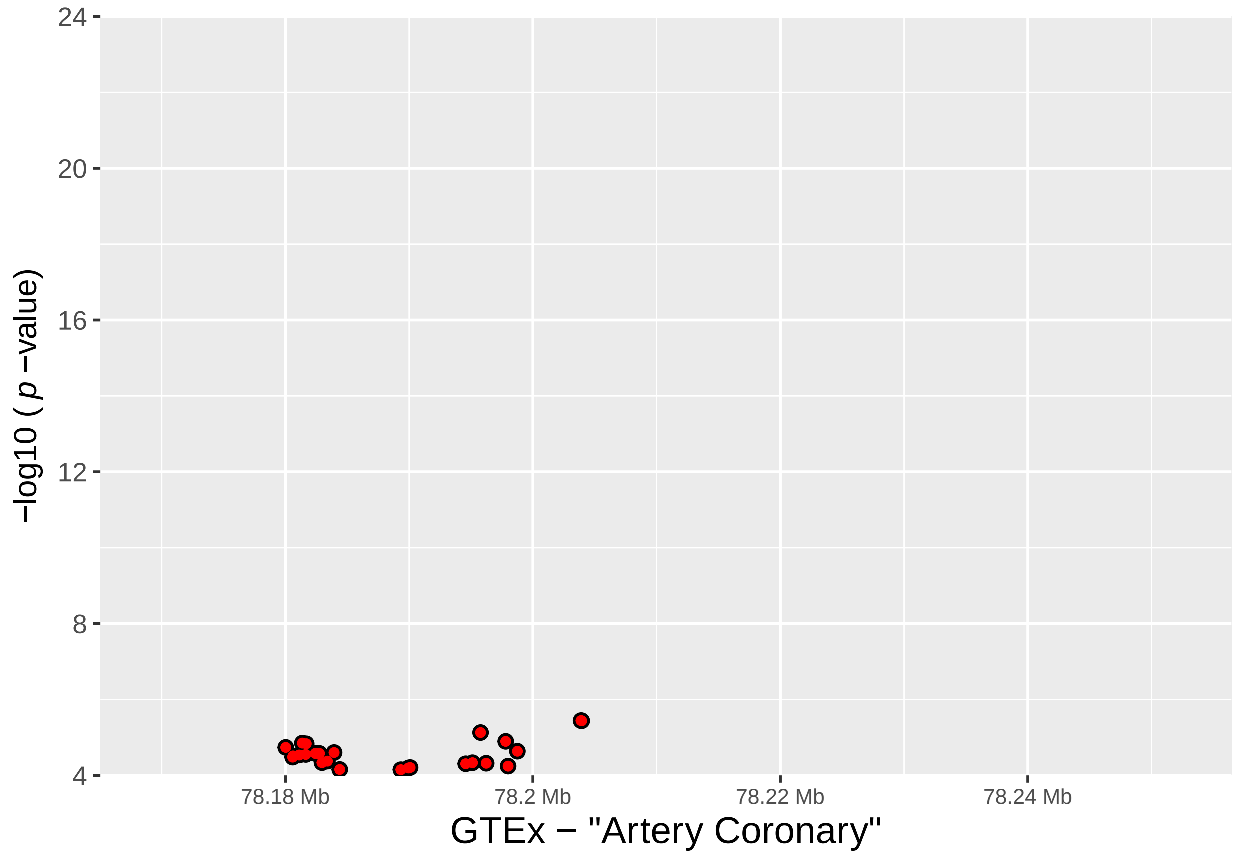


```r
# TIBIAL
SLC26A11_tibial_df <- as.data.frame(read.table("input_files/GTEx_files/SLC26A11_tibial.txt",
                                               sep = "\t", header = TRUE,
                                               na.strings = c(".", "NA"),
                                               stringsAsFactors = FALSE))

SLC26A11_tibial <- ggplot(SLC26A11_tibial_df, aes(x = position, y = -log10(pval_nominal))) +
        xlim (SLC26A11_range) +
        scale_y_continuous(limits = c(4,24), breaks=seq(4,24,4), expand = c(0,0)) +
        geom_point(size = 2.25, data=subset(SLC26A11_tibial_df,
                                            -log10(SLC26A11_tibial_df$pval_nominal)>4),
                   aes(position, -log10(pval_nominal)), color = "black") +
        geom_point(size = 1.25, data=subset(SLC26A11_tibial_df,
                                            -log10(SLC26A11_tibial_df$pval_nominal)>4),
                   aes(position, -log10(pval_nominal)), color = "red") +
        geom_text(data=subset(SLC26A11_tibial_df, Include == "Y"),
                  aes(position, -log10(pval_nominal),label=SNP_ID, hjust = 0, angle = 10)) +
        scale_x_sequnit("Mb") +
        labs(x = x_lab_tibial, y = y_lab) +
        theme(axis.text.x = element_text(size=8), axis.text.y = element_text(size=10),
              axis.title.x = element_text(size=14), axis.title.y = element_text(size=12))
fixed(SLC26A11_tibial) <- TRUE
SLC26A11_tibial
```

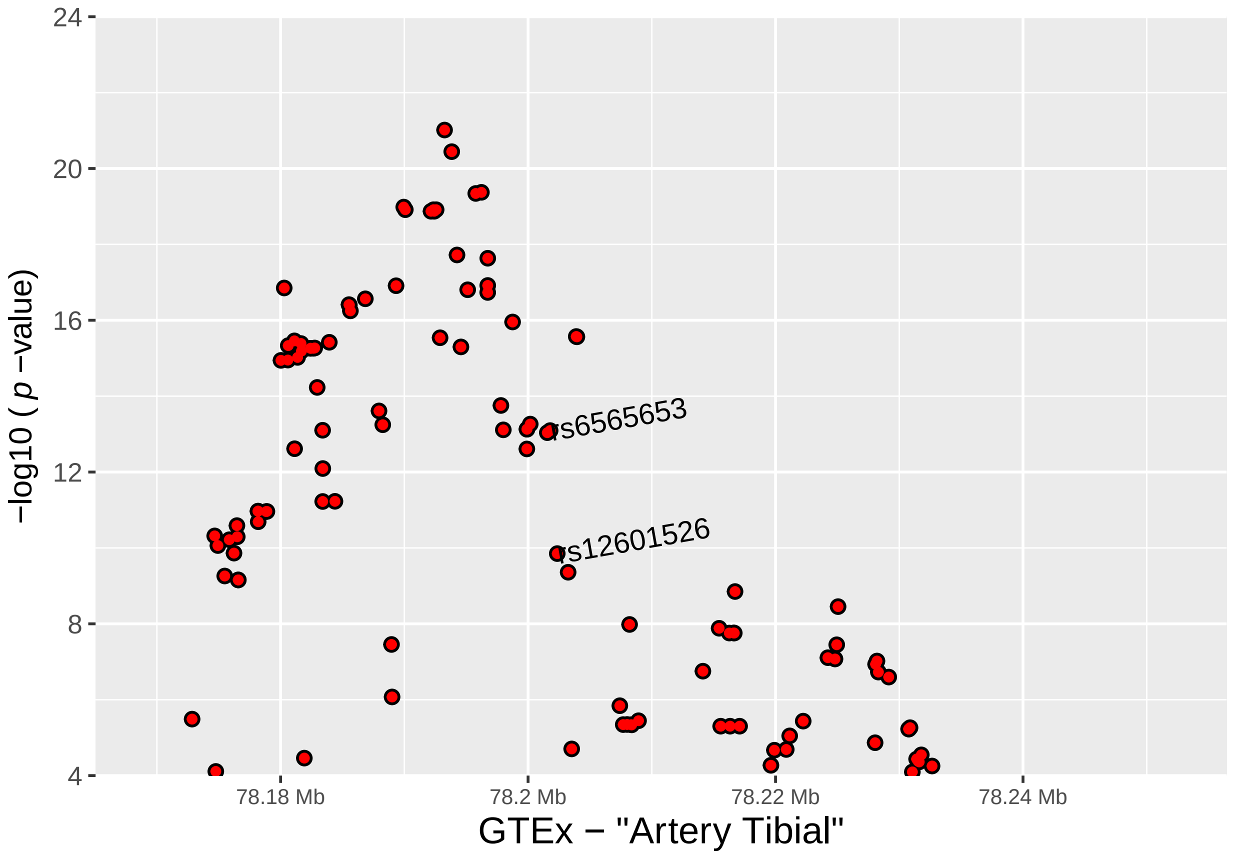


```r
# Plots for RNF213 significant eQTL SNPs
# RNF213 Genes Plot
wh_RNF213 <- genesymbol[c("RNF213")]
RNF213_start <- as.vector(slot(wh_RNF213@ranges, "start") - 25000)
RNF213_end <- as.vector(slot(wh_RNF213@ranges, "start") +
                          slot (wh_RNF213@ranges, "width") - 1 + 25000)
RNF213_range <- GRanges('chr17', IRanges(start = RNF213_start, end = RNF213_end))
wh_RNF213 <- range(RNF213_range, ignore.strand = TRUE)


RNF213_GENES <- autoplot(Homo.sapiens, which = wh_RNF213,
                         xlab = "Chromosome 17", ylab = "GENES",
                         label.color = "black", color = "brown", fill = "brown",
                         columns = c("ALIAS", "GO"), scale = "Mb") +
        xlim (RNF213_range) +
        scale_x_sequnit("Mb") +
        theme(axis.text.x = element_text(size=8), axis.text.y = element_text(size=10),
              axis.title.x = element_text(size=14), axis.title.y = element_text(size=12))
RNF213_GENES
```

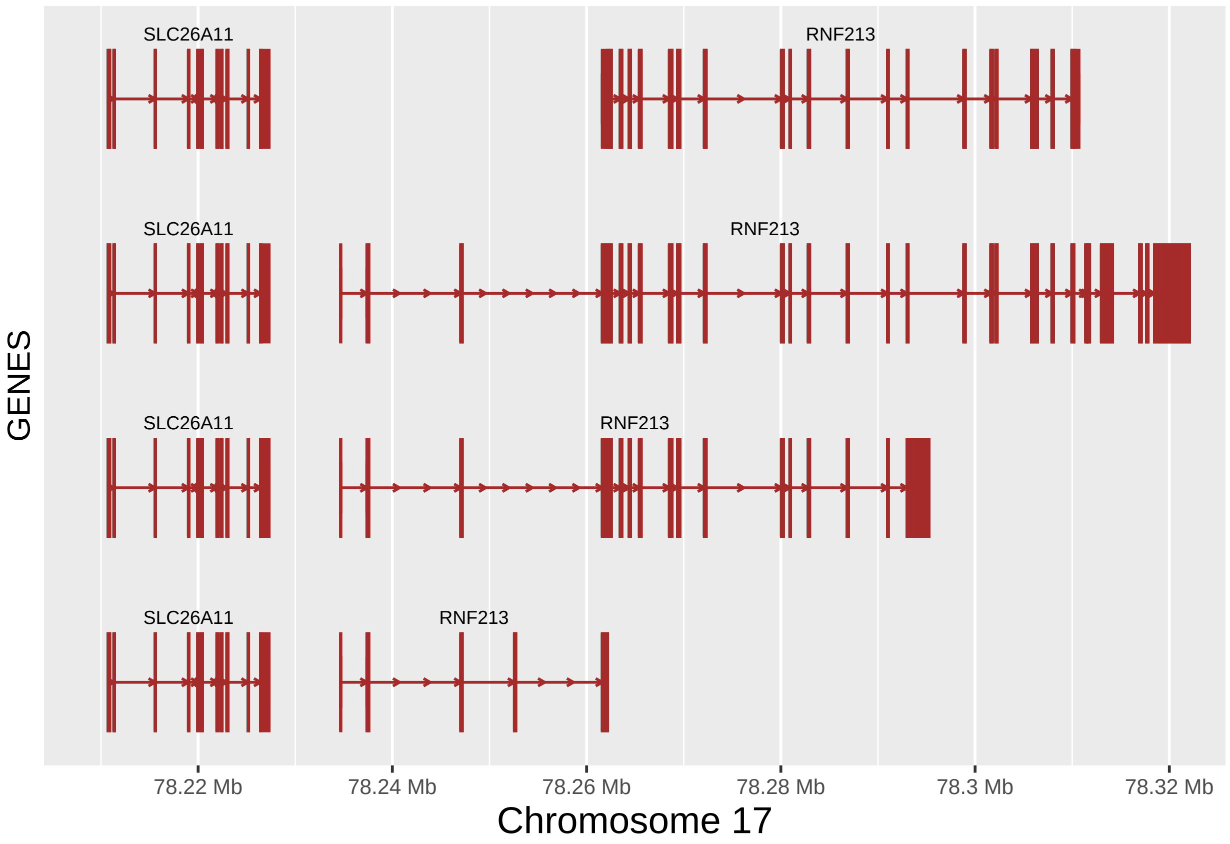


```
fixed (RNF213_GENES) <- TRUE

RNF213_GENES.gg <- RNF213_GENES@ggplot
RNF213_GENES.gg
```

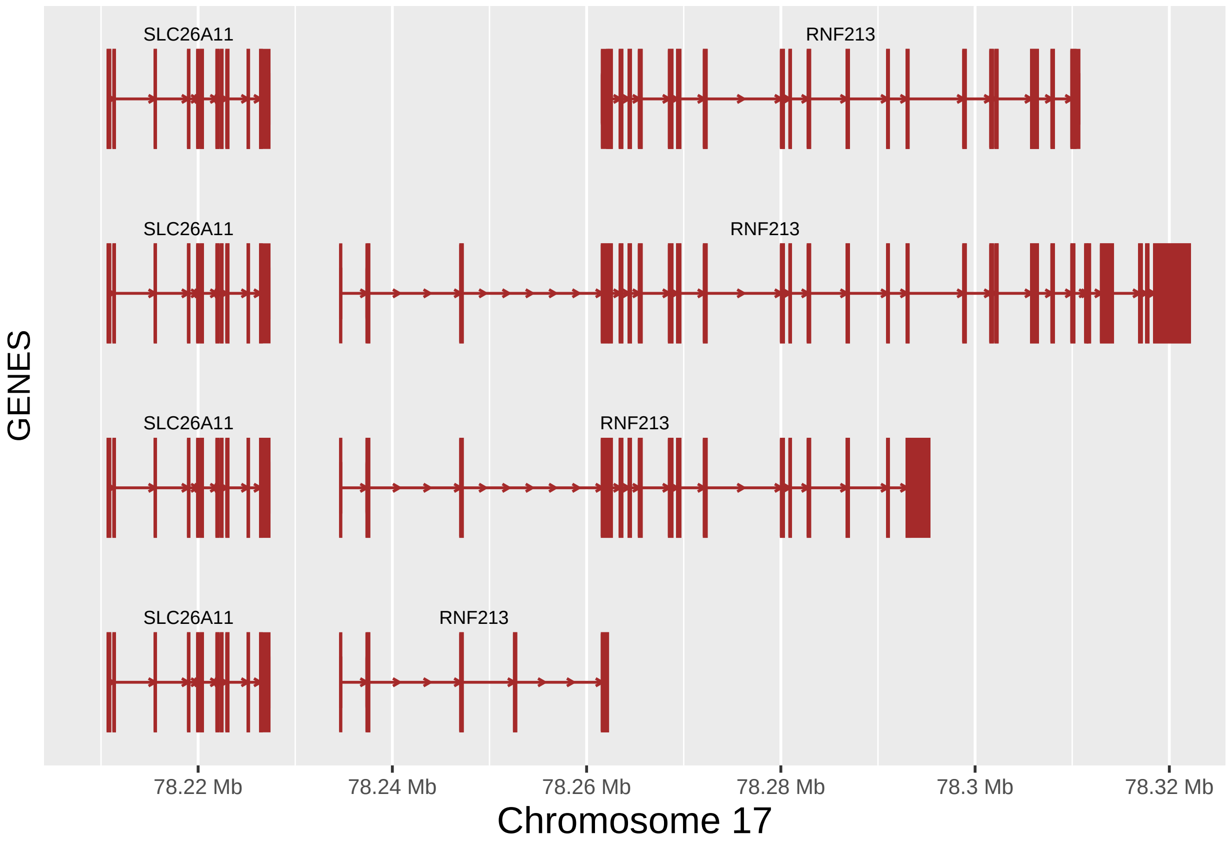


```r
# AORTA
RNF213_aorta_df <- as.data.frame(read.table("input_files/GTEx_files/RNF213_aorta.txt",
                                             sep = "\t", header = TRUE,
                                             na.strings = c(".", "NA"), stringsAsFactors = FALSE))

RNF213_aorta <- ggplot(RNF213_aorta_df, aes(x = position, y = -log10(pval_nominal))) +
        xlim (RNF213_range) +
        scale_y_continuous(limits = c(4,24), breaks=seq(4,24,4), expand = c(0,0)) +
        geom_point(size = 2.25, data=subset(RNF213_aorta_df,
                                            -log10(RNF213_aorta_df$pval_nominal)>4),
                   aes(position, -log10(pval_nominal)), color = "black") +
        geom_point(size = 1.25, data=subset(RNF213_aorta_df,
                                            -log10(RNF213_aorta_df$pval_nominal)>4),
                   aes(position, -log10(pval_nominal)), color = "red") +
        ##geom_text(data=subset(RNF213_aorta_df, Include == "Y"),
        ##           aes(position, -log10(pval_nominal),label=SNP_ID, hjust = 0, angle = 10)) +
        scale_x_sequnit("Mb") +
        labs(x = x_lab_aorta, y = y_lab) +
        theme(axis.text.x = element_text(size=8), axis.text.y = element_text(size=10),
              axis.title.x = element_text(size=14), axis.title.y = element_text(size=12))
fixed(RNF213_aorta) <- TRUE
RNF213_aorta
```

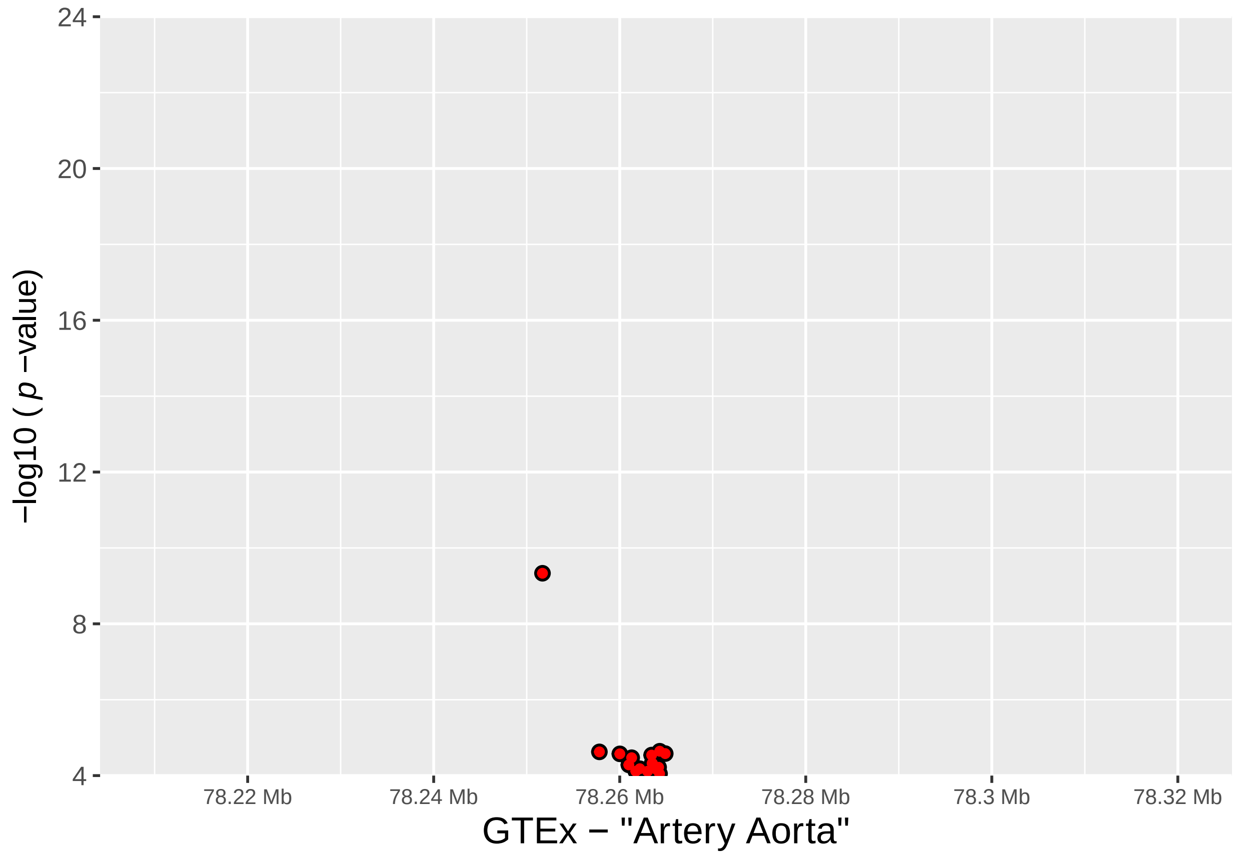


```
# CORONARY
RNF213_coronary_df <- as.data.frame(read.table("input_files/GTEx_files/RNF213_coronary.txt",
                                               sep = "\t", header = TRUE,
                                               na.strings = c(".", "NA"),
                                               stringsAsFactors = FALSE))

RNF213_coronary <- ggplot(RNF213_coronary_df, aes(x = position, y = -log10(pval_nominal))) +
        xlim (RNF213_range) +
        scale_y_continuous(limits = c(4,24), breaks=seq(4,24,4), expand = c(0,0)) +
        geom_point(size = 2.25, data=subset(RNF213_coronary_df,
                                            -log10(RNF213_coronary_df$pval_nominal)>4),
                   aes(position, -log10(pval_nominal)), color = "black") +
        geom_point(size = 1.25, data=subset(RNF213_coronary_df,
                                            -log10(RNF213_coronary_df$pval_nominal)>4),
                   aes(position, -log10(pval_nominal)), color = "red") +
        ##geom_text(data=subset(RNF213_coronary_df, Include == "Y"),
        ##          aes(position, -log10(pval_nominal),label=SNP_ID, hjust = 0, angle = 10)) +
        scale_x_sequnit("Mb") +
        labs(x = x_lab_coronary, y = y_lab) +
        theme(axis.text.x = element_text(size=8), axis.text.y = element_text(size=10),
              axis.title.x = element_text(size=14), axis.title.y = element_text(size=12))
fixed(RNF213_coronary) <- TRUE
RNF213_coronary
```

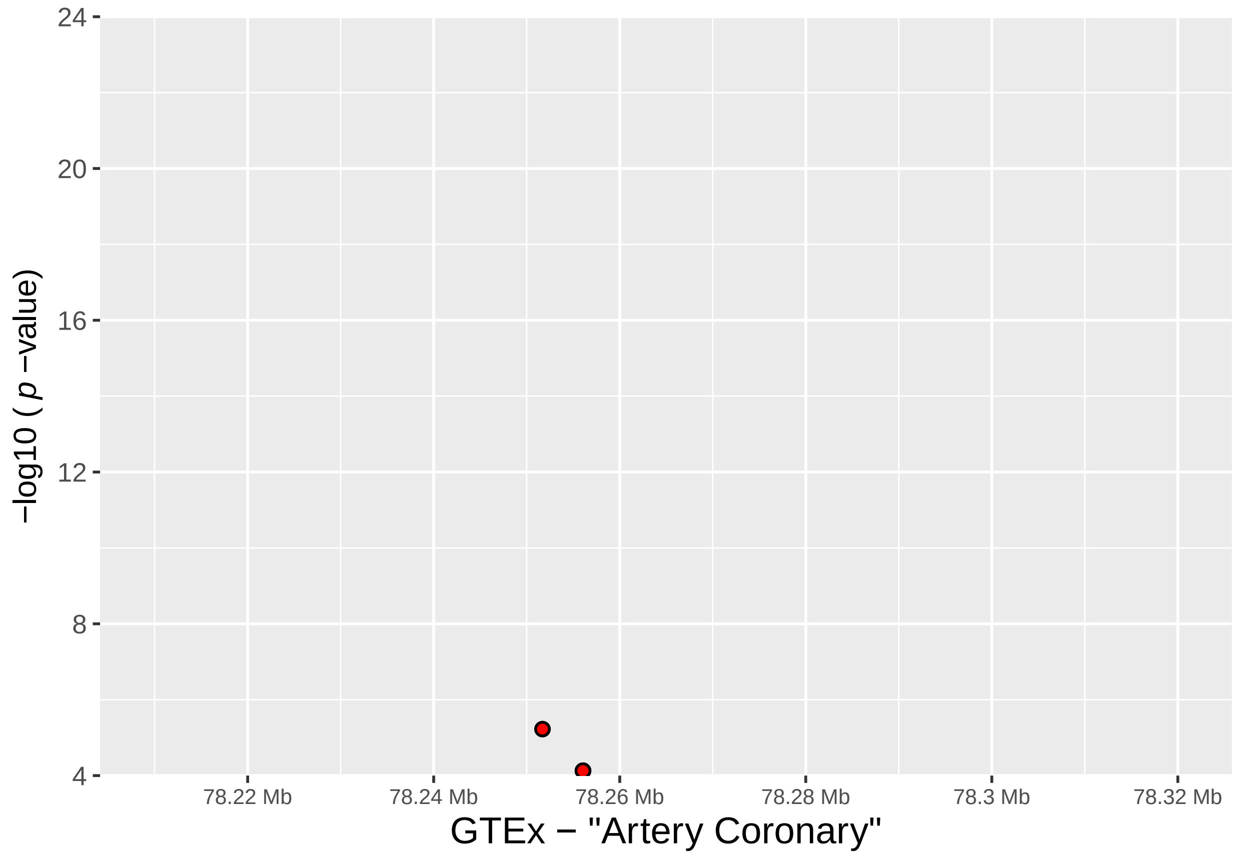


```
# TIBIAL
RNF213_tibial_df <- as.data.frame(read.table("input_files/GTEx_files/RNF213_tibial.txt",
                                             sep = "\t", header = TRUE,
                                             na.strings = c(".", "NA"),
                                             stringsAsFactors = FALSE))

RNF213_tibial <- ggplot(RNF213_tibial_df, aes(x = position, y = -log10(pval_nominal))) +
        xlim (RNF213_range) +
        scale_y_continuous(limits = c(4,24), breaks=seq(4,24,4), expand = c(0,0)) +
        geom_point(size = 2.25, data=subset(RNF213_tibial_df,
                                            -log10(RNF213_tibial_df$pval_nominal)>4),
                   aes(position, -log10(pval_nominal)), color = "black") +
        geom_point(size = 1.25, data=subset(RNF213_tibial_df,
                                            -log10(RNF213_tibial_df$pval_nominal)>4),
                   aes(position, -log10(pval_nominal)), color = "red") +
        ##geom_text(data=subset(RNF213_tibial_df, Include == "Y"),
        ##          aes(position, -log10(pval_nominal),label=SNP_ID, hjust = 0, angle = 10)) +
        scale_x_sequnit("Mb") +
        labs(x = x_lab_tibial, y = y_lab) +
        theme(axis.text.x = element_text(size=8), axis.text.y = element_text(size=10),
              axis.title.x = element_text(size=14), axis.title.y = element_text(size=12))
fixed(RNF213_tibial) <- TRUE
RNF213_tibial
```

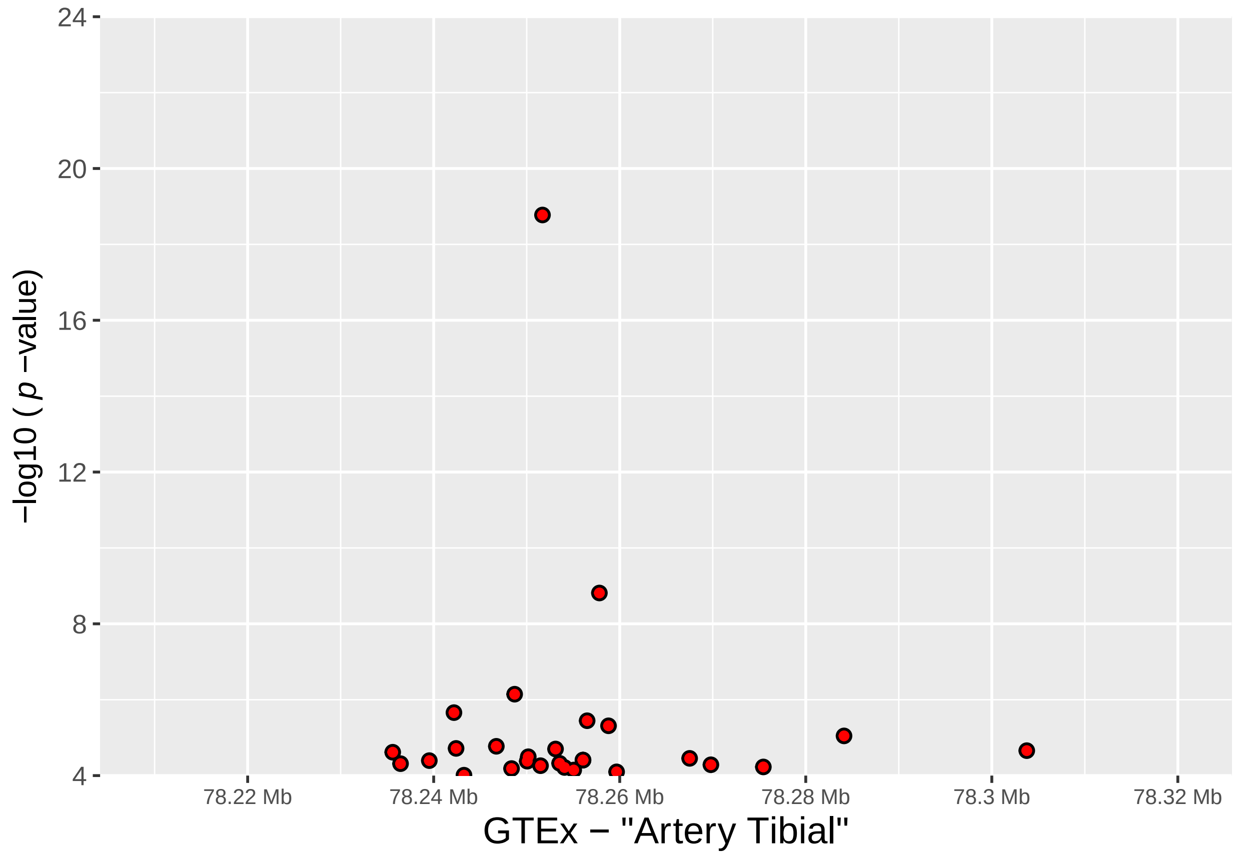


```
# Get the gtables
gA <- ggplotGrob (SLC26A11_aorta)
gB <- ggplotGrob (SLC26A11_coronary)
gC <- ggplotGrob (SLC26A11_tibial)
gD <- ggplotGrob (RNF213_aorta)
gE <- ggplotGrob (RNF213_coronary)
gF <- ggplotGrob (RNF213_tibial)
gG <- ggplotGrob (SLC26A11_GENES.gg)
gH <- ggplotGrob (RNF213_GENES.gg)

# Set the widths
gA$widths <- gC$widths
gB$widths <- gC$widths
gD$widths <- gC$widths
gE$widths <- gC$widths
gF$widths <- gC$widths
gG$widths <- gC$widths
gH$widths <- gC$widths

# Arrange the SLC26A11 charts
SLC26A11_title = textGrob("SLC26A11\n", gp=gpar(fontsize=12, font = 2))
grid.newpage()
SLC26A11_plot <- grid.arrange(gA, gB, gC, gG, heights = c(6,6,6,10), top =SLC26A11_title)
```

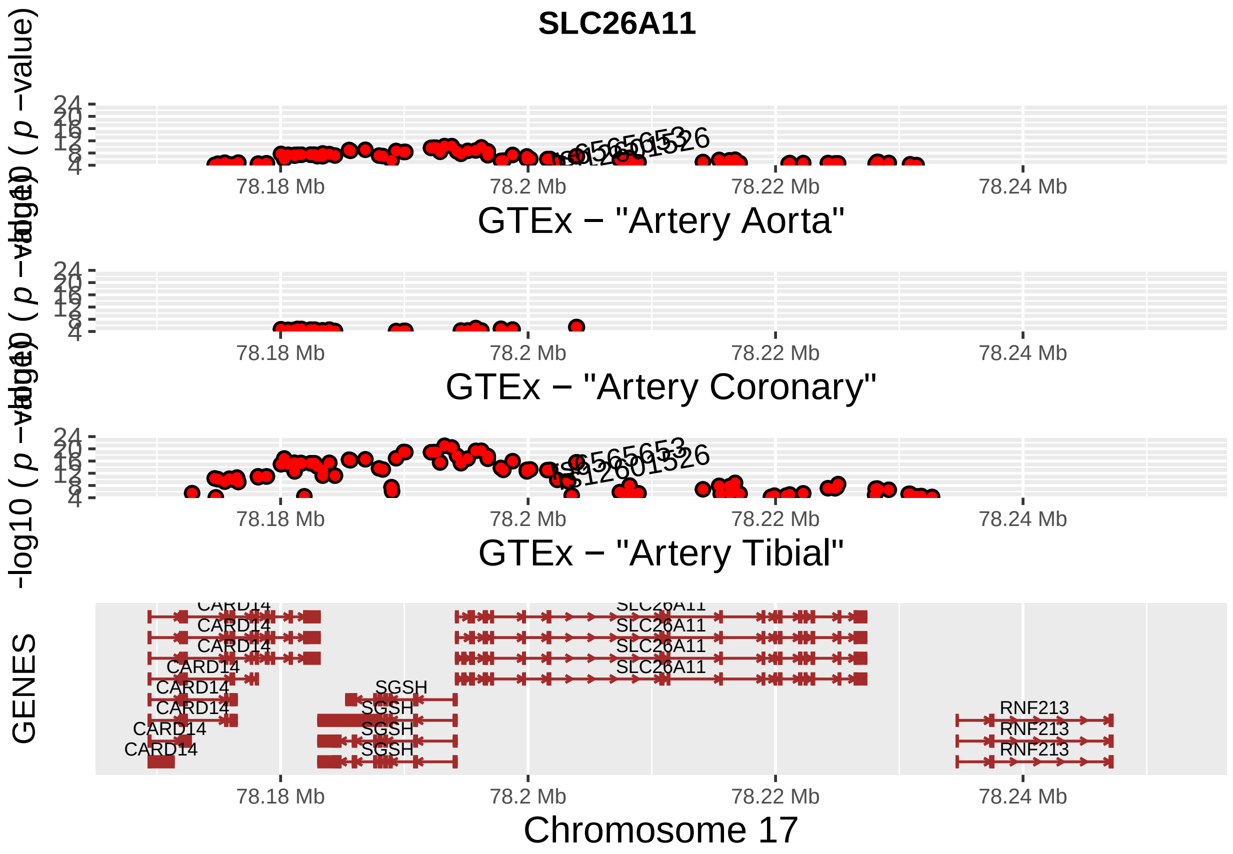


```
# Arrange the RNF213 charts
RNF213_title = textGrob("RNF213\n", gp=gpar(fontsize=12, font = 2))
grid.newpage()
RNF213_plot <- grid.arrange(gD, gE, gF, gH, heights = c(6,6,6,10), top =RNF213_title)
```

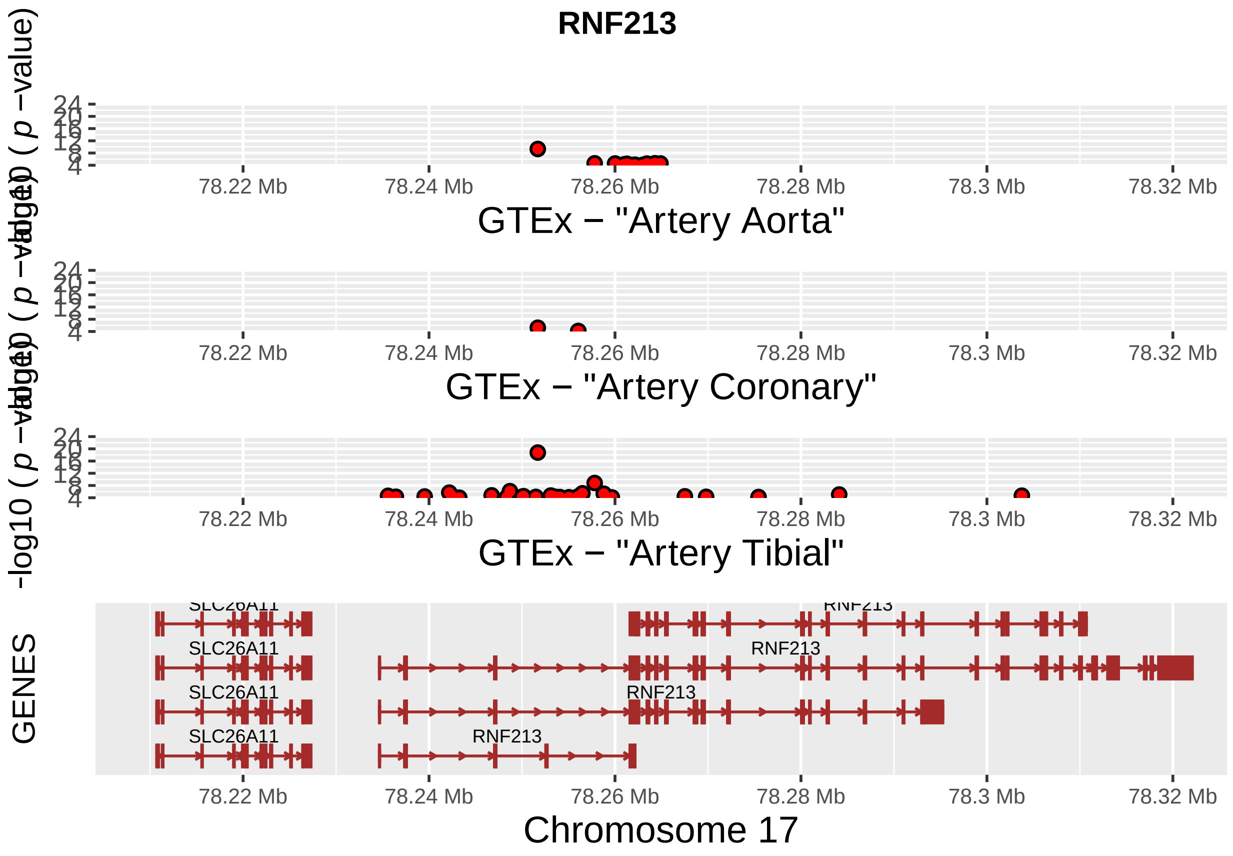


##### Supplementary Figure 4

```
# Arrange SLC26A11 - RNF213 plots together
main_title = textGrob("Arterial tissue cis-eQTL data for SLC26A11 and RNF213 SNPs and their expression\r
                      gp=gpar(fontsize=16, font = 2))
grid.newpage()
FINAL_PLOT <- grid.arrange(SLC26A11_plot, RNF213_plot, ncol = 2, top = main_title)
```

#### Arterial tissue cis−eQTL data for SLC26A11 and RNF213 SNPs and their expression

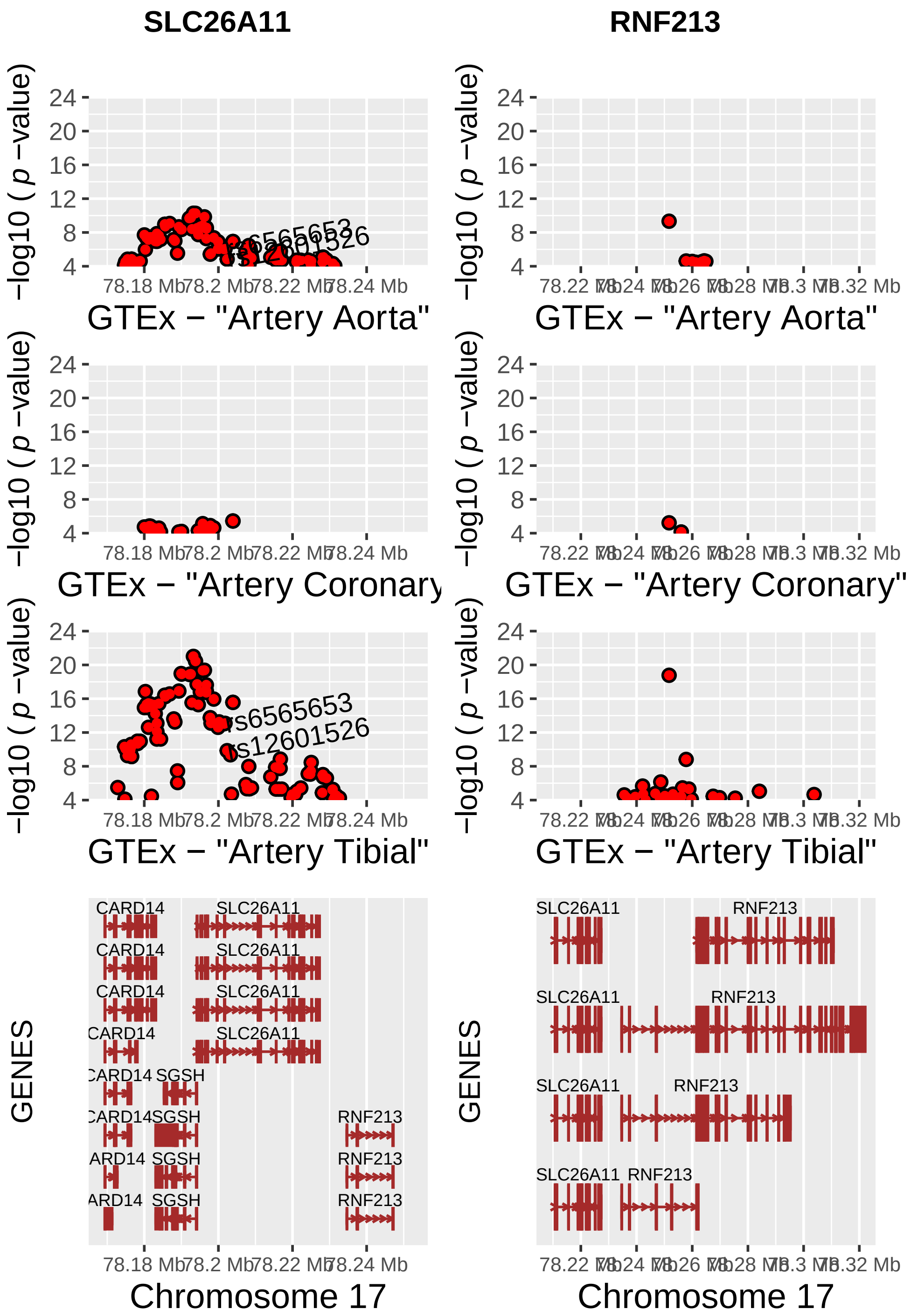


```r
dev.off()
```

```
## null device
##           1
```

#### Artery_expression_changes_RNF213_SLC26A11

Data extracted from the GTEx database

```
data(hg19IdeogramCyto, package = "biovizBase")
data(hg19Ideogram, package = "biovizBase")
data(genesymbol, package = "biovizBase")

hg19 <- keepSeqlevels(hg19IdeogramCyto, paste0("chr", c(1:22, "X", "Y")))

# RNF213_SLC26A11_eQTL_association_files

x_lab_aorta <- "GTEx - \"Artery Aorta\""
x_lab_coronary <- "GTEx - \"Artery Coronary\""
x_lab_tibial <- "GTEx - \"Artery Tibial\""
y_lab <- as.expression("Expression variation in %", fontsize=6, font = 2)
size_label = bquote("Magnitude of association\n-log10("*italic(p)*"-value)")

# Plots for SLC26A11 significant expression changes

# SLC26A11 Genes Plot
wh_SLC26A11 <- genesymbol[c("SLC26A11")]
SLC26A11_start <- as.vector(slot(wh_SLC26A11@ranges, "start") - 25000)
SLC26A11_end <- as.vector(slot(wh_SLC26A11@ranges, "start") +
                            slot (wh_SLC26A11@ranges, "width") - 1 + 25000)
SLC26A11_range <- GRanges('chr17', IRanges(start = SLC26A11_start, end = SLC26A11_end))
wh_SLC26A11 <- range(SLC26A11_range, ignore.strand = TRUE)


SLC26A11_GENES <- autoplot(Homo.sapiens, which = wh_SLC26A11, xlab = "Chromosome 17",
                          ylab = "GENES",
                          label.color = "black", color = "brown", fill = "brown",
                          columns = c("ALIAS", "GO"), scale = "Mb") +
        xlim (SLC26A11_range) +
        scale_x_sequnit("Mb") +
        theme(axis.text.x = element_text(size=8), axis.text.y = element_text(size=10),
              axis.title.x = element_text(size=14), axis.title.y = element_text(size=12))
SLC26A11_GENES
```

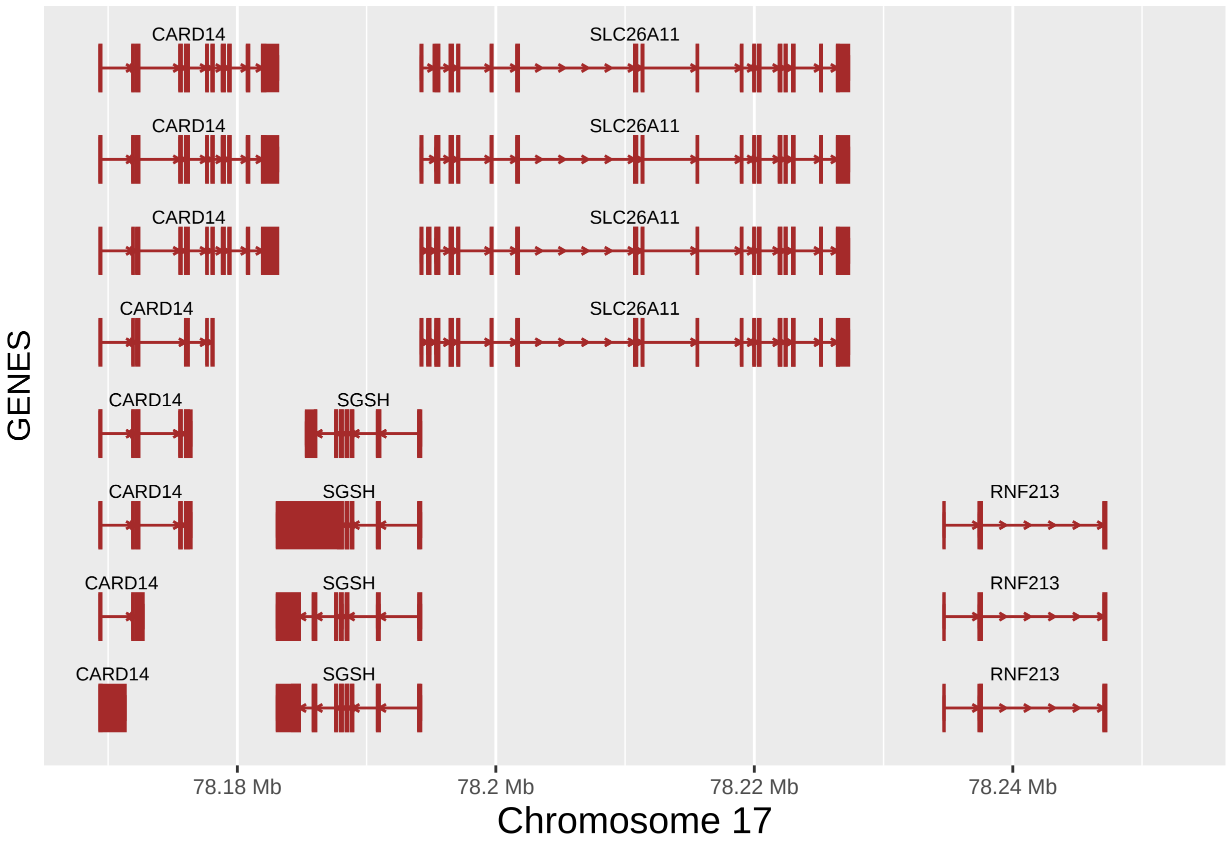


```
fixed (SLC26A11_GENES) <- TRUE

SLC26A11_GENES.gg <- SLC26A11_GENES@ggplot
SLC26A11_GENES.gg
```

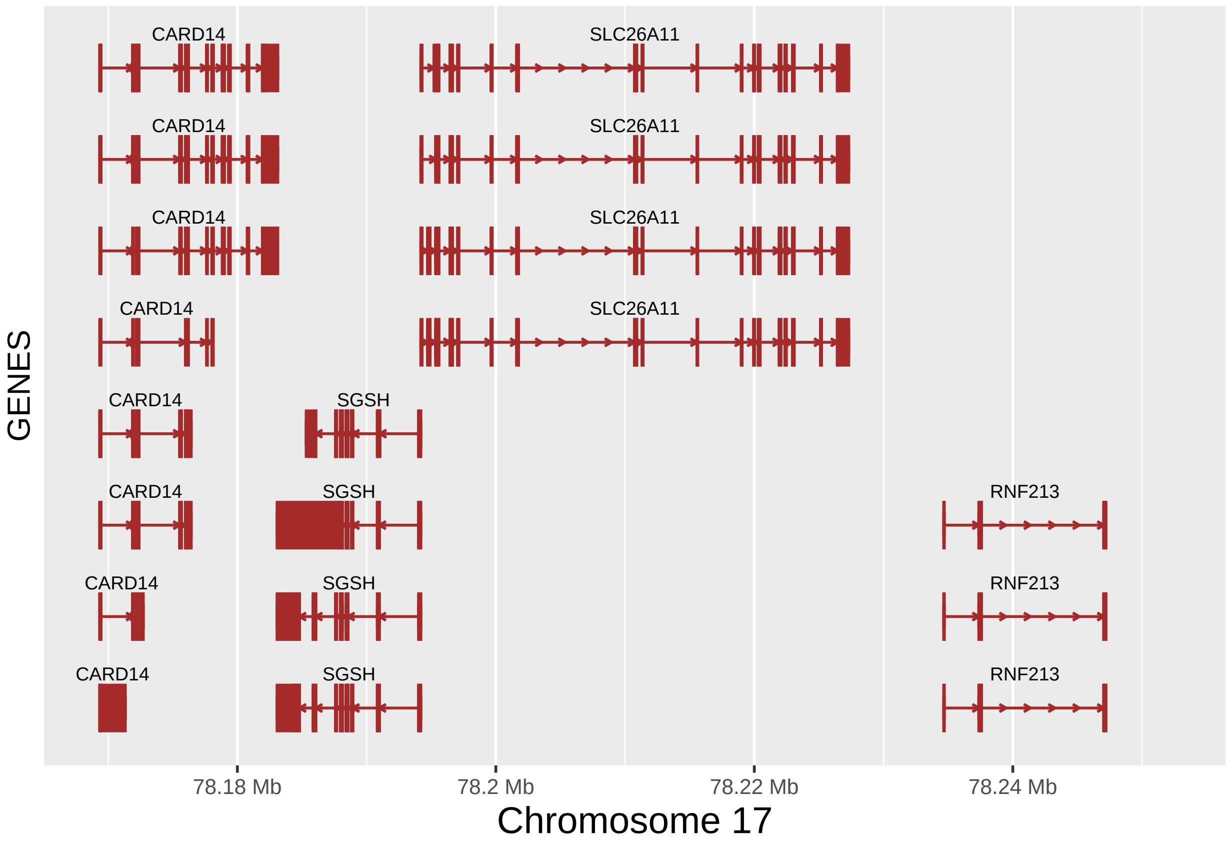


```
# SLC26A11 expression changes
# AORTA
SLC26A11_aorta_df <- as.data.frame(read.table("input_files/GTEx_files/SLC26A11_aorta.txt",
                                            sep = "\t", header = TRUE,
                                            na.strings = c(".", "NA"),
                                            stringsAsFactors = FALSE))

Sign_eQTLs_SAo <- subset(SLC26A11_aorta_df,-log10(SLC26A11_aorta_df$pval_nominal)>4)
Sign_eQTLs_SAo$log10 <- -log10(Sign_eQTLs_SAo$pval_nominal)
Sign_eQTLs_SAo$ranges <- cut(Sign_eQTLs_SAo$log10, seq(4,24,2),
                            labels = c("4 - 5.99","6 - 7.99","8 - 9.99","10 - 11.99",
                                       "12 - 13.99","14 - 15.99","16 - 17.99","18 - 19.99",
                                       "20 - 21.99","22 - 24"))
Sign_eQTLs_SAo$color <- ifelse(Sign_eQTLs_SAo$slope<0, "Underexpression","Overespression")


SLC26A11_aorta_ex <- ggplot(Sign_eQTLs_SAo) +
        xlim (SLC26A11_range) +
        scale_x_sequnit("Mb") +
        scale_y_continuous(limits = c(-0.8,0.8),
                          breaks=c(-0.8,-0.6,-0.4,-0.2,0,0.2,0.4,0.6,0.8), expand = c(0,0)) +
        geom_point(aes(position, slope, size = Sign_eQTLs_SAo$ranges,
                       color =Sign_eQTLs_SAo$color)) +
        geom_text(data=subset(SLC26A11_aorta_df, Include == "Y"),
                  aes( x= position, y = slope, label = SNP_ID), hjust = 0, angle = 10) +
```

```
        labs(x = x_lab_aorta, y = y_lab, size = size_label,
             color = "Expression change") +
        theme(axis.text.x = element_text(size=8), axis.text.y = element_text(size=10),
              axis.title.x = element_text(size=14), axis.title.y = element_text(size=12)) +
        theme(legend.position="none")
fixed(SLC26A11_aorta_ex) <- TRUE
SLC26A11_aorta_ex
```

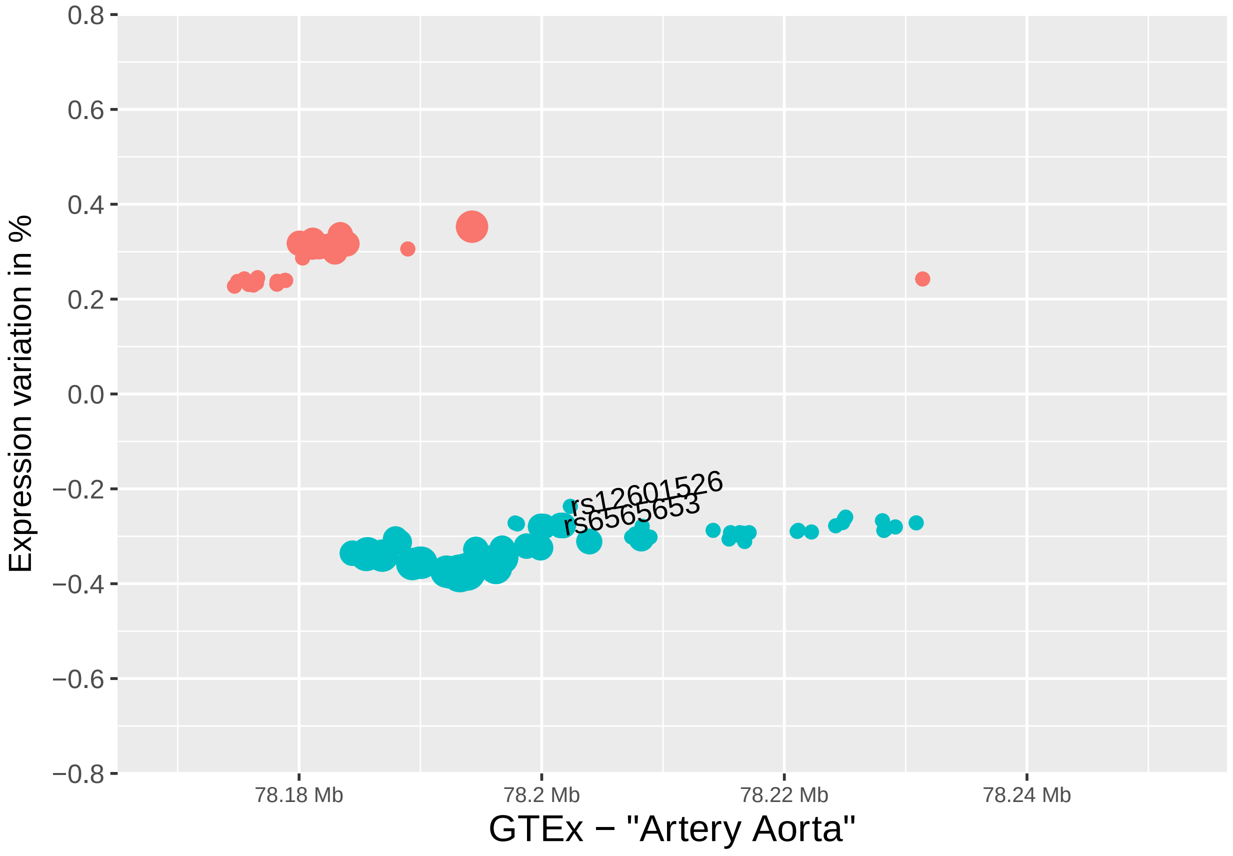


```
# CORONARY
SLC26A11_coronary_df <- as.data.frame(read.table("input_files/GTEx_files/SLC26A11_coronary.txt",
                                                 sep = "\t", header = TRUE,
                                                 na.strings = c(".", "NA"),
                                                 stringsAsFactors = FALSE))

Sign_eQTLs_SCo <- subset(SLC26A11_coronary_df,-log10(SLC26A11_coronary_df$pval_nominal)>4)
Sign_eQTLs_SCo$log10 <- -log10(Sign_eQTLs_SCo$pval_nominal)
Sign_eQTLs_SCo$ranges <- cut(Sign_eQTLs_SCo$log10, seq(4,24,2),
                             labels = c("4 - 5.99","6 - 7.99","8 - 9.99","10 - 11.99",
                                        "12 - 13.99","14 - 15.99","16 - 17.99","18 - 19.99",
                                        "20 - 21.99","22 - 24"))
Sign_eQTLs_SCo$color <- ifelse(Sign_eQTLs_SCo$slope<0, "Underexpression","Overespression")


SLC26A11_coronary_ex <- ggplot(Sign_eQTLs_SCo) +
```

```
        xlim (SLC26A11_range) +
        scale_x_sequnit("Mb") +
        scale_y_continuous(limits = c(-1,1),
                           breaks=c(-1.0,-0.8,-0.6,-0.4,-0.2,0,0.2,0.4,0.6,0.8,1.0),
                           expand = c(0,0)) +
        geom_point(aes(position, slope, size = Sign_eQTLs_SCo$ranges,
                       color =Sign_eQTLs_SCo$color)) +
        geom_text(data=subset(SLC26A11_coronary_df, Include == "Y"),
                  aes( x= position, y = slope, label = SNP_ID), hjust = 0, angle = 10) +
        labs(x = x_lab_coronary, y = y_lab, size = size_label,
             color = "Expression change") +
        theme(axis.text.x = element_text(size=8), axis.text.y = element_text(size=10),
              axis.title.x = element_text(size=14), axis.title.y = element_text(size=12)) +
        theme(legend.position="none")
fixed(SLC26A11_coronary_ex) <- TRUE
SLC26A11_coronary_ex
```

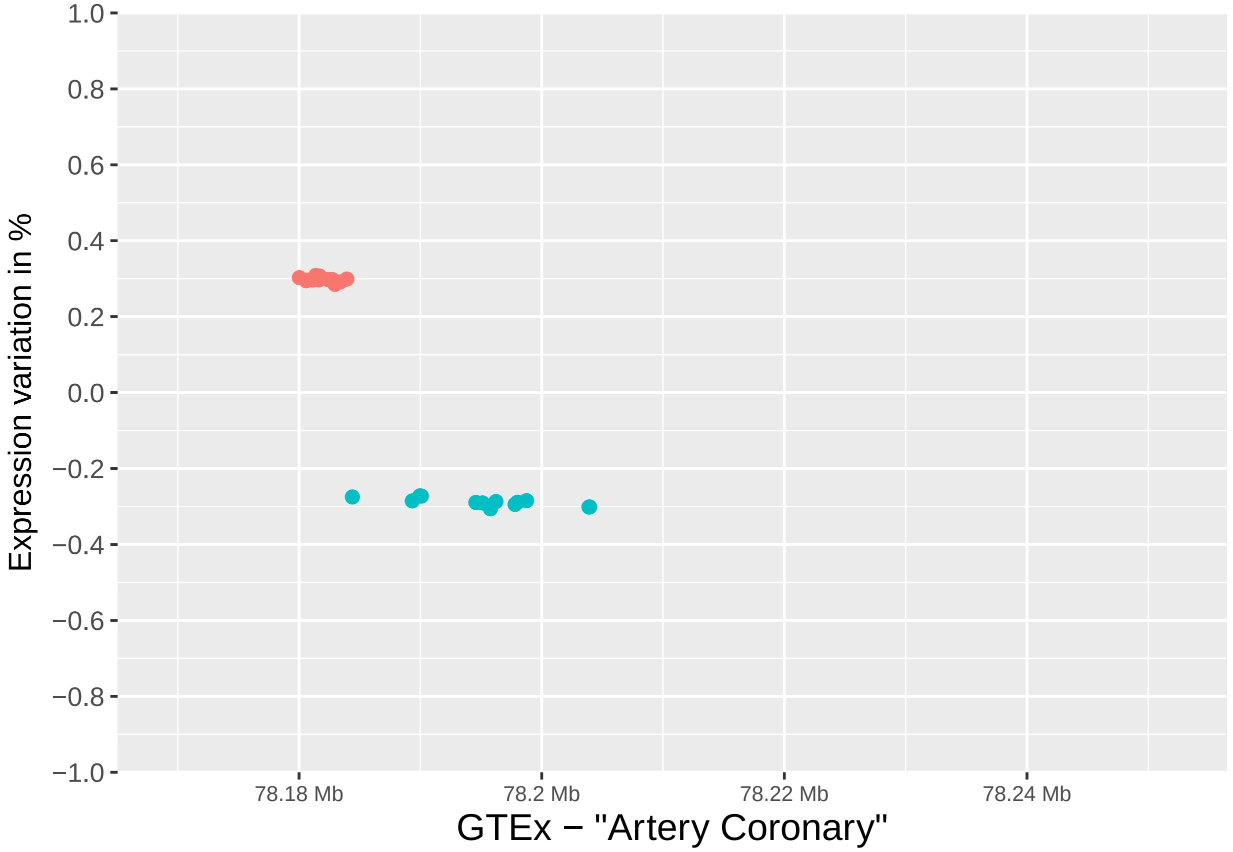


```
# TIBIAL
SLC26A11_tibial_df <- as.data.frame(read.table("input_files/GTEx_files/SLC26A11_tibial.txt",
                                               sep = "\t", header = TRUE,
                                               na.strings = c(".", "NA"),
                                               stringsAsFactors = FALSE))

Sign_eQTLs_STi <- subset(SLC26A11_tibial_df,-log10(SLC26A11_tibial_df$pval_nominal)>4)
```

```r
Sign_eQTLs_STi$log10 <- -log10(Sign_eQTLs_STi$pval_nominal)
Sign_eQTLs_STi$ranges <- cut(Sign_eQTLs_STi$log10, seq(4,24,2),
                             labels = c("4 - 5.99","6 - 7.99","8 - 9.99","10 - 11.99",
                                        "12 - 13.99","14 - 15.99","16 - 17.99","18 - 19.99",
                                        "20 - 21.99","22 - 24"))
Sign_eQTLs_STi$color <- ifelse(Sign_eQTLs_STi$slope<0, "Underexpression","Overespression")


SLC26A11_tibial_ex <- ggplot(Sign_eQTLs_STi) +
        xlim (SLC26A11_range) +
        scale_x_sequnit("Mb") +
        scale_y_continuous(limits = c(-1,1),
                           breaks=c(-1.0,-0.8,-0.6,-0.4,-0.2,0,0.2,0.4,0.6,0.8,1.0),
                           expand = c(0,0)) +
        geom_point(aes(position, slope, size = Sign_eQTLs_STi$ranges,
                       color =Sign_eQTLs_STi$color)) +
        geom_text(data=subset(SLC26A11_tibial_df, Include == "Y"),
                  aes( x= position, y = slope, label = SNP_ID), hjust = 0, angle = 10) +
        labs(x = x_lab_tibial, y = y_lab, size = size_label,
             color = "Expression change") +
        theme(axis.text.x = element_text(size=8), axis.text.y = element_text(size=10),
              axis.title.x = element_text(size=14), axis.title.y = element_text(size=12)) +
        theme(legend.position="none")
fixed(SLC26A11_tibial_ex) <- TRUE
SLC26A11_tibial_ex
```

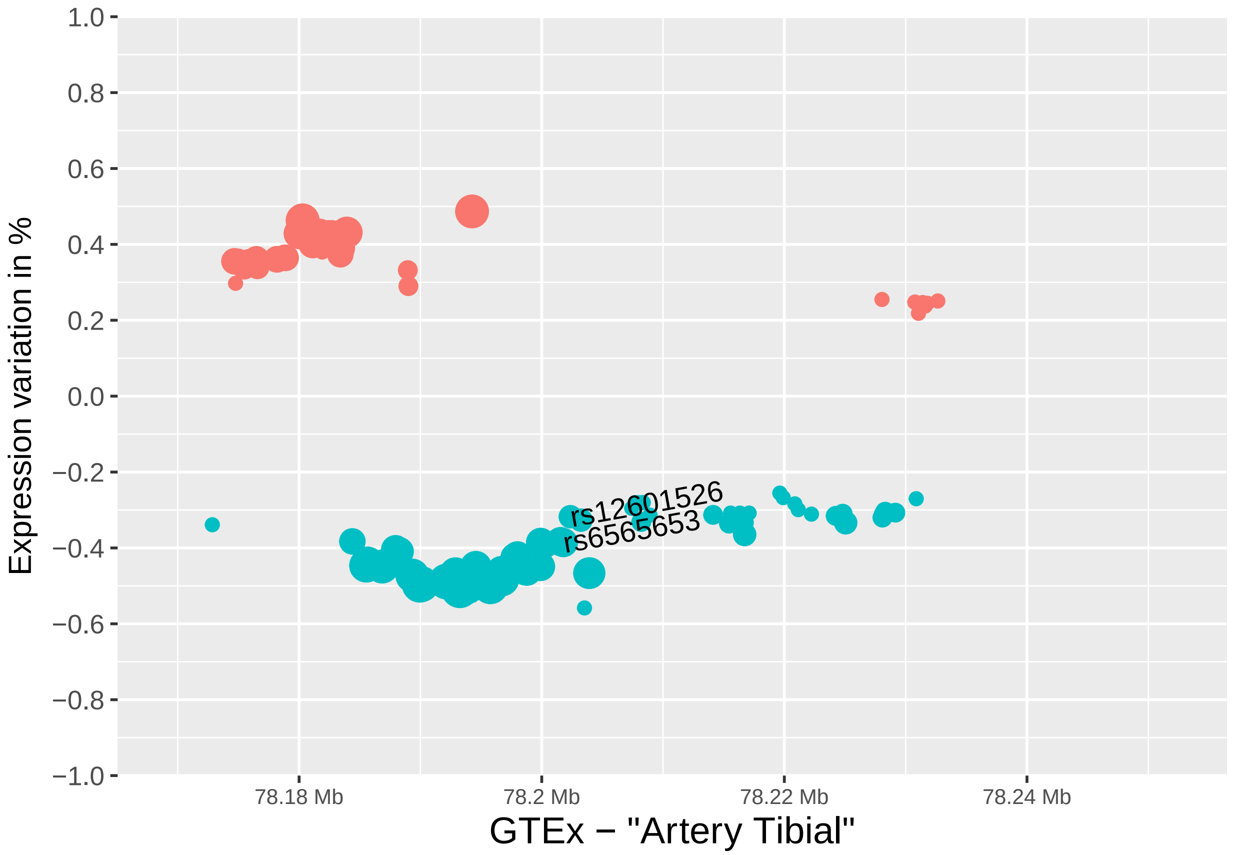


```
# Plots for RNF213 significant expression changes
# RNF213 Genes Plot
wh_RNF213 <- genesymbol[c("RNF213")]
RNF213_start <- as.vector(slot(wh_RNF213@ranges, "start") - 25000)
RNF213_end <- as.vector(slot(wh_RNF213@ranges, "start") +
                          slot (wh_RNF213@ranges, "width") - 1 + 25000)
RNF213_range <- GRanges('chr17', IRanges(start = RNF213_start, end = RNF213_end))
wh_RNF213 <- range(RNF213_range, ignore.strand = TRUE)


RNF213_GENES <- autoplot(Homo.sapiens, which = wh_RNF213,
                         xlab = "Chromosome 17", ylab = "GENES",
                         label.color = "black", color = "brown", fill = "brown",
                         columns = c("ALIAS", "GO"), scale = "Mb")+
        xlim (RNF213_range) +
        scale_x_sequnit("Mb") +
        theme(axis.text.x = element_text(size=8), axis.text.y = element_text(size=10),
              axis.title.x = element_text(size=14), axis.title.y = element_text(size=12))
RNF213_GENES
```

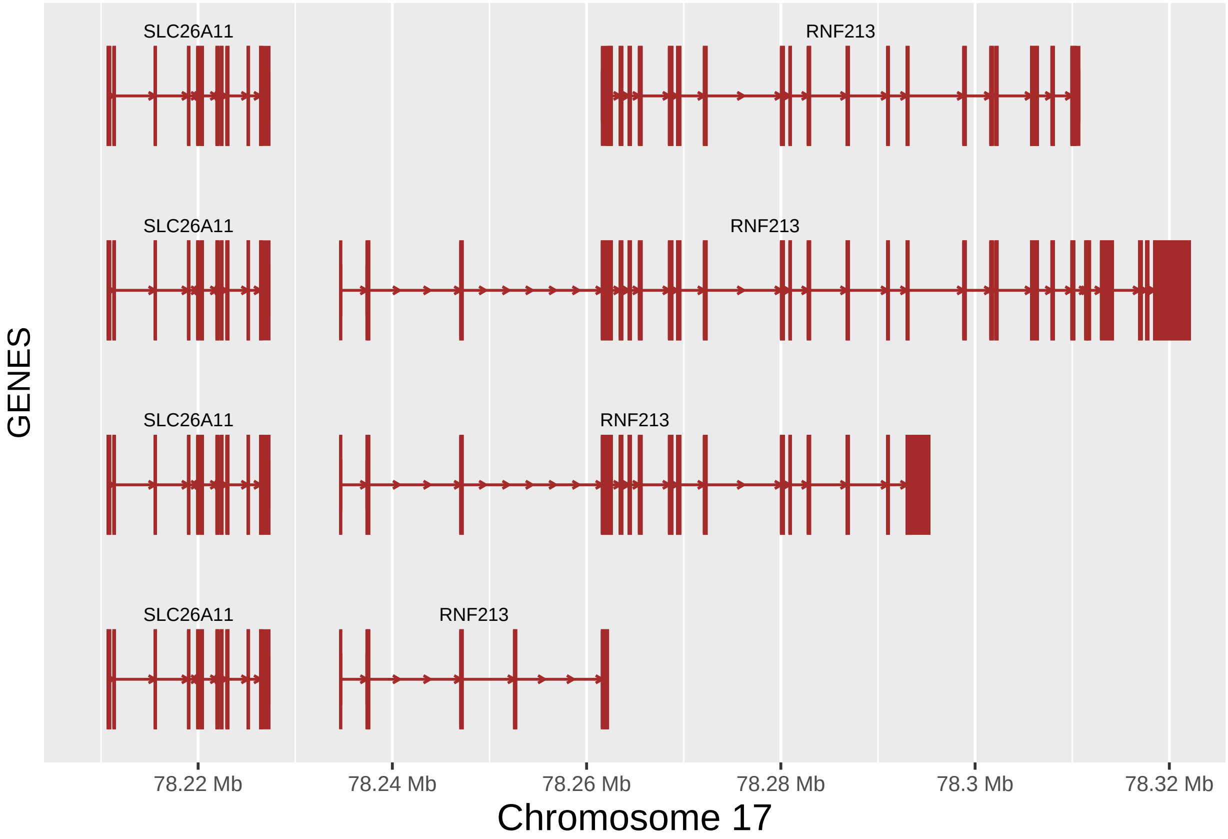


```
fixed (RNF213_GENES) <- TRUE

RNF213_GENES.gg <- RNF213_GENES@ggplot
RNF213_GENES.gg
```

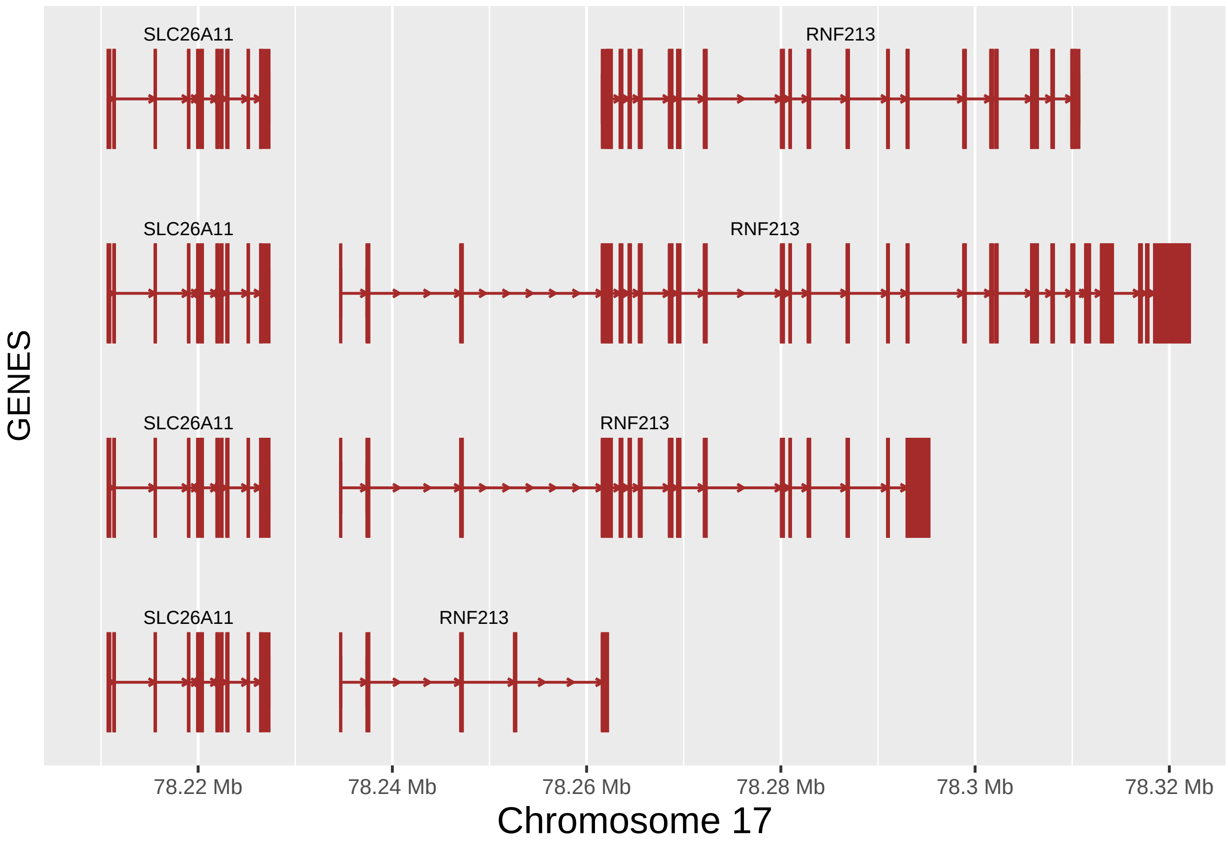


```
# RNF213 expression changes
# AORTA
RNF213_aorta_df <- as.data.frame(read.table("input_files/GTEx_files/RNF213_aorta.txt",
                                            sep = "\t", header = TRUE,
                                            na.strings = c(".", "NA"),
                                            stringsAsFactors = FALSE))

Sign_eQTLs_RAo <- subset(RNF213_aorta_df,-log10(RNF213_aorta_df$pval_nominal)>4)
Sign_eQTLs_RAo$log10 <- -log10(Sign_eQTLs_RAo$pval_nominal)
Sign_eQTLs_RAo$ranges <- cut(Sign_eQTLs_RAo$log10, seq(4,24,2),
                             labels = c("4 - 5.99","6 - 7.99","8 - 9.99","10 - 11.99",
                                        "12 - 13.99","14 - 15.99","16 - 17.99","18 - 19.99",
                                        "20 - 21.99","22 - 24"))
Sign_eQTLs_RAo$color <- ifelse(Sign_eQTLs_RAo$slope<0, "Underexpression","Overespression")


RNF213_aorta_ex <- ggplot(Sign_eQTLs_RAo) +
        xlim (RNF213_range) +
        scale_x_sequnit("Mb") +
        scale_y_continuous(limits = c(-1,1),
                           breaks=c(-1.0,-0.8,-0.6,-0.4,-0.2,0,0.2,0.4,0.6,0.8,1.0),
                           expand = c(0,0)) +
        geom_point(aes(position, slope, size = Sign_eQTLs_RAo$ranges,
                       color =Sign_eQTLs_RAo$color)) +
        #geom_text(data=subset(RNF213_aorta_df, Include == "Y"),
```

```r
        #           aes( x= position, y = slope, label = SNP_ID), hjust = 0, angle = 10) +
        labs(x = x_lab_aorta, y = y_lab, size = size_label,
             color = "Expression change") +
        theme(axis.text.x = element_text(size=8), axis.text.y = element_text(size=10),
              axis.title.x = element_text(size=14), axis.title.y = element_text(size=12)) +
        theme(legend.position="none")
fixed(RNF213_aorta_ex) <- TRUE
RNF213_aorta_ex
```

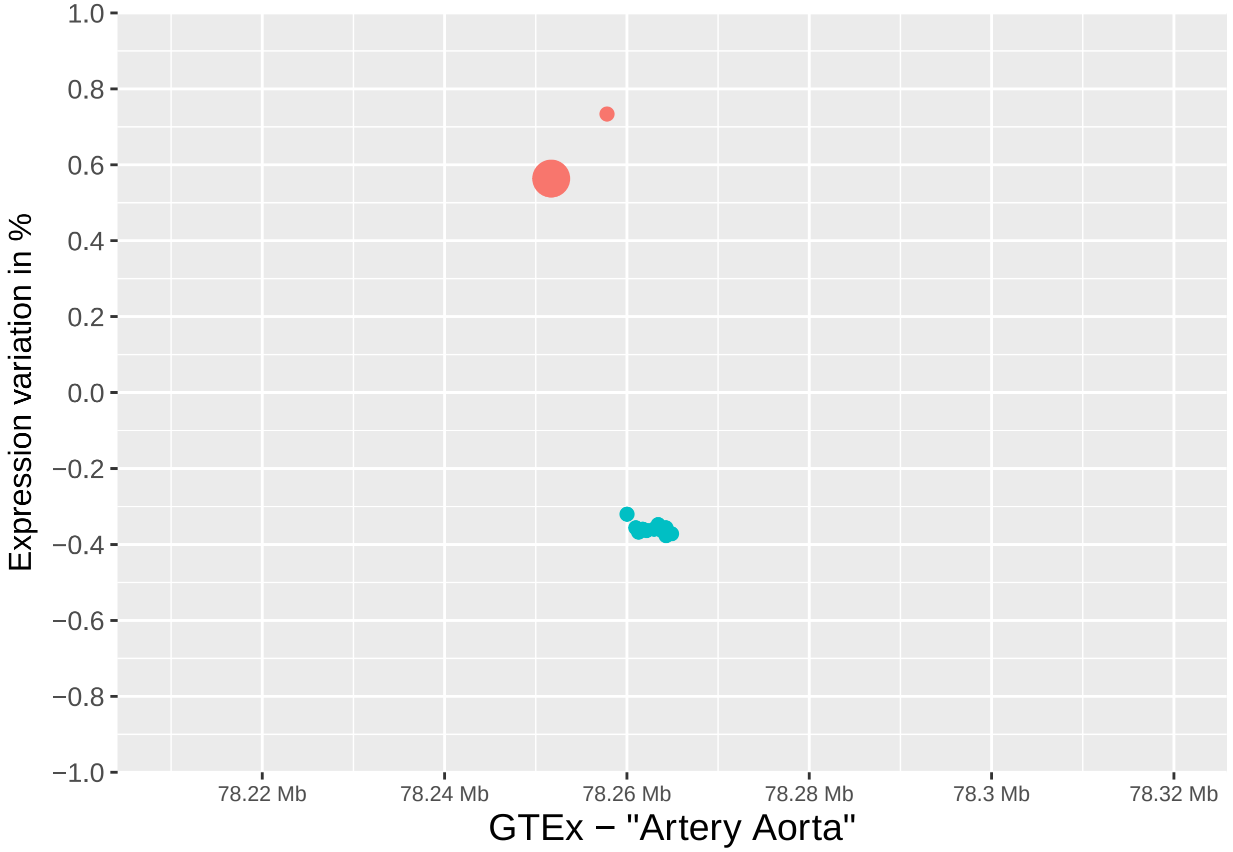


```r
# CORONARY
RNF213_coronary_df <- as.data.frame(read.table("input_files/GTEx_files/RNF213_coronary.txt",
                                               sep = "\t", header = TRUE,
                                               na.strings = c(".", "NA"),
                                               stringsAsFactors = FALSE))

Sign_eQTLs_RCo <- subset(RNF213_coronary_df,-log10(RNF213_coronary_df$pval_nominal)>4)
Sign_eQTLs_RCo$log10 <- -log10(Sign_eQTLs_RCo$pval_nominal)
Sign_eQTLs_RCo$ranges <- cut(Sign_eQTLs_RCo$log10, seq(4,24,2),
                             labels = c("4 - 5.99","6 - 7.99","8 - 9.99","10 - 11.99",
                                        "12 - 13.99","14 - 15.99","16 - 17.99","18 - 19.99",
                                        "20 - 21.99","22 - 24"))
Sign_eQTLs_RCo$color <- ifelse(Sign_eQTLs_RCo$slope<0, "Underexpression","Overespression")
```

```r
RNF213_coronary_ex <- ggplot(Sign_eQTLs_RCo) +
        xlim (RNF213_range) +
        scale_x_sequnit("Mb") +
        scale_y_continuous(limits = c(-1,1),
                           breaks=c(-1.0,-0.8,-0.6,-0.4,-0.2,0,0.2,0.4,0.6,0.8,1.0),
                           expand = c(0,0)) +
        geom_point(aes(position, slope, size = Sign_eQTLs_RCo$ranges,
                       color =Sign_eQTLs_RCo$color)) +
        #geom_text(data=subset(RNF213_coronary_df, Include == "Y"),
        #          aes( x= position, y = slope, label = SNP_ID), hjust = 0, angle = 10) +
        labs(x = x_lab_coronary, y = y_lab, size = size_label,
             color = "Expression change") +
        theme(axis.text.x = element_text(size=8), axis.text.y = element_text(size=10),
              axis.title.x = element_text(size=14), axis.title.y = element_text(size=12)) +
        theme(legend.position="none")
fixed(RNF213_coronary_ex) <- TRUE
RNF213_coronary_ex
```

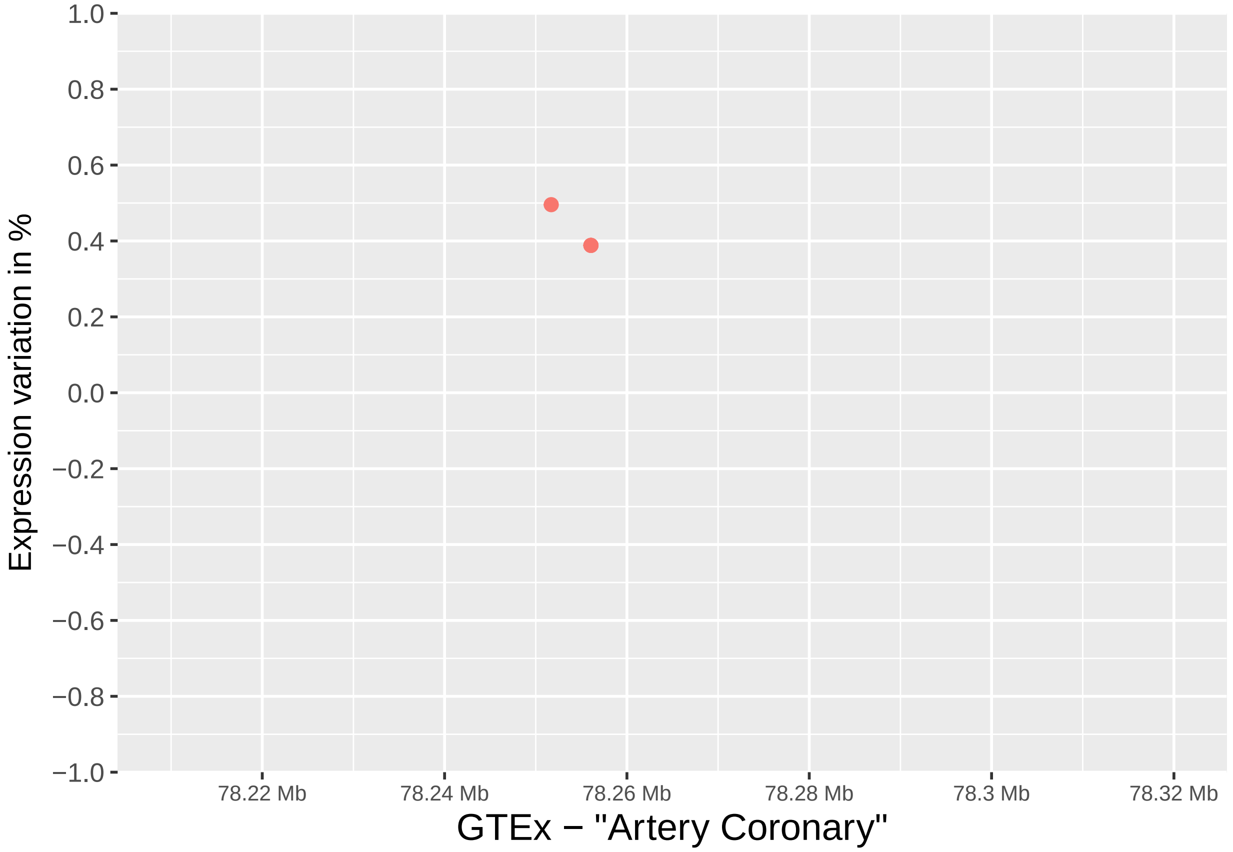


```r
# TIBIAL
RNF213_tibial_df <- as.data.frame(read.table("input_files/GTEx_files/RNF213_tibial.txt",
                                             sep = "\t", header = TRUE,
                                             na.strings = c(".", "NA"),
                                             stringsAsFactors = FALSE))
```

```
Sign_eQTLs_RTi <- subset(RNF213_tibial_df,-log10(RNF213_tibial_df$pval_nominal)>4)
Sign_eQTLs_RTi$log10 <- -log10(Sign_eQTLs_RTi$pval_nominal)
Sign_eQTLs_RTi$ranges <- cut(Sign_eQTLs_RTi$log10, seq(4,24,2),
                             labels = c("4 - 5.99","6 - 7.99","8 - 9.99","10 - 11.99",
                                        "12 - 13.99","14 - 15.99","16 - 17.99","18 - 19.99",
                                        "20 - 21.99","22 - 24"))
Sign_eQTLs_RTi$color <- ifelse(Sign_eQTLs_RTi$slope<0, "Underexpression","Overespression")


RNF213_tibial_ex <- ggplot(Sign_eQTLs_RTi) +
        xlim (RNF213_range) +
        scale_x_sequnit("Mb") +
        scale_y_continuous(limits = c(-1,1),
                           breaks=c(-1.0,-0.8,-0.6,-0.4,-0.2,0,0.2,0.4,0.6,0.8,1.0),
                           expand = c(0,0)) +
        geom_point(aes(position, slope, size = Sign_eQTLs_RTi$ranges,
                       color =Sign_eQTLs_RTi$color)) +
        geom_text(data=subset(RNF213_tibial_df, Include == "Y"),
                  aes( x= position, y = slope, label = SNP_ID), hjust = 0, angle = 10) +
        labs(x = x_lab_tibial, y = y_lab, size = size_label,
             color = "Expression change") +
        theme(axis.text.x = element_text(size=8), axis.text.y = element_text(size=10),
              axis.title.x = element_text(size=14), axis.title.y = element_text(size=12)) +
        theme(legend.position="none")
fixed(RNF213_tibial_ex) <- TRUE
RNF213_tibial_ex
```

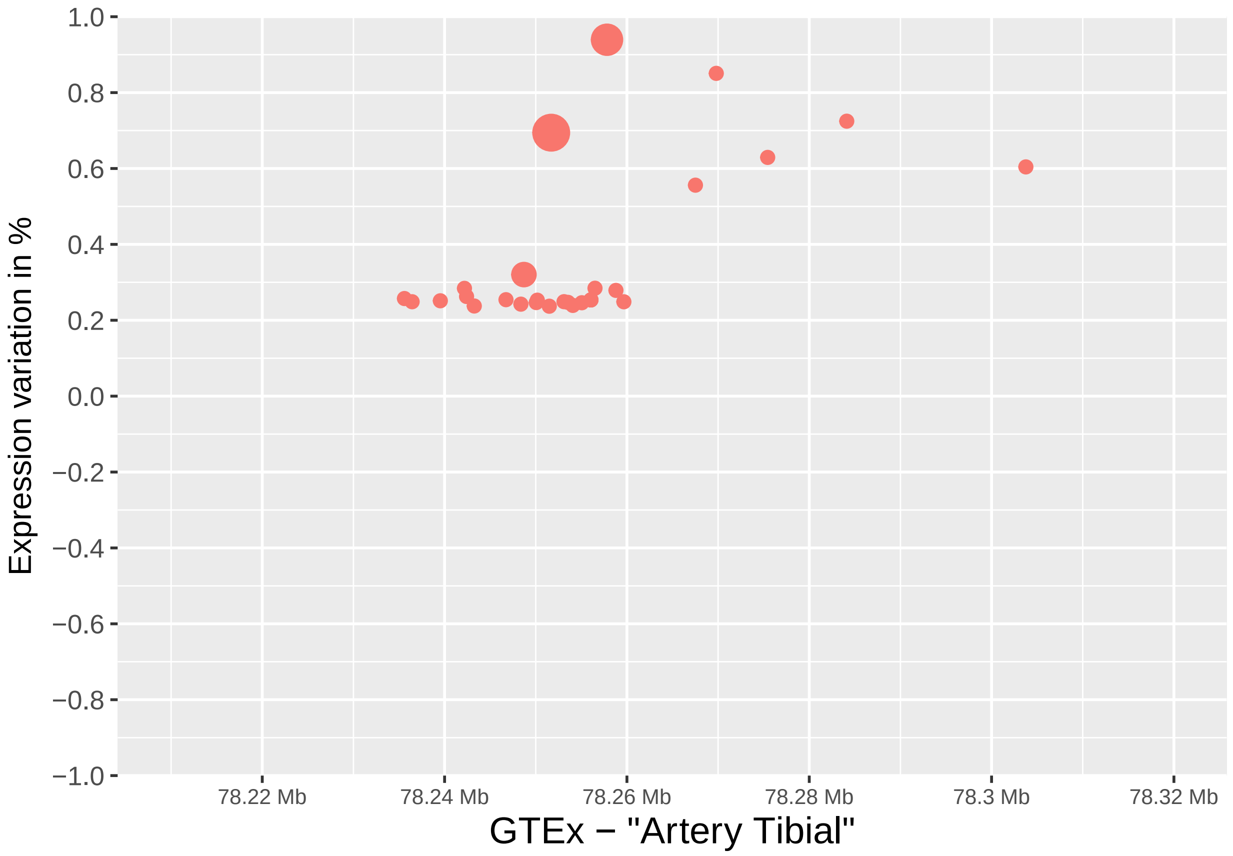


```
# Get the gtables
gA <- ggplotGrob (SLC26A11_aorta_ex)
gB <- ggplotGrob (SLC26A11_coronary_ex)
gC <- ggplotGrob (SLC26A11_tibial_ex)
gD <- ggplotGrob (RNF213_aorta_ex)
gE <- ggplotGrob (RNF213_coronary_ex)
gF <- ggplotGrob (RNF213_tibial_ex)
gG <- ggplotGrob (SLC26A11_GENES.gg)
gH <- ggplotGrob (RNF213_GENES.gg)

# Set the widths
gA$widths <- gC$widths
gB$widths <- gC$widths
gD$widths <- gC$widths
gE$widths <- gC$widths
gF$widths <- gC$widths
gG$widths <- gC$widths
gH$widths <- gC$widths

# Arrange the SLC26A11 charts
SLC26A11_title = textGrob("SLC26A11 expression\n", gp=gpar(fontsize=12, font = 2))
grid.newpage()
SLC26A11_plot <- grid.arrange(gA, gB, gC, gG, heights = c(8,8,8,10), top =SLC26A11_title)
```

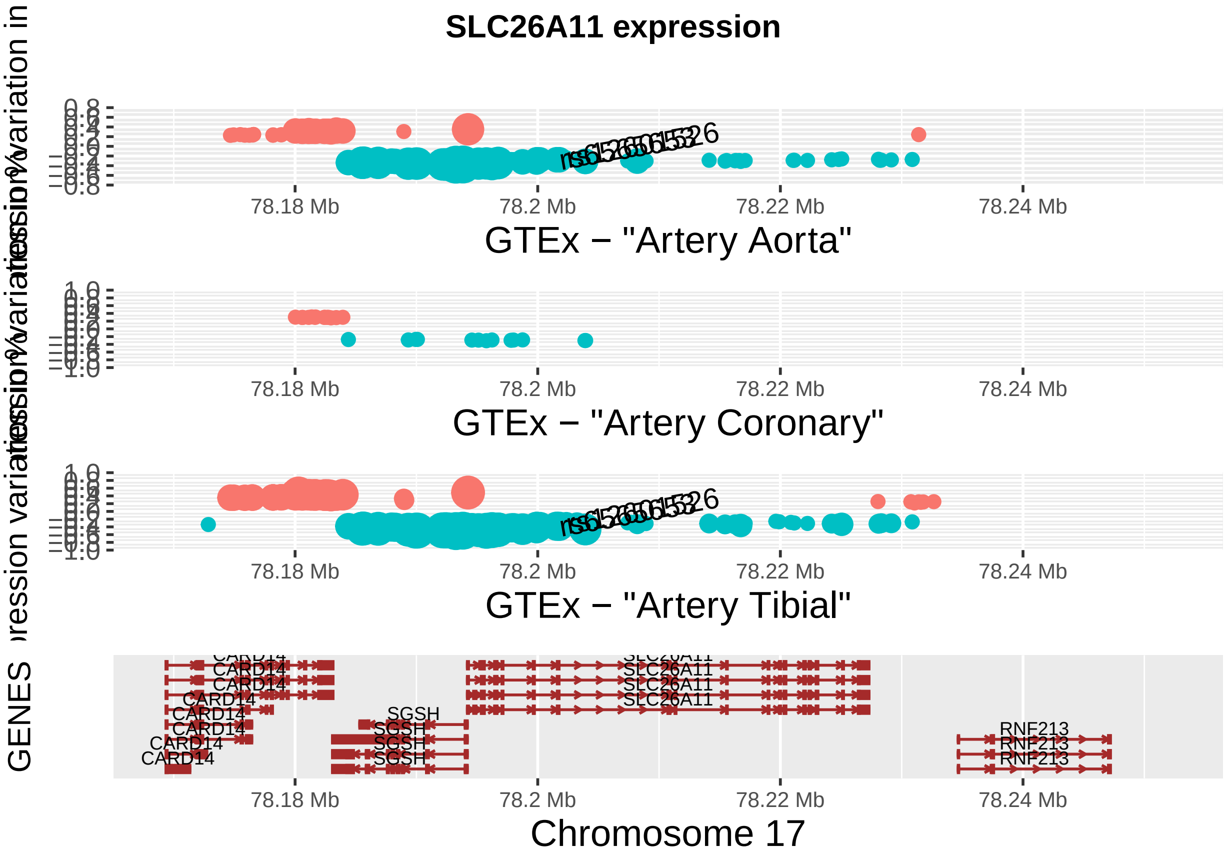


```
# Arrange the RNF213 charts
RNF213_title = textGrob("RNF213 expression\n", gp=gpar(fontsize=12, font = 2))
grid.newpage()
RNF213_plot <- grid.arrange(gD, gE, gF, gH, heights = c(8,8,8,10), top =RNF213_title)
```

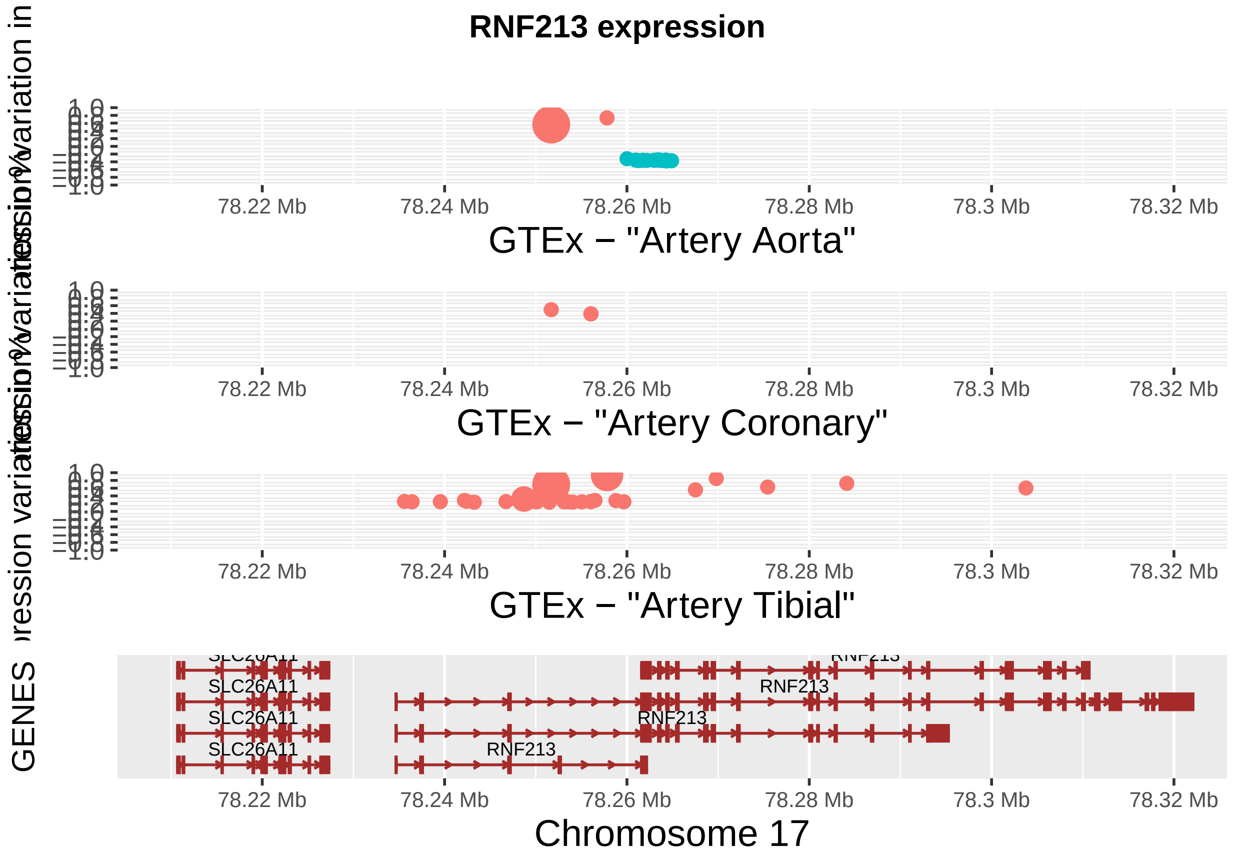


**Supplementary Figure 5**

```r
# Arrange SLC26A11 - RNF213 plots together
main_title = textGrob("Arterial tissue expression changes due to the SLC26A11 and RNF213 variation\n",
                      gp=gpar(fontsize=16, font = 2))
grid.newpage()

FINAL_PLOT <- grid.arrange(SLC26A11_plot, RNF213_plot, ncol = 2, top = main_title)
```

### Arterial tissue expression changes due to the SLC26A11 and RNF213 variation

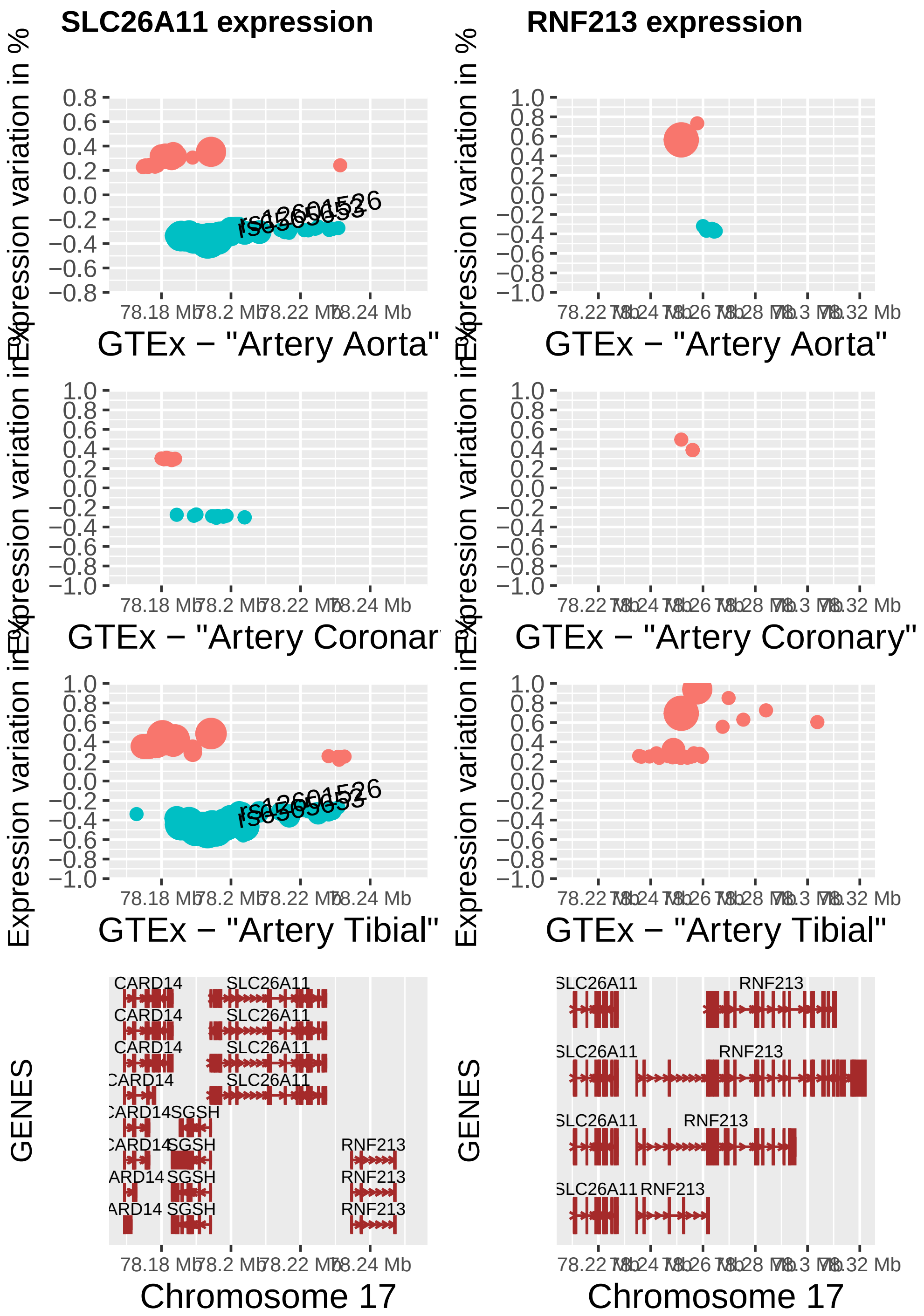


```
dev.off()
```

```
## null device
##           1
```

```r
sessionInfo()
```

```
## R version 3.5.1 (2018-07-02)
## Platform: x86_64-apple-darwin15.6.0 (64-bit)
## Running under: macOS High Sierra 10.13.6
##
## Matrix products: default
## BLAS: /Library/Frameworks/R.framework/Versions/3.5/Resources/lib/libRblas.0.dylib
## LAPACK: /Library/Frameworks/R.framework/Versions/3.5/Resources/lib/libRlapack.dylib
##
## locale:
## [1] en_US.UTF-8/en_US.UTF-8/en_US.UTF-8/C/en_US.UTF-8/en_US.UTF-8
##
## attached base packages:
##  [1] stats4    grid      parallel  stats     graphics  grDevices utils
##  [8] datasets  methods   base
##
## other attached packages:
##  [1] VariantAnnotation_1.28.13
##  [2] SummarizedExperiment_1.12.0
##  [3] DelayedArray_0.8.0
##  [4] BiocParallel_1.16.6
##  [5] matrixStats_0.55.0
##  [6] scales_1.1.0
##  [7] Rsamtools_1.34.1
##  [8] reshape_0.8.8
##  [9] raster_3.0-7
## [10] sp_1.3-2
## [11] plyr_1.8.5
## [12] LDheatmap_0.99-7
## [13] BSgenome.Hsapiens.UCSC.hg19_1.4.0
## [14] BSgenome_1.50.0
## [15] rtracklayer_1.42.2
## [16] Biostrings_2.50.2
## [17] XVector_0.22.0
## [18] Homo.sapiens_1.3.1
## [19] TxDb.Hsapiens.UCSC.hg19.knownGene_3.2.2
## [20] org.Hs.eg.db_3.7.0
## [21] GO.db_3.7.0
## [22] OrganismDbi_1.24.0
## [23] genetics_1.3.8.1.2
## [24] mvtnorm_1.0-11
## [25] MASS_7.3-51.5
## [26] gtools_3.8.1
## [27] gdata_2.18.0
## [28] combinat_0.0-8
## [29] ensembldb_2.6.8
## [30] AnnotationFilter_1.6.0
## [31] GenomicFeatures_1.34.8
## [32] AnnotationDbi_1.44.0
## [33] Biobase_2.42.0
## [34] GenomicRanges_1.34.0
```

```
## [35] GenomeInfoDb_1.18.2
## [36] IRanges_2.16.0
## [37] S4Vectors_0.20.1
## [38] cowplot_1.0.0
## [39] gtable_0.3.0
## [40] png_0.1-7
## [41] gridExtra_2.3
## [42] ggbio_1.30.0
## [43] BiocGenerics_0.28.0
## [44] extrafont_0.17
## [45] ggplot2_3.2.1
##
## loaded via a namespace (and not attached):
##  [1] colorspace_1.4-1        biovizBase_1.30.1        htmlTable_1.13.3
##  [4] base64enc_0.1-3         dichromat_2.0-0          rstudioapi_0.10
##  [7] farver_2.0.1            bit64_0.9-7              codetools_0.2-16
## [10] splines_3.5.1           knitr_1.26               zeallot_0.1.0
## [13] Formula_1.2-3           Rttf2pt1_1.3.7           cluster_2.1.0
## [16] graph_1.60.0            BiocManager_1.30.10      compiler_3.5.1
## [19] httr_1.4.1              backports_1.1.5          assertthat_0.2.1
## [22] Matrix_1.2-18           lazyeval_0.2.2           acepack_1.4.1
## [25] htmltools_0.4.0         prettyunits_1.0.2        tools_3.5.1
## [28] glue_1.3.1              GenomeInfoDbData_1.2.0   reshape2_1.4.3
## [31] dplyr_0.8.3             Rcpp_1.0.3               vctrs_0.2.1
## [34] extrafontdb_1.0         xfun_0.11                stringr_1.4.0
## [37] lifecycle_0.1.0         XML_3.98-1.20            zlibbioc_1.28.0
## [40] hms_0.5.2               ProtGenerics_1.14.0      RBGL_1.58.2
## [43] RColorBrewer_1.1-2      yaml_2.2.0               curl_4.3
## [46] memoise_1.1.0           biomaRt_2.38.0           rpart_4.1-15
## [49] latticeExtra_0.6-28     stringi_1.4.3            RSQLite_2.1.5
## [52] checkmate_1.9.4         rlang_0.4.2              pkgconfig_2.0.3
## [55] bitops_1.0-6            evaluate_0.14            lattice_0.20-38
## [58] purrr_0.3.3             labeling_0.3             GenomicAlignments_1.18.1
## [61] htmlwidgets_1.5.1       bit_1.1-14               tidyselect_0.2.5
## [64] GGally_1.4.0            magrittr_1.5             R6_2.4.1
## [67] Hmisc_4.3-0             DBI_1.1.0                pillar_1.4.3
## [70] foreign_0.8-74          withr_2.1.2              survival_3.1-8
## [73] RCurl_1.95-4.12         nnet_7.3-12              tibble_2.1.3
## [76] crayon_1.3.4            rmarkdown_2.0            progress_1.2.2
## [79] data.table_1.12.8       blob_1.2.0               digest_0.6.23
## [82] munsell_0.5.0
```
